# Intellectual disability risk gene RFX4 regulates cortical neurogenesis by restraining neuronal differentiation

**DOI:** 10.64898/2026.02.11.705165

**Authors:** Julianna J. Determan, Gareth Chapman, Faiza Batool, Sydney R. Crump, Jemma C. Gaines, Komal Kaushik, Sofia Malik, Monica Sentmanat, Xiaoxia Cui, Haley Jetter, Kristen L. Kroll

## Abstract

Despite the recent identification of RFX4 as a neurodevelopmental disorder risk gene, its role in cortical development remained unclear. Here, we derived human stem cell models of deficiency and pathogenic mutation and demonstrated that RFX4 restrains neurogenesis by repressing pro-neuronal gene expression, while maintaining lineage-specific roles in regulating proliferation during cortical inhibitory neuron development. We also defined requirements for RFX4 in recruiting RFX3 to chromatin to regulate synaptic and activity-dependent gene expression in progenitors and neurons, respectively. RFX4 deficiency persistently dysregulated neuronal differentiation in cortical organoids, while not affecting neurons generated by direct differentiation. Finally, we modeled pathogenic mutation of the RFX4 DNA-binding domain, finding that DNA binding was strongly reduced but gene expression was dysregulated differently from our deficiency models, supporting a mechanism of pathogenicity distinct from haploinsufficiency. Together, this work identified requirements for RFX4 during cortical development, building a necessary foundation for elucidating the etiology of RFX4-associated disorders.

## Introduction

Development of the human cerebral cortex is a highly regulated process, with disruptions contributing to the etiology of neurodevelopmental disorders (NDDs) including intellectual disability (ID) and autism spectrum disorder (ASD) (Albert et al., 2017; Cadwell et al., 2019; Lui et al., 2011). A central hypothesis for the neuronal dysfunction underlying these NDDs involves imbalanced excitatory and inhibitory neuronal activity in the cerebral cortex (E/I imbalance) (Hollestein et al., 2023; Siegel-Ramsay et al., 2021) resulting from altered development and/or function of glutamatergic excitatory neurons (cEXs) and/or GABAergic inhibitory neurons (cINs). A major contributor to E/I imbalance is changes in the highly regulated process of cortical neurogenesis, the process by which proliferative neural progenitors differentiate into post-mitotic cortical neurons. While transcription factors (TFs) are major regulators of cortical neurogenesis (Chapman et al., 2024; Cholfin & Rubenstein, 2007; Vasan et al., 2025; Villalba et al., 2021), with pathogenic mutations in these identified as frequent NDD contributors, it remains unclear how NDD-associated TF dysfunction differentially affects cEX versus cIN development to cause NDDs in humans. Thus, identifying both cell type-specific and –agnostic roles for NDD-associated TFs in regulating cortical neurogenesis is critical to understanding NDD etiology.

Towards this end, we modeled epigenetic regulation of human cIN development in prior work, deriving an atlas of histone modification state changes accompanying cIN specification and differentiation from human pluripotent stem cells (hPSCs) (Chapman et al., 2024). This work identified Regulatory Factor X (RFX) 4 and RFX3 as key TFs regulating this process. As both *RFX3* and *RFX4* were also recently identified as NDD risk genes, with pathogenic mutations associated with global developmental delay, epilepsy, and ASD (Harris et al., 2021), these observations merited further study. Furthermore, while RFX3 is broadly expressed during human cortical development and included in multi-gene deletions that contribute to 9p deletion syndrome (Ajami et al., 2023; Banerjee et al., 2019; Wang et al., 2025), RFX4 expression is largely restricted to neural progenitors (Wang et al., 2025), suggesting a distinct role for RFX4 in regulating cortical neurogenesis.

RFX4 was initially identified as a gene involved in ciliogenesis (Ashique et al., 2009), while work in murine Rfx4 knockout (KO) models demonstrated Rfx4’s necessity for proper brain development, with homozygous (HOM) KO causing lethality by embryonic day 18.5 (Ashique et al., 2009; Blackshear et al., 2003; Xu et al., 2018). Rfx4 heterozygous (HET) knockdown animal models exhibited altered dorsal-ventral brain patterning associated with dysregulation of WNT, BMP, and SHH signaling (Ashique et al., 2009; Sedykh et al., 2018; Zhang et al., 2006; Zhang et al., 2007), suggesting that RFX4 may regulate lineage specification. More recently, studies using hPSC models have provided conflicting evidence for the role of RFX4 in cortical neurogenesis, suggesting that RFX4 overexpression could either produce a more homogenous neuronal progenitor population (Joung et al., 2023) or instead induce neuronal differentiation (Choi et al., 2024). Therefore, identifying the role of RFX4 during human cortical neurogenesis across cEX and cIN development is a necessary precursor to defining how pathogenic RFX4 mutations may alter cortical neurogenesis to disrupt E/I balance and contribute to NDDs.

Here, we used hPSC models with RFX4 loss of function (LOF) mutation to demonstrate that RFX4 LOF triggers dose-dependent precocious neurogenesis during both cIN and cEX development, ultimately resulting in deficits in cortical neuron differentiation in organoids. Supporting previous animal model studies, we found that RFX4 LOF also altered dorsal-ventral patterning in neural progenitors. We defined the pattern of endogenous RFX4 genome-wide binding, demonstrating that RFX4 commonly represses the expression of pro-neuronal and synaptic genes during both cEX and cIN development, while also activating proliferative gene expression, specifically in medial ganglionic eminence-like progenitors during cIN development. We further demonstrated that regulation of cortical neurogenesis by RFX4 involves functional interactions with RFX3, which mediates activation of neuronal gene expression, while RFX4 LOF alters the capacity for RFX3 to bind the genome in both neural progenitors and cEXs. Finally, we modeled a patient-specific missense mutation in the RFX4 DNA binding domain, demonstrating that this causes a dramatic loss of RFX4 genome binding, but results in transcriptomic dysregulation and cellular phenotypes distinct from LOF. Together, this defines RFX4 requirements for cortical neurogenesis, elucidates the underlying mechanisms including a transcriptional regulatory relationship with RFX3, and demonstrates that the etiology of RFX4-associated NDDs is more complex than haploinsufficiency-related LOF, as previously hypothesized. These findings provide a foundation for understanding transcriptional control of cortical neurogenesis, elucidating the basis of *RFX4* variant pathogenicity, and targeting these mechanisms for reversal to ameliorate NDD-related consequences.

## Results

### Loss of RFX4 causes dosage-dependent induction of cortical neurogenesis

To characterize the effects of RFX4 loss of function (LOF) on human cortical and subcortical (hereafter denoted cortical) development, we developed human pluripotent stem cell (hPSC; AN1.1 and WA09 cell background) models with heterozygous (HET) or homozygous (HOM) RFX4 gene disruption (Figure 1A, S1A). Using our established protocols (Chapman et al., 2022; Chapman et al., 2024; Meganathan et al., 2017), we specified these hPSCs into neuronal progenitors (NPCs) with a dorsal (D-) or ventral (V-) telencephalic character (Figure 1B), validating a ∼60% and >90% loss of both RFX4 mRNA and protein in the HET and HOM models, respectively (Figure S1B-C). We next assessed how RFX4 deficiency changed neurosphere outgrowth during D-NPC and V-NPC specification, observing a significant reduction of outgrowth from both V-NPC (Figure S2A-C) and D-NPC neurospheres (Figure S2D-F) at day 12 (D12) of differentiation, without impacting germ-layer specification (Data S1). Interestingly, while neurosphere size was reduced during V-NPC specification (Figure S2C), it was not significantly affected by RFX4 LOF during D-NPC specification (Figure S2F), suggesting lineage specific roles for RFX4.

**Figure 1.**
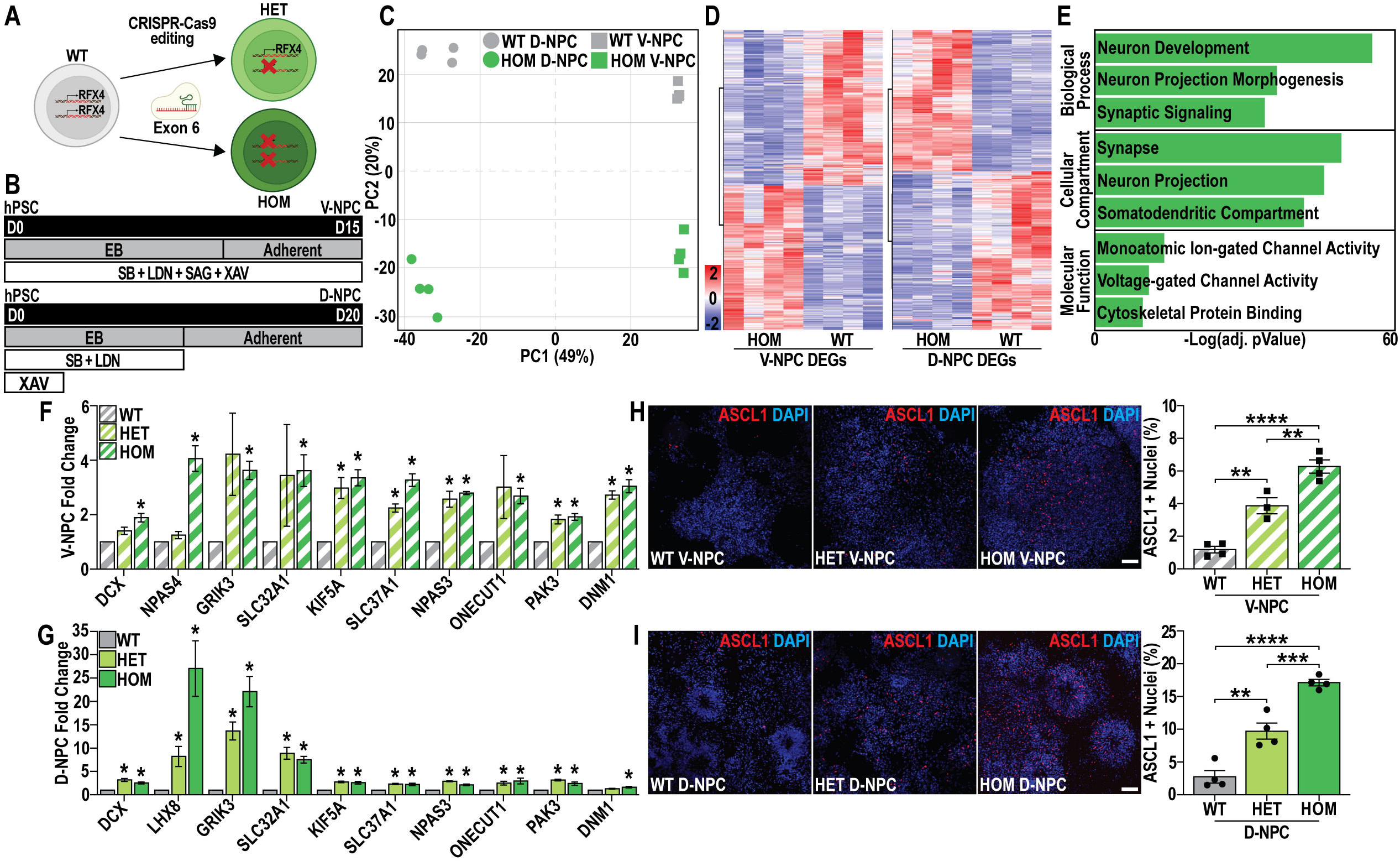
Loss of RFX4 causes dosage-dependent induction of cortical neurogenesis. (**A-B**) Schematic of **(A)** heterozygous (HET) and homozygous (HOM) RFX4 loss of function (LOF) hPSC model generation in WT hPSCs (WA09 and AN1.1) and **(B)** protocols used to specify hPSCs to ventral (V-) and dorsal (D-) neural progenitor cells (NPCs). **(C)** Principal component analysis (PCA), highlighting variation between WT D-NPCs (grey circles), RFX4 HOM D-NPCs (green circles), WT V-NPCs (grey squares), and RFX4 HOM V-NPCs (green squares). **(D)** Heatmaps of differentially expressed genes (DEGs) between WT and HOM V-NPCs (left) or D-NPCs (right). **(E)** Summary of gene ontology (GO) enrichment analysis for DEGs upregulated by RFX4 LOF in both D-NPCs and V-NPCs. **(F-G)** RT-qPCR assessed altered expression of genes significantly upregulated upon RFX4 LOF, showing relative expression in WT, HET and HOM **(F)** V-NPCs and **(G)** D-NPCs. **(H-I)** Representative images and quantification of ASCL1+ **(H)** V-NPCs or **(I)** D-NPCs. Data are represented as **(F-I)** mean ± SEM, with **(F-G)** fold change calculated relative to GAPDH, and **(E)** as the –Logfold change of the Bonferroni adjusted pValue. Data in **(F-G)** was analyzed using Student’s t-test (n=4) or **(H-I)** one-way ANOVA (n=4); ns = not significant, \**p*<0.05, \*\**p*<0.01, \*\*\**p*<0.001, and \*\*\*\**p*<0.0001, **(H-I)** scale bars = 50 μm.

To further investigate lineage-specific RFX4 requirements, we performed RNA-sequencing in WT versus HOM D-NPCs and V-NPCs, finding shared transcriptomic changes by RFX4 LOF in both D-NPCs and V-NPCs (Figure 1C). We identified similar numbers of differentially expressed genes (DEGs) between WT and HOM D-NPCs (2,718) and HOM V-NPCs (2,456), respectively, with ∼50% of DEGs upregulated upon RFX4 LOF (Figure 1D, Data S2). Notably, DEGs downregulated in both HOM D-NPCs and V-NPCs were enriched predominantly for genes involved in early developmental processes (Data S3), while genes jointly upregulated upon RFX4 LOF were instead enriched for ‘synaptic signaling’ genes (Figure 1E, Data S3). Together, these data suggest that RFX4 may regulate neuronal gene expression similarly in both D-NPCs and V-NPCs.

We next validated altered expression of 10 upregulated (Figure 1F-G) and 10 downregulated (Figure S2G-H) DEGs, using RT-qPCR in WT, HET, and HOM D– and V-NPCs and finding that RFX4 LOF caused dosage-dependent increase in neuronal and synaptic gene expression (Figure 1F-G). Using immunocytochemistry to examine the fraction of cells expressing ASCL1, a neurogenesis-promoting bHLH transcription factor, we found that the loss of RFX4 caused a dosage-dependent increase in the ASCL1 immunopositive (+) cells in both V-NPCs and D-NPCs (Figure 1H-I). The area of DCX+, a protein essential for neuronal migration, and TUBB3+, an early neuronal marker, cells relative to total nuclei were likewise increased in both HET and HOM D-NPCs (Figure S3A-B), while HET and HOM V-NPCs only showed consistent increases in TUBB3 (Figure S3C-D). Furthermore, while V-NPCs exhibited decreased proliferation, proliferation remained unchanged in D-NPCs (Figure S3E-F), supporting lineage-specific requirements for RFX4. Similar to findings in Rfx4 knockout animal models (Ashique et al., 2009; Sedykh et al., 2018), the loss of RFX4 also altered lineage specification to dorsal versus ventral lineages, with RFX4 LOF D-NPCs now expressing the medial ganglionic eminence-specific marker NKX2.1 (Figure S3G-H). Upon assessing the dorsal telencephalic marker PAX6 in V-NPCs, we found that neither the WT nor the RFX4 LOF V-NPCs exhibited greater than 1% of PAX6+ cells (Figure S3I-J), supporting a lack of dorsalization resulting from RFX4 LOF in V-NPCs. Ultimately, these results support prior work showing altered dorsal-ventral patterning upon RFX4 LOF, while demonstrating that RFX4 LOF also triggers dosage-dependent neuronal differentiation.

Interestingly, these results appeared inconsistent with a previous publication (Choi et al., 2024), which concluded that RFX4 overexpression (OE) induced neuronal differentiation in hPSCs. Therefore, we tested whether RFX4 OE alone was sufficient to induce neuronal differentiation (Figure S4A). After confirming RFX4 OE (Figure S4B), we assessed expression levels of multiple neuronal genes by RT-qPCR, finding no significant increases (Figure S4C). We found that RFX4 OE was insufficient to induce the expression of TUBB3 or ASCL1 (Figure S4D-E) nor change the proportion of cells immunopositive for pluripotency markers (Figure S4F-G). Furthermore, using this same construct in D-NPCs (Figure S4H-I), we found that the overexpression of RFX4 in D-NPCs did not change the proportion of cells immunopositive for PAX6, TUBB3, or DCX (Figure S4J-K). Together, these results further support our hypothesis that RFX4 is required to regulate neurogenesis by restraining premature neuronal differentiation, while RFX4 OE alone is insufficient to induce neurogenesis.

### RFX4 directly restrains the onset of neuronal and synaptic gene expression

To identify genes directly regulated by RFX4, we defined endogenous RFX4 genome-wide binding using CUT&Tag, confirming an almost complete loss of RFX4 genome-wide binding in HOM D-NPCs (Figure 2A). We analyzed the quality of these data by footprinting, identifying a footprint under RFX4 bound peaks in WT D-NPCs corresponding with a canonical RFX transcription factor binding site (TFBS) that was lost in HOM D-NPCs (Figure S5A). We further compared our endogenous RFX4 binding data to previously published chromatin immunoprecipitation (ChIP) data obtained by overexpressing epitope-tagged RFX4 (Joung et al., 2023), finding over 12,000 binding sites specific to endogenous RFX4 (Figure S5B). Finally, we used a deep learning model to analyze the sequence syntax present at RFX4 bound peaks, demonstrating that the presence of an RFX TFBS was highly correlated with sites of RFX4 binding (Figure S5C). This work therefore identified the endogenous binding profile for RFX4 during human cortical development, confirming both the data validity and quality.

**Figure 2.**
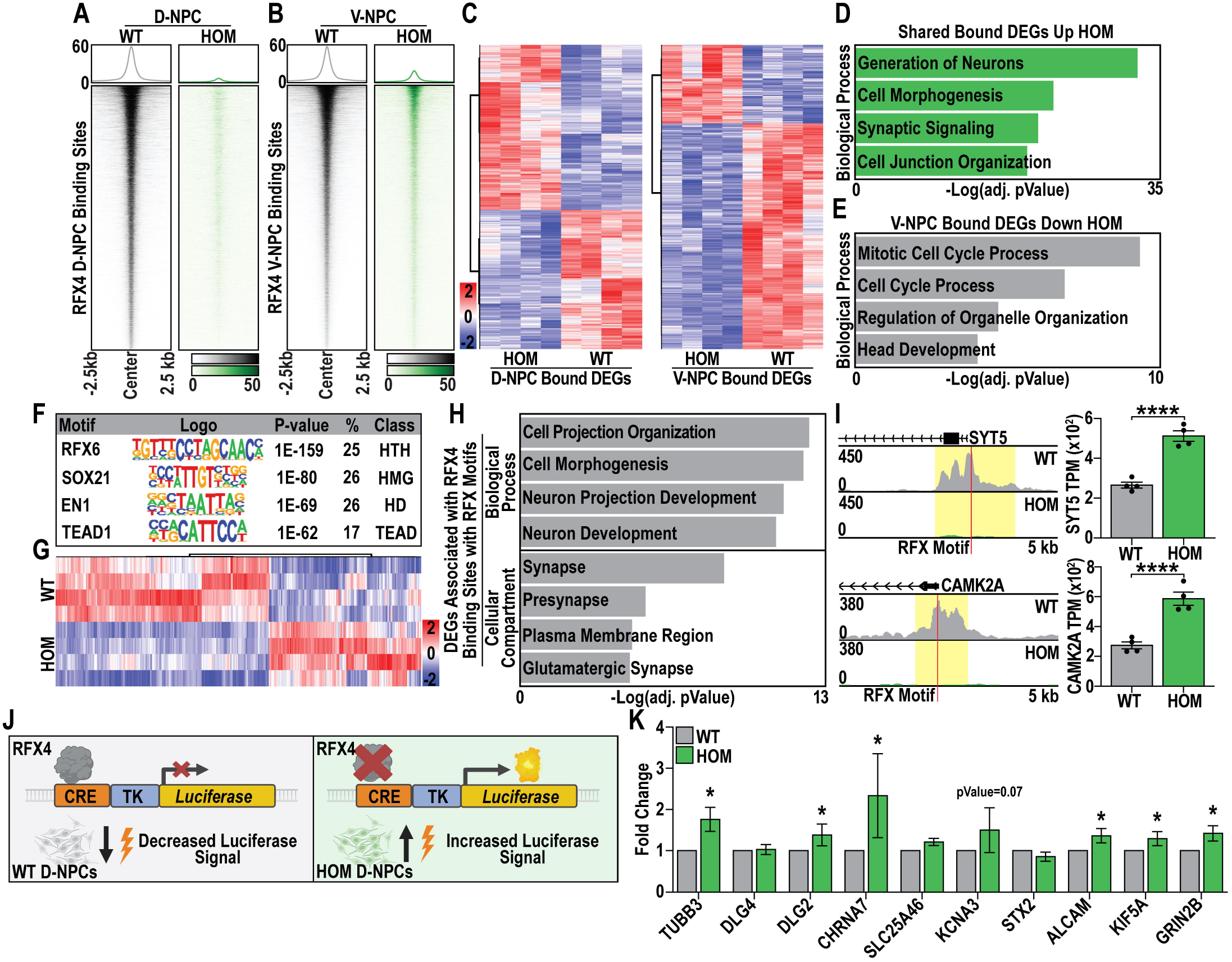
RFX4 directly restrains the onset of neuronal and synaptic gene expression. (**A-B**) Heatmaps of all RFX4 bound sites in WT and HOM **(A)** D-NPCs and **(B)** V-NPCs. **(C)** Heatmaps of DEGs associated with RFX4 binding (bound DEGs) in D-NPCs (left) and V-NPCs (right). **(D-E)** Summary of GO enrichment analysis for bound DEGs **(D)** upregulated in HOM V-and D-NPCs or **(E)** downregulated in HOM V-NPCs. **(F)** Summary of transcription factor binding site (TFBS) enrichment analysis of RFX4 bound DEGs. **(G)** Heatmap of RFX4 D-NPC bound DEGs associated with an RFX TFBS. **(H)** Summary of GO enrichment analysis for RFX4 bound DEGs associated with an RFX TFBS. **(I)** Examples of RFX4 D-NPC bound DEGs containing an RFX TFBS. **(J)** Experimental scheme and model for testing the hypothesized regulatory relationship between RFX4 and neuronal gene expression using luciferase assays. **(K)** Relative fold change in luciferase signal in HOM D-NPCs relative to the WT control. Data is represented as **(I,K)** mean ± SEM, **(D,E,H)** the –Logfold change of the Bonferroni adjusted pValue and was analyzed by **(I)** differential gene expression analysis (n=4), or **(K)** Student’s t-test (n≥3); **(I,K)** ns = not significant, \**p*<0.05 and \*\*\*\**p*<0.0001.

We next generated similar genome-wide binding data for RFX4 in WT versus HOM V-NPCs (Figure 2B), identifying both D-NPC and V-NPC shared and lineage-specific RFX4 binding sites (Figure S5D). To define genes directly regulated by RFX4, we associated RFX4 binding sites with their nearest transcription start site (TSS) and intersected these genes with our DEGs defined in D-NPCs and V-NPCs, finding substantial RFX4 binding near promoters and introns of bound-DEGs (Figure S5E-J). Bound-DEGs upregulated upon RFX4 LOF in both D-NPCs and V-NPCs were enriched for neuronal and synaptic genes (Figure 2C-D, Data S4), supporting a role for RFX4-mediated repression in regulating neurogenesis. By contrast, bound-DEGs downregulated upon RFX4 LOF uniquely in V-NPCs were associated with cell proliferation (Figure 2E, Data S4), consistent with our previous finding of reduced proliferation in HOM V-NPCs. These results suggest that while RFX4 has lineage-specific functions, it broadly represses neuronal and synaptic gene expression in both D-NPCs and V-NPCs.

As our work above highlighted shared cellular phenotypes across NPC lineages were more severe during D-NPC specification (Figure 1I, Figure S3A-B), we next focused on understanding RFX4’s regulatory function in D-NPCs. We found that binding sites that were significantly lost upon RFX4 LOF (>2 fold, pValue<0.05) and were associated with DEGs (Data S5) were enriched for both RFX motifs and for TEAD, homeodomain, and SOX family TFBSs (Figure 2F, Data S6), reminiscent of predictions made by our deep learning model (Data S5). To identify a subset of key RFX4 direct targets for further experimental validation, we next focused on RFX4 bound sites that were both associated with a DEG and contained an RFX4 TFBS (Figure 2G), finding that this DEG subset was enriched for synapse-associated genes upregulated upon RFX4 LOF (e.g. *SYT5*, *CAMK2A*; Figure 2H-I, Data S4). We cloned a set of RFX4 putative cis-regulatory elements (CREs) associated with these bound-DEGs into a minimal promoter-luciferase reporter construct (Figure 2J), finding that luciferase expression driven by a majority of these putative CREs increased significantly upon RFX4 LOF in D-NPCs (Figure 2K). These data demonstrated that RFX4 represses neuronal and synaptic gene expression at these sites, identifying new requirements for RFX4-mediated transcriptional repression of synaptic gene expression during cortical neurogenesis.

### RFX3 binding is contingent on RFX4 but plays a unique neurodevelopmental role

RFX family TFs can both homo– and hetero-dimerize (Morotomi-Yano et al., 2002) while RFX3 deletion is also associated with NDDs (Ajami et al., 2023; Banerjee et al., 2019); therefore, we sought to understand if RFX4 and RFX3 functionally interact to regulate the onset of neurogenesis. To assess this potential functional relationship, we first compared their expression, finding that both RFX3 and RFX4 expression peaked in NPCs, while only RFX3 expression persisted following neurogenesis (Figure 3A). We next examined how RFX4 LOF affected RFX3’s ability to bind DNA, finding a substantial loss of RFX3 genome-wide binding in RFX4 HOM D-NPCs (Figure 3B, Data S7), particularly at neuronal and synaptic gene CREs (Figure 3C, Figure S6A, Data S8), despite RFX3 expression remaining unchanged in HOM D-NPCs (Data S2, Figure S6B). Furthermore, we identified RFX3 and RFX4 co-bound sites (Figure 3D, Figure S6C) and associated these sites with RFX4 DEGs, finding an enrichment for RFX, TEAD, and SOX TFBSs (Figure 3E, Data S9), with the co-bound genes predominately involved in synapse development (Figure 3F, Data S8). Together, these results indicate that RFX3 binding is contingent upon RFX4 expression and suggest that both RFX TFs play roles in direct regulation of synapse-related gene expression.

**Figure 3.**
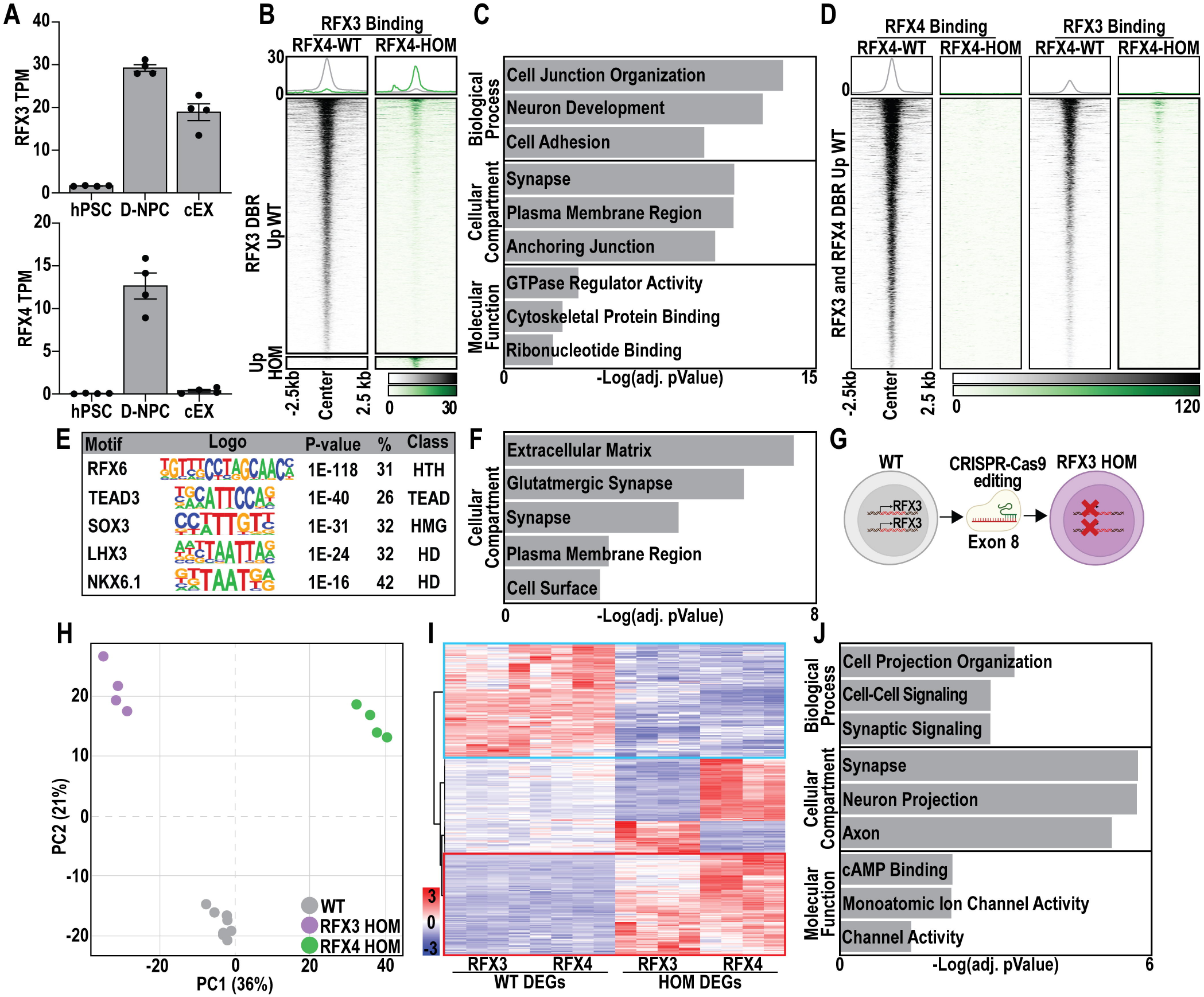
RFX3 binding is contingent on RFX4 but plays a unique neurodevelopmental role. **(A)** RFX3 (top) and RFX4 (bottom) gene expression during cortical excitatory (cEX) development highlighting that RFX3 continues to be expressed in cEXs, while RFX4 does not. **(B)** Heatmap of RFX3 binding in RFX4 WT and HOM D-NPCs, separating sites that lost (upper panel) or gained (lower) RFX3 binding in response to RFX4 LOF. **(C)** Summary of GO enrichment analysis for genes associated with an RFX3 bound site that was lost in RFX4 HOM D-NPCs. **(D)** Heatmaps of RFX3 and RFX4 binding for co-bound sites that lost RFX4 binding in RFX4 HOM D-NPCs. **(E)** Summary of TFBS enrichment analysis for RFX3/4 co-bound sites at which binding is lost in RFX4 HOM D-NPCs. **(F)** Summary of GO enrichment analysis of RFX4 DEGs that are associated with RFX3/4 co-bound sites lost in RFX4 HOM D-NPCs. **(G)** Schematic of homozygous (HOM) RFX3 loss of function (LOF) hPSC model generation. **(H)** PCA plot highlighting variation in gene expression between WT (grey circles), RFX3 HOM (purple circles), and RFX4 HOM (green circles) D-NPCs. **(I)** Heatmap of DEGs dysregulated upon both RFX3 LOF or RFX4 LOF, with genes similarly downregulated or upregulated in both RFX3 and RFX4 HOM models highlighted (blue and red boxes, respectively). **(J)** Summary of GO enrichment analysis for DEGs downregulated upon RFX3 LOF. Data represents **(A)** mean ± SEM **(C,F,J)** the –Logfold change of the Bonferroni adjusted pValue.

To assess if the RFX4-dependence of RFX3 binding we observed above was reciprocal, we next generated hPSC models with homozygous RFX3 LOF (RFX3-HOM, Figure 3G, Figure S6D) and validated loss of RFX3 expression in RFX3-HOM D-NPCs (Figure S6E-F). We assessed RFX3 genome-wide binding in WT and RFX3-HOM D-NPCs, demonstrating that RFX3 binding was enriched at gene promoters and was lost upon RFX3 LOF (Figure S6G-H, Data S7). However, RFX3 LOF resulted in minimal alteration of RFX4 binding (Figure S6I), indicating that RFX4 genome-binding does not reciprocally depend upon RFX3. Additionally, RFX3 LOF resulted in unique cellular phenotypes; for example, unlike RFX4 LOF, RFX3 LOF did not change neurosphere outgrowth nor the proliferative or neurogenic marker immunopositivity (Figure S6J-N), suggesting that RFX3, unlike RFX4, is not required to restrain cortical neurogenesis. Interestingly, transcriptomic analysis confirmed both distinct and shared gene expression changes upon either RFX3 LOF or RFX4 LOF (Figure 3H, Data S10). Specifically, LOF of each TF resulted in shared upregulation of apoptosis associated genes (Figure 3I, red box, Data S8) and shared downregulation of early neurodevelopment genes (Figure 3I, blue box, Data S8). However, RFX3 LOF uniquely caused the downregulation of synapse-related genes, many of which were bound by RFX3 at promoters under WT conditions (Figure 3J, Figure S6O-P, Data S8). Many of these genes were instead upregulated by RFX4 LOF (Data S8). These results highlight both shared and antagonistic roles for RFX3 and RFX4 during cortical neurogenesis, specifically in restraining synaptic gene expression, with RFX4 but not RFX3 having a substantial role in regulating this process.

### RFX4 expression during neurogenesis is necessary for neuronal layering in cortical organoids

To examine the neuronal consequences of altered cortical neurogenesis resulting from RFX4 LOF, we next generated cortical excitatory neurons (cEXs) by differentiating WT and HOM D-NPCs (Figure 4A) and glutamatergic-like neurons (iGluts, Figure 4B) by overexpressing Neurogenin-2 (NGN2), in WT and HOM hPSCs. As the generation of iGluts bypasses the formation of neuronal progenitors and RFX4 expression is temporally restricted to D-NPCs (Figure 4C, Figure S7A-B), we tested whether transcriptomic changes caused by RFX4 LOF specifically in D-NPCs were detected in cEXs but not iGluts. We examined neuronal genes directly regulated by RFX4 in D-NPCs, observing continued upregulation of these genes in HOM cEXs, while HOM iGluts exhibited no consistent changes in expression of these genes (Figure 4D). This assessment confirmed that neuronal consequences resulting from RFX4 LOF require transit through a prior progenitor stage. To further examine the neuronal consequences of LOF, we performed RNA-sequencing in WT and HOM cEXs (Figure S7C) identifying consistently dysregulated DEGs common to both HOM D-NPCs and HOM cEXs (Data S2, Data S11). This analysis identified a persistent upregulation of TFs involved in neuronal specification (e.g. ASCL1 Figure 4E-F, Data S12) in both HOM D-NPCs and cEXs, including ventral progenitor-specific markers. These data confirm that transcriptomic disruptions caused by RFX4 LOF originate during D-NPC specification and persist through cEX differentiation; these both ventralize D-NPCs and cause persistent elevation of pro-neurogenesis gene expression in neurons that could potentially impact their later differentiation or maturation.

**Figure 4.**
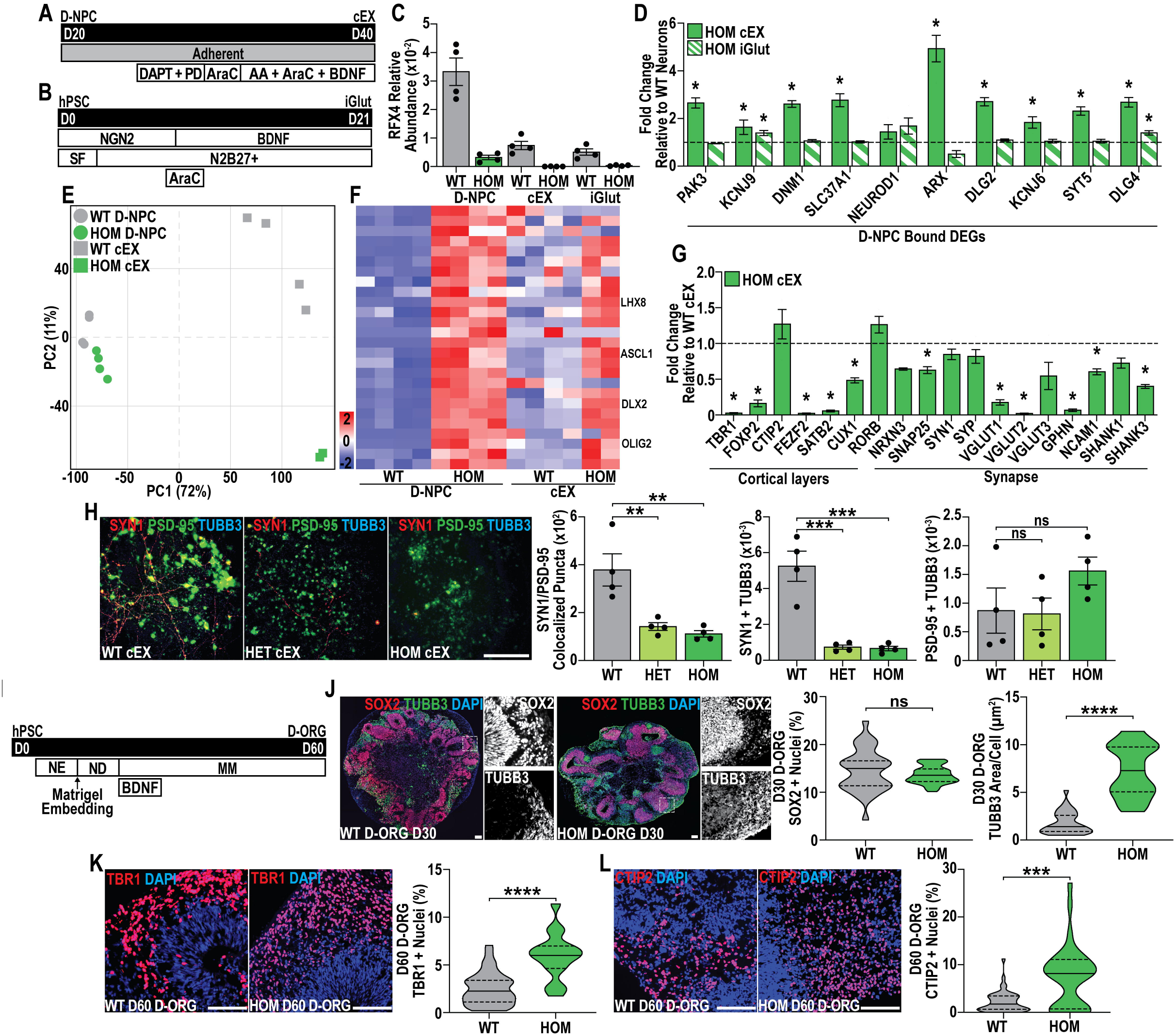
RFX4 expression during neurogenesis is necessary for neuronal layering in cortical organoids. (**A-B**) Differentiation schematic for generating **(A)** cortical excitatory neurons (cEXs) and **(B)** glutamatergic-like neurons (iGluts). **(C)** RT-qPCR of RFX4 mRNA expression in WT and HOM D-NPCs, cEXs, and iGluts relative to RPL30 endogenous control, highlighting restriction of RFX4 expression to D-NPCs. **(D)** RT-qPCR for a selection of RFX4 D-NPC bound DEGs upregulated in RFX4 HOM D-NPCs, showing fold change in expression across RFX4 HOM cEXs and iGluts relative to expression levels in the corresponding WT cells. **(E)** PCA plot highlights an increased divergence between the transcriptomic profiles of WT (grey) and HOM (green) cEXs (squares) versus D-NPCs (circles). **(F)** Heatmap of transcription factors upregulated upon RFX4 LOF in both D-NPCs and cEXs. **(G)** RT-qPCR for cortical layer and synaptic markers in HOM cEXs relative to WT cEXs. **(H)** Representative images and quantification of SYN1+ and PSD-95+ puncta either colocalized to each other or to TUBB3+ neurons in WT, RFX4-HET, and RFX4-HOM cEXs. **(I)** Schematic of the protocol used to generate dorsal cortical organoids (D-ORG). **(J-L)** Representative images and quantification of **(J)** SOX2+ and the area of TUBB3+ cells in D30 D-ORGs and **(K)** TBR1+ and **(L)** CTIP2+ in D60 D-ORGs. Data is represented as **(C,D,G,H)** mean ± SEM **(C,D,G)** with fold changes calculated relative to RPL30, or **(J-L)** as data distributions with quartiles (dashed lines) and median (solid line) marked. Data was analyzed by **(D)** Student’s t-test (n=4), **(H)** one-way ANOVA, or **(J-L)** Mann-Whitney test (for n values, see Data S20); ns = not significant, \**p*<0.05, \*\*\**p*<0.001, and \*\*\*\**p*<0.0001. Scale bars = **(H,J-L)** 100 μm.

To examine if these disruptions altered the later differentiation capacity, identity, and/or maturation of HOM cEXs, we next profiled the expression of cortical layer markers and synaptic genes critical for glutamatergic neuron function, finding that many of these genes were downregulated in HOM cEXs (Figure 4G), an interesting finding given that progenitors substantially upregulated the same classes of neuronal and synaptic genes. This finding, supported by our RNA-sequencing data (Data S12), suggests that undergoing precocious neurogenesis may impact the differentiation and/or maturation of neurons later in development. While we did not find a significant difference in total neuron production as assessed with the early neuronal marker TUBB3 (Figure S7D), we found the number of colocalized SYN1+ and PSD95+ puncta reduced by RFX4 LOF (Figure 4H) and a reduction of the glutamatergic puncta marker VGLUT1 (Figure S7E). Together, these data suggest that the premature initiation of neurogenesis caused by RFX4 LOF impacts some aspects of later neuronal differentiation.

The reduction of some neuronal marker expression in cEXs suggested that LOF effects on neurogenesis could persistently alter later neuronal differentiation and layering of the developing cortex. To examine this, we generated dorsal telencephalic patterned 3-D cortical organoids (D-ORGs, Figure 4I). We first assessed key neurogenesis-related phenotypes observed above in D-NPCs, focusing on an approximately analogous D-ORG timepoint, day 30 (D30). Like our prior findings for D-NPCs, HOM D-ORGs exhibited precocious neurogenesis, as marked by an increased area expressing the newborn neuron marker TUBB3, with no change to the fraction of SOX2-expressing progenitors (Figure 4J). This was congruent with the decreased KI67+ and increased TBR1+, the first-born neuronal subtype, cells (Figure S7F-G). We further detected an increase in FOXG1+, a cortical TF expressed throughout differentiation shown to be necessary for cIN development, highlighting deficits in neuronal regionalization by RFX4 LOF (Figure S7H). Finally, neither the intermediate progenitor marker TBR2 (Figure S7I) nor ASCL1 were affected in organoid models (Figure S7J). Together these results demonstrate that RFX4 restrains cortical neurogenesis, while LOF causes precocious neurogenesis in both 2-D and 3-D models of cortical development.

We further differentiated D-ORGs to day 60 (D60) to examine if the differentiation into different neuronal types expressed by different cortical layers is similarly altered upon RFX4 LOF. We found significant increases in the production of TBR1+ (Figure 4K) and CTIP2+ (Figure 4L) neurons upon RFX4 LOF and increases the TUBB3+ area relative to total nuclei in D60 D-ORGs, with no change in the apoptosis marker Cleaved Caspase 3 (CC3) (Figure S7K). These results demonstrate that the increased production of early-born neurons persists, altering later 3-D organoid gene expression associated with cortical neuron layering. Therefore, these results support a progenitor-specific requirement for RFX4 to restrain cortical neurogenesis and differentiation, with its disruption causing persistent effects in neuron differentiation and layering in these hPSC-based models of human cortical development.

### RFX3 function is altered in RFX4-deficient neurons

Our above data identified a set of RFX3 and RFX4 co-bound sites where RFX3 binding was dependent upon RFX4 expression, specifically at synaptic genes. We also found that RFX4 LOF altered gene expression in neurons, despite restriction of RFX4’s temporal expression to progenitors. Furthermore, RFX3 was previously shown to activate synaptic gene expression, specifically by interacting with CREB and in an activity-dependent manner (Lai et al., 2025; Neumann et al., 2025). Lai et al. also demonstrated the loss of RFX3 increased deep-layer neuron production in organoid culture, reminiscent of our findings in RFX4 LOF D-ORGs. Therefore, we sought to determine whether RFX3’s activity and function in neurons is altered or changed by RFX4 deficiency. To begin, we identified 10,319 endogenous RFX3 binding sites in cEXs (Figure S8A), 2,376 of which overlapped with previously defined RFX3 genome-wide binding in iGluts (Figure S8B) (Lai et al., 2025). We then found RFX3 binding significantly lost in RFX3-HOM cEXs (>2 fold, pValue<0.05) was associated with genes involved with cilium and channel activity (Figure 5A-B, Data S13), similar to prior findings (Lai et al., 2025; Neumann et al., 2025). We next examined if RFX3 genome-wide binding was changed in RFX4-HOM cEXs, finding a significant loss of RFX3 binding in RFX4-deficient neurons, which was associated with neuronal and synaptic genes (Figure 5C-D, Data S13-14) despite the continued expression of RFX3 in RFX4-HOM cEXs (Figure S8C).

**Figure 5.**
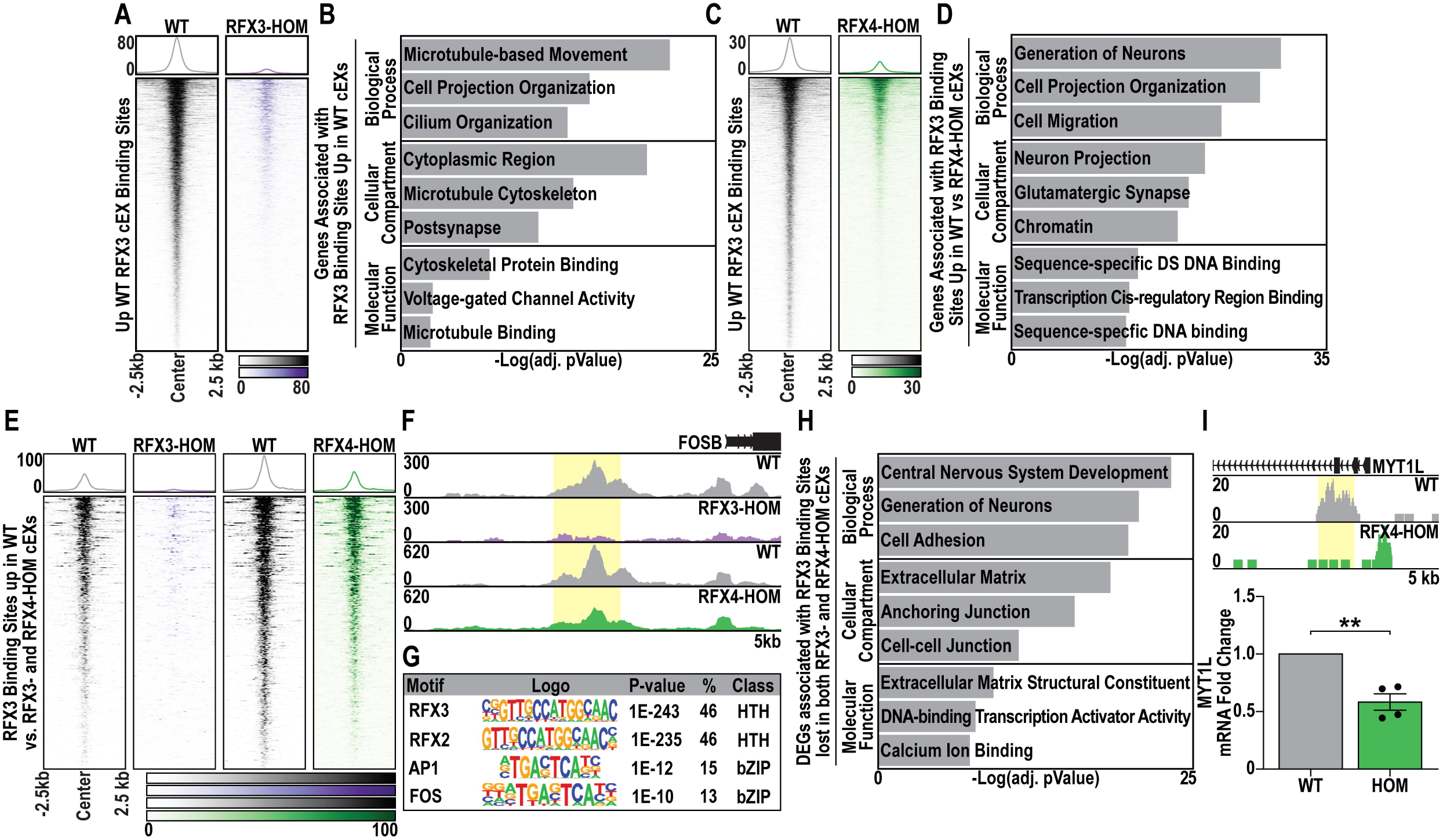
RFX3 function is altered in RFX4-deficient neurons. **(A)** Heatmap of RFX3 binding sites significantly lost in RFX3-HOM cEXs in WT versus RFX3-HOM cEXs. **(B)** GO analysis of genes associated with RFX3 binding sites that were significantly lost in RFX3-HOM cEXs. **(C)** Heatmap of RFX3 binding sites that were significantly lost by RFX4 LOF in WT and RFX4-HOM cEXs. **(D)** GO analysis of genes associated with RFX3 binding sites significantly lost in RFX4-HOM cEXs. **(E)** Heatmap of RFX3 binding sites lost in both RFX3-and RFX4-HOM cEXs, comparing WT, RFX3-HOM, and RFX4-HOM cEXs. **(F)** Example browser track of FOSB, highlighting a loss of RFX3 binding in both RFX3-and RFX4-HOM cEXs. **(G)** Summary of TFBS enrichment analysis for RFX3 binding sites significantly lost upon both RFX3 LOF and RFX4 LOF. **(H)** GO analysis of cEX DEGs downregulated upon RFX4 LOF and associated with significant loss of RFX3 binding. **(I)** Example browser track of MYT1L and gene expression changes, defined by RT-qPCR, highlight a loss of RFX3 binding associated with downregulation of MYT1L. Data is represented as **(I)** fold change relative to endogenous RPL30 and **(B,D,H)** the –Logfold change of the Bonferroni adjusted pValue and analyzed by **(I)** Student’s t-test (n=4). \*\**p*<0.01.

Despite a lack of appreciable RFX4 expression in neurons, we found that a subset of RFX3 binding sites were significantly lost upon either RFX3 LOF or RFX4 LOF (Figure 5E). These sites were associated with genes involved in synaptic plasticity (Data S13), suggesting that the loss of RFX4 during neurogenesis impairs the later capacity of RFX3 to regulate synaptic gene expression. For example, FOSB, an activity-dependent gene shown to respond to seizure-like activity, is reduced in RFX4-HOM cEXs, much like RFX3-HOM cEXs (Figure 5F). Furthermore, motif analysis of RFX3 binding sites lost in both RFX3– and RFX4-HOM cEXs shows an enrichment for the expected RFX motifs as well as the TFBSs recognized by the bZIP class transcription factors, AP1 and FOS (Figure 5G, Data S15), both of which are transcription factors that regulate gene expression in response to neuronal activity. These data suggest that RFX3 is unable to bind and regulate these activity-dependent genes in RFX4-deficient cEXs.

To understand if the altered RFX3 genome-wide binding could account for gene expression changes observed in RFX4 LOF cEXs, we associated genes that significantly lost RFX3 binding upon RFX4 LOF with RFX4 HOM cEX DEGs (871), most of which are downregulated in RFX4-HOM cEXs (769). These downregulated DEGs were associated with genes involved in the generation of neurons and cell-cell junctions (e.g. the ASD risk gene *MYT1L*, Figure 5H-I, Data S13). Taken together, these data suggest that RFX3 genome-wide binding and neuronal gene regulation is altered by RFX4 LOF in cEXs, despite the restriction of RFX4 expression to NPCs. Ultimately, this loss of RFX3 activity results in downregulation of genes necessary for neuronal maturation and responsive to neuronal activity, similar to prior work on RFX3-HOM neurons.

### Pathogenic *RFX4* mutation disrupts genome-wide binding, but has distinct effects from LOF on transcriptional regulation

As RFX4 was recently classified as an NDD risk gene with pathogenic mutations hypothesized to act by LOF (Harris et al., 2021), we next generated hPSC models heterozygous or homozygous for a pathogenic p.R79C missense mutation in the RFX4 DNA binding domain (R79C-HET and R79C-HOM, respectively; Figure 6A, Figure S9A). RFX4 mRNA and protein levels were assessed in R79C-HET and R79C-HOM D-NPCs, highlighting reduced RFX4 protein levels (∼50% decrease) in the R79C-HOM but not R79C-HET D-NPCs (Figure S9B-C); these data suggest that this mutation could disrupt the expression and/or protein stability of RFX4 and may alter its function. Consistent with pathogenic disruption of the DNA binding domain of RFX4, we found an almost complete loss of RFX4 genome-wide binding in R79C-HOM D-NPCs (Figure 6B, Figure S9D, Data S16). This involved loss of binding at many of the same locations and associated genes defined in the LOF HOM D-NPCs (e.g. *DLG4* and *SCRN1*; Figure 6C-E, Data S17), supporting LOF as a potential mechanism of R79C pathogenicity.

**Figure 6.**
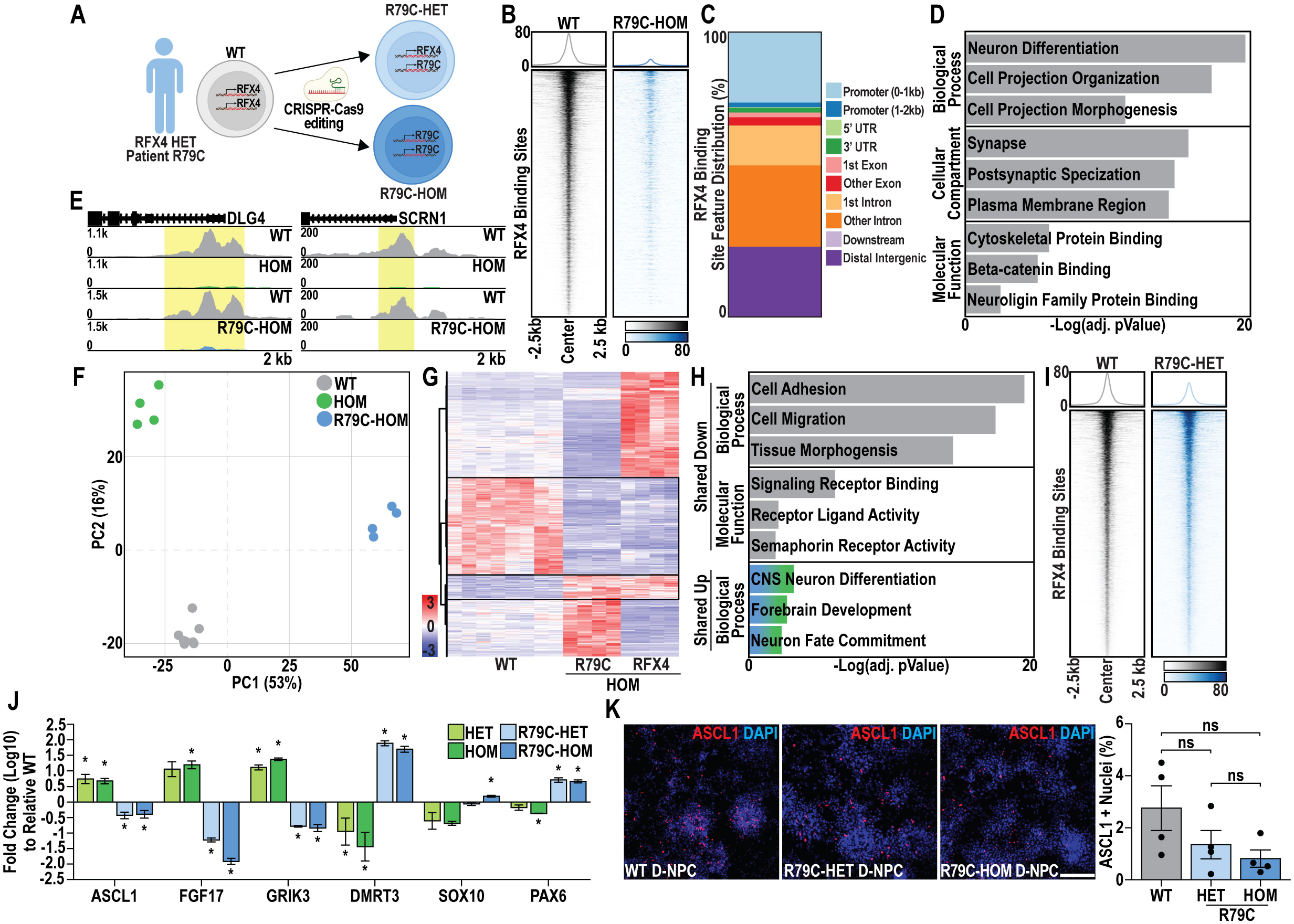
Pathogenic *RFX4* mutation disrupts genome-wide binding, but has distinct effects from LOF on transcriptional regulation. **(A)** Variant knock-in of a heterozygous (R79C-HET) or homozygous (R79C-HOM) RFX4 missense mutation into WT hPSCs to model RFX4 pathogenic mutation. **(B)** Heatmap of RFX4 binding in WT and R79C-HOM D-NPCs. **(C)** Distribution of RFX4 binding sites relative to key genome features. **(D)** Summary of GO enrichment analysis for genes associated with RFX4 binding sites that were lost in R79C-HOM D-NPCs. **(E)** Examples of RFX4 binding in browser track views, highlighting similar losses in RFX4 HOM and R79C-HOM D-NPCs at two genes. **(F)** PCA plots highlighting distinct alterations in gene expression between the WT control (grey circles), RFX4 HOM (green circles), and R79C-HOM (blue circles) D-NPCs. **(G)** Heatmap of DEGs dysregulated upon RFX4 LOF and pathogenic mutation, highlighting genes similarly dysregulated by both perturbations (black boxes). **(H)** Summary of GO enrichment analysis for DEGs downregulated (top, grey) or upregulated (bottom, blue/green) in both R79C-HOM and RFX4 HOM D-NPCs. **(I)** Heatmap of RFX4 bound sites associated in R79C-HET D-NPCs. **(J)** RT-qPCR for a selection of R79C-HOM and RFX4 HOM DEGs that showed opposing gene expression changes across the LOF and pathogenic mutant HET and HOM models. **(K)** Representative images and quantification of ASCL1+ cells in WT, R79C-HET and –HOM D-NPCs. Data are represented as **(J,K)** mean ± SEM with fold changes calculated relative to RPL30 and **(D,H)** the –Logfold change of the Bonferroni adjusted pValue. Data was analyzed by **(J)** Student’s t-test (n=4) or **(K)** one-way ANOVA (n=4); ns = not significant and \**p*<0.05. **(K)** Scale bar = 50 μm.

We also assessed transcriptomic changes in R79C-HOM versus WT D-NPCs (Data S18), surprisingly finding substantially different DEGs in R79C-HOM versus LOF HOM D-NPCs, in DEG comparisons with their matched controls (Figure 6F). Likewise, unlike the LOF model, R79C-HOM D-NPCs did not alter neurosphere outgrowth (Figure S9E-G). Further analysis identified many R79C-HOM specific DEGs (Data S17), with downregulation of neuronal and synaptic gene expression prominent, by contrast with our findings for HOM D-NPCs (Data S2, Data S17). We also identified a small set of shared DEGs with similarly dysregulated expression in both R79C-HOM and HOM D-NPCs (Figure 6G); these shared, downregulated DEGs were enriched for cell migration genes, while shared upregulated DEGs were instead enriched for genes involved in early forebrain development (Figure 6H, Data S17). Interestingly, the profile of RFX3 genome-wide binding was not disrupted extensively in the R79C-HOM D-NPCs (Figure S9H, Data S16) as seen in the RFX4-HOM D-NPCs (Figure 3B), potentially due to maintenance of RFX4 protein carrying the R79C mutation (Figure S9C). Together, these data demonstrate that, while p.R79C pathogenic mutation disrupts genome-wide RFX4 binding like LOF, it does not impact RFX3 binding as severely as LOF and causes distinct transcriptomic changes.

We next tested whether these transcriptional consequences were dependent upon homozygosity for the p.R79C mutation, finding that, unlike R79C-HOM, D-NPCs heterozygous for this mutation (R79C-HET) exhibited minimal changes in RFX4 genome-wide binding in D-NPCs (Figure 6I, Figure S9I, Data S16). We also examined a selection of DEGs that were differentially regulated in the R79C versus the LOF models, finding that both R79C-HET and R79C-HOM D-NPCs exhibited similar gene expression changes, while these differed from those seen in the HET and HOM LOF models (Figure 6J). Furthermore, while LOF D-NPC models exhibited an increase in ASCL1+ (Figure 1I), neither R79C-HET or R79C-HOM D-NPCs did (Figure 6K). Likewise, the KI67+, FOXG1+, and PAX6+ cell fractions were unchanged upon p.R79C mutation (Figure S9J-L). These findings demonstrate that, despite similarly reducing genome-wide RFX4 binding, the p.R79C mutation disrupts transcriptional regulation distinctly from haploinsufficiency-related LOF.

## Discussion

In this work, we derived new hPSC models and used these to identify requirements for RFX4 and consequences of pathogenic mutation during human cortical development. We demonstrated that RFX4 restrains cortical neurogenesis in both 2-D and 3-D models and during both cortical excitatory (cEX) and inhibitory neuron (cIN) differentiation. While RFX transcription factors were previously assumed to behave as transcriptional activators (Choi et al., 2024; Harris et al., 2021), this work demonstrates a requirement for RFX4 to directly repress neuronal and synaptic gene expression during this process. We further found that both molecular and cellular phenotypes associated with RFX4 LOF in progenitors persist through neuronal differentiation, despite the cessation of RFX4 expression. We tested the consequences of RFX4 overexpression, as prior studies suggested that RFX4 could both stabilize neuronal progenitor production (Joung et al., 2023) and induce neuronal differentiation (Choi et al., 2024), finding that RFX4 overexpression alone was insufficient to induce neuronal differentiation without additional factors known to induce neurons (e.g. NEUROD1). Our work here in LOF models demonstrates that RFX4 is required to repress neuronal gene expression to maintain the neuronal progenitor state, while LOF induces neurogenesis.

To investigate the mechanisms by which RFX4 regulates cortical neurogenesis, we identified endogenous genome-wide RFX4 binding during both cEX and cIN development, confirming the validity and quality of these data across multiple paradigms. We demonstrate that RFX3 colocalizes on the genome with RFX4, co-occupying CREs associated with synaptic genes and functionally interacting to regulate gene expression, with RFX3 relying on the presence of RFX4 to bind most of these shared enhancers.

Despite co-occupying putative CREs at synaptic genes, RFX3 and RFX4 requirements for cortical neuron development differ; RFX4 expression specifically during neurogenesis is required to represses neuronal gene expression, while RFX3 can activate later neuronal and synaptic gene expression during differentiation, a finding substantiated by Lai and colleagues using RFX3 LOF iGlut and organoid models (Lai et al., 2025). RFX3 was previously shown to promote activity-dependent gene expression in cortical neurons through interactions with CREB1 (Lai et al., 2025; Neumann et al., 2025), versus the distinct role we describe here for RFX4 specifically in neural progenitors, suggesting distinct temporal and transcriptional activities for these TFs in cortical development. These divergent roles are congruent with the temporal restriction of RFX4 expression to progenitors, while RFX3 continues to be expressed in neurons (Wang et al., 2025). We additionally show that RFX3 genome-wide binding is altered in RFX4 LOF cEXs, suggesting that RFX4 expression in progenitors is required for RFX3 binding in cEXs. While our work here supports co-occupancy of RFX4 with RFX3 at enhancers during cortical neurogenesis, future work is needed to further define the mechanistic relationships between the activities of RFX4 and other TFs with enriched binding motifs under the RFX4 bound peaks we defined here.

Examining the persisting neuronal consequences of altered neurogenesis resulting from RFX4 LOF, we found that these were specific to neurons that underwent a progenitor stage rather than being directly differentiated into iGluts. Similar findings have been reported for other NDD hPSC models (Schafer et al., 2019), supporting a developmental etiology involving disrupted neurogenesis for NDDs resulting from RFX4 mutation. Studies exploring other NDD-associated mutations have also reported altered cortical neurogenesis (Adam & Harwell, 2020; Chapman et al., 2025; Kaushik & Zarbalis, 2016; Kaushik et al., 2025; Niu et al., 2024; Prakasam et al., 2025) that can cause later E/I functional neuron imbalance, a central hypothesis for the cellular etiology of NDDs (Hollestein et al., 2023; Siegel-Ramsay et al., 2021). For example, EZH1 LOF disrupts neurogenesis (Gracia-Diaz et al., 2023) while conditional Ezh2 knockout in mouse models enhances formation of TBR1+ and CTIP2+ neurons during peak cortical neurogenesis (Pereira et al., 2010). EZH2 has also been shown to act as an epigenetic barrier during neuronal differentiation of hPSC models (Ciceri et al., 2024). As mutations in each of these genes can cause NDDs, disruption of neurogenesis represents a critical developmental window sensitive to perturbation to cause NDDs.

Our cEX and cortical organoid models both exhibited precocious neurogenesis with persistent neuronal consequences, with LOF altering expression of lineage-specific neuronal markers and synaptic puncta localization. Interestingly, LOF in D-NPCs persistently upregulated some TFs specific to cINs in cEXs, while the increase in FOXG1 immunopositivity in LOF organoid models may also indicate altered progenitor identity, as FOXG1 overexpression in organoids was previously reported to increase GABAergic neuron production (Mariani et al., 2015). Our observations of perturbed progenitor identity here are reminiscent of the altered dorsal-ventral brain patterning seen in Rfx4 knockout mouse and zebrafish models (Ashique et al., 2009; Blackshear et al., 2003; Sedykh et al., 2018; Zhang et al., 2006), suggesting that RFX4 mutations may affect both neurogenesis and regional neuronal patterning.

Finally, having established requirements for RFX4 during cortical development, we examined how the pathogenic mutation p.R79C, a missense mutation in the DNA-binding domain predicted to act via LOF, affected regulation of cortical neurogenesis. Surprisingly, while this mutation largely abrogated RFX4 genome-wide binding like LOF models, it caused unique and substantial transcriptional changes, suggesting a distinct mechanism of pathogenicity. As RFX TFs can both homo– and heterodimerize (Morotomi-Yano et al., 2002) and co-bind shared genomic targets, pathogenic mutation may alter these dynamics. Furthermore, it has been shown that two RFX family TFs can co-bind a single canonical X-box motif (Gajiwala et al., 2000), suggesting that RFX family TF heterodimerization may alter their transcriptional regulatory activity. Studying the basis of pathogenicity of additional mutations will be required to understand the etiology of RFX4-associated NDDs and underlying mechanisms. Together, this study elucidated previously unknown roles for and mechanisms of transcriptional repression by RFX4 during cortical neurogenesis, highlighting their disruption upon both LOF and pathogenic missense mutation and providing a necessary foundation to develop approaches for reversal of pathogenesis-associated phenotypes.

## Author Contributions

J.J.D. and K.L.K. designed the study. J.J.D. performed developmentally patterned differentiations and downstream cellular phenotyping for RFX4 LOF, RFX3 LOF, and R79C experiments. G.C. performed iGlut experiments. J.J.D. generated dorsally patterned organoids, G.C. cryo-sectioned organoids, and J.J.D. and G.C. performed immunocytochemistry on organoids. F.B. performed and analyzed western blots. S.C. and J.C.G. performed RFX4 overexpression experiments. K.K. performed puncta staining and analysis. S.M. performed RT-qPCR and organoid immunocytochemistry. F.B. and H.J. generated immunofluorescence images. J.J.D. and G.C. performed bioinformatic analyses. M.S. and X.C. generated and confirmed validity of RFX3 KO hPSC lines. J.J.D. and G.C. analyzed data and generated figures. J.J.D. and K.L.K. wrote the manuscript and all authors approved the manuscript.

## Supporting information

Supplemental Data Table Titles and Supplemental Figures

## Acknowledgments

This work was supported by NIH Fellowship F31MH141956 to J.J.D., NIH Grants R01NS114551, R01MH124808, R01HD110556, R01MH138528, and R01NS152315 to K.L.K., and by NIH P50HD103525 to Joseph Dougherty and Christina Gurnett (K.L.K. is project PI for the Human Cellular Models Unit of the Washington University Intellectual and Developmental Disabilities Research Center, with project effort funded by this grant). Foundations supporting this work included the Engelhart Family Foundation, Simons Foundation, M-CM Network, and pilot awards from the WU Hope Center, Center of Regenerative Medicine, and Institute for Clinical and Translational Sciences to K.L.K.

We would like to acknowledge the Washington University Genome Engineering & Stem Cell Center (GESC@MGI) for generation of the genetically engineered induced human pluripotent stem cell lines used in this study. We further acknowledge the Washington University Genome Technology Access Center (GTAC@MGI) for their work in sequencing RNA and Cut&Tag data. Finally, we acknowledge the Washington University Human Cells, Tissue, and Organoids core (hCTO) for their work in generating early cortical organoid data.

## STAR Methods

### Resource Availability

#### Lead Contact

Further information and requests for resources and reagents should be directed to and will be fulfilled by the lead contact, Kristen L. Kroll

#### Materials Availability

All unique/stable reagents generated in this study are available from the lead contact with a completed Materials Transfer Agreement.

#### Data and code availability

All raw data (RNA and CUT&Tag) is available from the Gene Expression Omnibus database under accession GSE316377 and are publicly available as of the date of publication. This paper does not report original code. Any additional information required to reanalyze the data reported in this paper is available from the lead contact upon request.

### Experimental Model Details

#### Cell Lines

Work with human pluripotent stem cells (hPSCs) was performed under our approved Washington University Embryonic Stem Cell Research Office (ESCRO) protocol #12-002. RFX4 heterozygous (HET) and homozygous (HOM) knockout hPSCs were generated in both the human embryonic stem cell (hESC) line WA09 and human induced pluripotent stem cell (hiPSC) line AN1.1 by targeting the first canonical exon (6) with CRISPR-Cas9-based genome engineering (Figure 1A, Figure S1A). The same process was employed to generate RFX3 homozygous (RFX3-HOM) knockout AN1.1 hiPSC models by targeting exon 8 (Figure 3G, Figure S6D). Finally, a patient-specific *RFX4* mutation (c.235C>T) was genetically engineered into the WA01 hESC line targeting one or both alleles of the *RFX4* gene with CRISPR-Cas9-based genome editing (Figure 6A, Figure S9A). The validity of all models was confirmed by NGS-based amplicon sequencing(Sentmanat et al., 2018), with all model generation and validation conducted by the Washington University Genome Engineering & Stem Cell Center (GESC). All hPSCs were maintained under feeder-free conditions on vitronectin (Gibco™) coated plasticware in StemFlex Medium (Gibco™) at 37°C with 5% CO2, passed upon reaching 70% confluency using ReLeSR (StemCell Technologies™) at a ratio of 1:6. Furthermore, cell lines were selected after genetic manipulation and karyotyped, demonstrating all cell lines were karyotypically normal (Figure S1D). Cell lines were additionally validated for the pluripotency markers SOX2 and OCT4 (Figure S1E) and were routinely confirmed mycoplasma, bacterial, and fungus-free. Experiments were carried out between passages 15-60 for all hPSC lines upon reaching 70-80% confluency. Details of cell backgrounds used for each experiment are detailed in Data S19.

### Method Details

#### Subcortical medial ganglionic eminence-like progenitor specification

To model subcortical inhibitory neuron development, we specified hPSCs as medial ganglionic eminence (MGE)-like progenitors with a ventral telencephalic progenitor (V-NPC, day 15) identity, as previously described (Meganathan et al., 2017). Embryoid bodies (EBs) were produced from hPSCs using AggreWell™800 microwell culture plates (STEMCELL Technologies) and AggreWell™ embryoid body formation medium (STEMCELL Technologies) supplemented with 10 µM Y-27632 (Y-27, Selleckchem). EBs were maintained for 3 days (day –3 to 0) before specification was initiated by switching to ventral specification media (VSM), constituting day 0 of differentiation. VSM was comprised of Neurobasal-A, B-27 supplement, Glutamax, β-Mercaptoethanol, Non-Essential Amino Acids, Penicillin-Streptomycin Solution (all from Gibco™) supplemented with 10 µM SB-431542 (SB, Selleckchem), 100 nM LDN-193189 (LDN, Tocris Biosciences), 100 nM Smoothened Agonist (SAG, Selleckchem), and 2 µM XAV-939 (XAV, Selleckchem). VSM was supplemented with 10 µM Y-27 from day 0 to day 2 and EBs were maintained on an orbital shaker at 80 rpm and fed every other day by half media replacement from day 0 to 10 of differentiation.

On day 10 of differentiation, EBs were plated onto Matrigel (Corning) and laminin (Sigma-Aldrich) coated plasticware and, after 24 hours, 20 ng/mL thermo-stable bFGF (Gibco™) was added to cells. Cells were fed by half media replacement with VSM on day 12 and 14. On day 15 of differentiation, cells were considered V-NPCs, as assessed by marker expression in our current and prior work (Meganathan et al., 2017). For immunofluorescence analysis, cells were replated using Versene (Gibco™) onto Matrigel and laminin coated plasticware and cultured for a further 2 days in VSM before being fixed with 4% paraformaldehyde (PFA).

### Cortical excitatory neuron differentiation

Modeling of cortical excitatory neuron development was carried out as described in our prior work (Chapman et al., 2022; Chapman et al., 2024) by specifying hPSCs as progenitors with a dorsal telencephalic progenitor regional character (D-NPC, day 20) and further differentiated into immature cortical excitatory neurons (cEXs, day 40). hPSCs were used to form EBs as described above and maintained in EB formation media from day –3 to 0. D-NPC specification was initiated by replacement of EB formation media with N2B27 supplemented with 10 µM SB, 100 nM LDN, 10 µM Y-27 and 2 µM XAV. N2B27 was comprised of 2/3 DMEM/F12, 1/3 Neurobasal™ Medium (Gibco™), N2 supplement (Gibco™), B-27 supplement (without Vitamin A), Glutamax, β-mercaptoethanol, non-essential amino acids and penicillin-streptomycin solution. EBs were fed on day 2 and 4 by half media replacement using N2B27 supplemented with SB, LDN and XAV, and from day 6 to 10 with N2B27 supplemented with SB and LDN. EBs were plated onto Matrigel and laminin coated plasticware on day 10 of differentiation and treated with 20 ng/mL thermo-stable bFGF after 24 hours. Cells were then fed with un-supplemented N2B27 every other day by half media replacement until day 20 of differentiation, at which point cells were considered D-NPCs as validated by molecular marker expression after our prior work.

To differentiate D-NPCs into cEXs, D-NPCs were dissociated using Accutase and plated onto Matrigel and laminin coated plasticware in N2B27+, comprised of 2/3 DMEM/F12, 1/3 Neurobasal™ medium, N2 supplement, B-27 supplement with Vitamin A (Gibco™), GlutaMAX, β-mercaptoethanol and penicillin-streptomycin solution. Cells were fed by half media replacement using N2B27+, supplemented with CultureOne™ supplement (Gibco™) from day 22-24 and 5 µM DAPT (Selleckchem) and 1 µM PD0332991 (PD, Selleckchem) from day 26-28. Cells were then maintained in N2B27+ supplemented with 20 ng/mL BDNF (R&D Systems), 200 µM ascorbic acid (Sigma-Aldrich) and 0.5 µM Cytarabine (R&D Systems) until days 40-50 of differentiation, with half media replacements performed every 2 days. Cells were collected as cEXs after 50 days of differentiation for puncta staining and after day 40 for all other cEX assays. For RNA-seq and CUT&Tag applications, cells were collected on day 20 as D-NPCs using Accutase. For immunocytochemistry, D-NPCs were replated using Versene at day 20 and maintained in N2B27– and fixed with 4% PFA 2 days after replating.

### Cerebral organoids

Dorsally patterned cortical organoids were generated as previously described(Lunn, 2022) with minor modifications and collected on days 30 and 60.EBs were formed as described above and then maintained in EB formation media for 4 days (day 0 to 4 of differentiation) on an orbital shaker at 80 rpm. On day 5 of differentiation, EBs were transferred into neural ectoderm (NE) media, which consisted of Neurobasal™ medium, N2 supplement, Glutamax and non-essential amino acids supplemented with 10 µM SB, and 100 nM LDN. EBs were fed by 1/2 media replacement every 2 days; on day 11 of differentiation, patterned EBs were individually embedded into Matrigel by using organoid embedding sheets (STEMCELL Technologies). Following embedding, patterned EBs were incubated at 37°C and 5% CO_2_ for 20 minutes, then removed from the embedding sheets and transferred into ultra-low adherence plasticware (Corning) in neural differentiation (ND) media. ND media was comprised of Neurobasal™ medium, B27 supplement (without Vitamin A), N2 supplement, Glutamax, non-essential amino acids, β-mercaptoethanol, 3 µg/mL Insulin (Sigma-Aldrich), 20 ng/mL epidermal growth factor (Gibco™), and 20 ng/mL thermally stable bFGF. Organoids were then fed by half media replacement using ND media from day 11-21 and using maturation media (MM) after day 21. MM was comprised of half DMEM/F12, half Neurobasal™ media, B27 supplement (without Vitamin A), N2 supplement, Glutamax, penicillin-streptomycin solution, non-essential amino acids, β-mercaptoethanol, 3 µg/mL Insulin, 10 µM cyclic adenosine monophosphate (Sigma-Aldrich), and 10 µM ascorbic acid.

Organoids were collected at day 30 and day 60, washed 3 times with PBS, and then fixed in 4% PFA overnight at 4°C. Following fixation, PFA was removed, and organoids were washed three times with PBS, followed by incubation in 30% (w/w) sucrose (Sigma-Aldrich) in PBS for a minimum of 24 hours at 4°C. Organoids were then embedded in 50:50 optimal cutting temperature (O.C.T.) compound (Fisher Scientific) in 30% sucrose solution and cryosectioned at 8-10 µm.

### RFX4 Overexpression

For RFX4 overexpression experiments, wildtype hPSCs were transduced with an RFX4 open reading frame plasmid (Genecopoeia, H2578-Lv203). Lentivirus for RFX4 overexpression was produced in Lenti-X-293T cells (Takara Bio) by transfecting the cells with envelope (pMD2.G), packaging (psPAX2), and RFX4 open reading frame plasmid (Genecopoeia, H2578-Lv203) at a ratio of 2:3:6 using Trans-IT Lenti transfection reagent (Mirus). pMD2.G and psPAX2 were gifts from Didier Trono (Addgene plasmids #12259 and #12260, respectively). Lenti-X-293T cells were grown in HEK media containing DMEM, high glucose, and no glutamine (Gibco™), and supplemented with 10% FBS (Gibco™), Glutamax, β-mercaptoethanol, Non-Essential Amino Acids and Penicillin-Streptomycin Solution, Sodium Pyruvate (Gibco™) and HEPES buffer (Gibco™). Lentivirus was concentrated using Lenti-X-Concentrator (Takara Bio) per the manufacturer’s instructions, re-suspended in Opti-MEM media (Gibco™), and stored at –80°C. Transduction of hESCs was aided by addition of 5 µg/mL polybrene (Sigma-Aldrich) immediately before addition of lentivirus. Cells were allowed to recover for 48 hours before selection with 500 ng/mL puromycin (Invitrogen™) was applied. Once cells were selected, they were maintained under selection for the duration of subsequent experiments. RFX4 overexpression was confirmed via RT-qPCR (Figure S4B); to test the ability of RFX4 overexpression to induce neuronal differentiation, cells were cultured in StemFlex from day 0-1 and N2B27+ from day 2 to 6. Cells were harvested for downstream analyses on day 6. D-NPCs were replated using versene and transduced with RFX4 ORF lentivirus on day 21. Cells collected for RFX4 overexpression validity (Figure S4I) and with downstream experiments on day 28 of differentiation.

### Induced Glutamatergic-like Neuron Differentiation

Induced glutamatergic-like neurons (iGluts) were generated by overexpressing the bHLH transcription factor Neurogenin-2 (NGN2) and as described in our prior work (Chapman et al., 2025). hPSCs were transduced with a lentivirus construct which induces expression of Neurogenin-2 (NGN2) in a Doxycycline-dependent manner, maintained in StemFlex, and selected with 0.5 µg puromycin. To start iGlut differentiation, hPSCs were dissociated using Accutase and plated on Matrigel-coated cultureware in StemFlex media supplemented with Y-27632 and puromycin. Cultures were allowed to recover for 48 hours before the addition of 4 µg/mL Doxycycline (Sigma-Aldrich) to induce NGN2 expression, which constituted day 0 of differentiation. Cultures were fed daily with full media changes of StemFlex from day 0 to 2 and then N2B27+ supplemented with Doxycycline and puromycin from day 3 to 5. From days 6 to 8, cultures were fed daily with N2B27+ supplemented with cytarabine (1µM); subsequently, cultures were fed every 2 days with half media changes of N2B27+ supplemented with 10 ng/mL BDNF. On day 21, cells were harvested as iGluts.

### Sphere Size and Outgrowth Measurements

Images were taken at day 12 of cortical inhibitory or excitatory neuron differentiation, using a 4x objective. A minimum of 15 neurospheres per biological replicate (n) were imaged, analyzing 4 biological replicates per condition. Sphere size and outgrowth were measured using ImageJ, with relative outgrowth defined as the area of cell outgrowth from the plated neurospheres, normalized to the corresponding sphere size. For details into on n values/biological replicates for each experiment, see Data S20.

### RNA-Sequencing

RNA-sequencing was performed as previously described (Chapman et al., 2024) on RFX4 LOF V-NPCs at day 15, RFX4 and RFX3 LOF and R79C D-NPCs at day 20, and WT and RFX4 HOM cEXs at day 40. All raw data is available from the Gene Expression Omnibus database under accession GSE316377. RNA was isolated using the NucleoSpin RNA Plus kit (Macherey-Nagel) and RNA-seq library preparation performed using the KAPA RNA HyperPrep Kit with RiboErase (Roche). Libraries were sequenced on an Illumina NovaSeq X Plus sequencer to a depth of ∼30 million (M) reads per sample. Library preparation and sequencing was performed by the Washington University Genome Technology Access Center at McDonnell Genome Institute (GTAC@MGI).

Raw reads from RNA-seq samples were quality trimmed using Cutadapt (Kechin et al., 2017) (v2.4) with the options of quality-cutoff=15,10 and minimum-length=36. Reads were aligned and quantified using hg38 GENCODE V42 comprehensive gene annotations using Salmon (Patro et al., 2017) in mapping-based mode correcting for sequence and GC bias (--seqBias –-gcBias) using options –– validateMappings and –-rangeFactorizationBins 4. Quantifications were summarized to gene level using the tximport (Soneson et al., 2015) (v1.32) R package using hg38 GENCODE V42 comprehensive gene annotations.

For differential expression analysis, only genes with ≥5 counts in ≥2 samples were included. Differential expression analysis was performed using DESeq2 (Love et al., 2014) in negative binomial mode, using counts for genes that passed the cutoff. A 1.5-fold linear expression change and a Benjamini and Hochberg false discovery rate of <0.05 (FDR; adjusted p-value) were set as cutoff values for significance. All significant differentially expressed gene output from each DESeq2 analysis before the above cutoffs were imposed are presented as supplementary datasets. Expression data is presented and visualized using transcript per million mapped reads (TPM) and all raw and processed RNA-seq data is available from the Gene Expression Omnibus database under accession GSE316377.

### Reverse Transcription and Quantitative PCR (RT-qPCR)

RNA was isolated as described above and quantified using a NanoDrop ND-1000 spectrophotometer (Thermo Scientific). Using ≥1 µg of RNA, cDNA was made with the High-Capacity cDNA Reverse Transcription Kit (Applied Biosystems). Equal quantities of cDNA were used as template for quantitative PCR (qPCR) using the Applied Biosystem StepOne Plus or Quantstudio 3 quantitative PCR system. The endogenous loading controls used for each experiment are detailed in the corresponding figures legends, with all primers used detailed in the Key Resource Table, Oligonucleotides. Data was generated from a minimum of 3 biological replicates (n), each conducted in technical triplicate. All statistics were performed using relative abundance and data is presented as fold change relative to its appropriate control unless otherwise stated in figure legends. RT-qPCR primers are detailed below in the Key Resource Table, Oligonucleotides.

### Immunocytochemistry

For immunocytochemistry in monolayer cultures, cells were fixed in 4% PFA (Sigma-Aldrich) before blocking using 5% donkey serum (Sigma-Aldrich) plus 0.1% Triton (Sigma-Aldrich) in PBS. Primary antibodies were added overnight at 4°C before cells were washed with PBS and incubated with secondary antibodies for 1 hour at room temperature. Cells were counterstained with DAPI (Sigma-Aldrich) for 3 minutes where appropriate and then washed with PBS and mounted with ProLong™ Glass Antifade Mountant (Invitrogen™). Images were obtained using an epifluorescence microscope and processed with ImageJ. Quantification was performed using a minimum of 3 random images taken from at least 3 independent biological replicates using CellProfiler (Stirling et al., 2021), requiring a minimum of 60% overlap between immunofluorescence signal and DAPI counterstaining for co-localization to be scored. For puncta staining, quantification was performed using Intellicount (Fantuzzo et al., 2017), requiring pre– and post-synaptic puncta to be localized to TUBB3+ neurites with data represented as puncta density normalized to total TUBB3 area within each image.

For immunocytochemistry of organoid sections, antigen retrieval was performed in 1X citrate buffer (Sigma) for 12 minutes at 80°C before washing sections with TBS containing 0.1% Tween-20 (Sigma-Aldrich) and blocking in TBS containing 1% Triton and 5% donkey serum. Primary antibodies were incubated overnight in TBS containing 0.5% Triton and 5% donkey serum at 4°C. Sections were then washed 3 times in TBS containing 0.1% Tween-20 (TBST) for 5 minutes and incubated with secondary antibodies for 1 hour at room temperature. Sections were washed an additional 3 times with TBST before counterstaining with DAPI for 3 minutes, followed by an additional 3 TBST washes. Sections were then mounted with ProLong™ Glass Antifade Mountant. Images were obtained using a Zeiss AxioScan Z1 and quantified with CellProfiler (Stirling et al., 2021), requiring a minimum of 60% overlap between immunofluorescence signal and DAPI counterstaining to score co-localization. TUBB3 area was calculated as the total area of TUBB3 positive neurites normalized relative to the total number of DAPI+ cell nuclei. All primary and secondary antibodies used are detailed in the Key Resource Table, Antibodies.

### Western Blotting

Protein was isolated in RIPA buffer (Abcam) supplemented with Halt™ Protease and Phosphatase Inhibitor Cocktail (Thermo Scientific™) and western blotted using Bolt™ 4-12%, Bis-Tris, 1.5 mm, Mini Protein Gels (Invitrogen™). Proteins were transferred to 0.2 µm nitrocellulose membranes (Bio-Rad) and nonspecific antibody binding was blocked by incubation for 1 hour in a 5% milk solution. Blots were incubated with primary antibodies (Key Resource Table, Antibodies) overnight at 4°C and secondary HRP conjugated antibodies (Invitrogen), were applied at a 1:5,000 dilution for 1 hour at room temperature after blots were washed. Visualization was performed by chemiluminescence using the SuperSignal™ West Pico PLUS Chemiluminescent Substrate (Thermo Scientific™), all band intensities were normalized to GAPDH loading controls, and a minimum of 3 biological replicates were used for statistical analysis. Statistical analysis was performed on normalized signal intensity, data is presented as fold change compared to appropriate controls, and complete blots are presented in the supplemental figures with raw images available in Data S21. All antibodies used in this study are detailed in the Key Resource Table, Antibodies.

### CUT&Tag

CUT&Tag was performed as in our prior work (Chapman et al., 2024) using 2.5×10^5^ cells/sample and ≥3 independent biological replicates. Primary and secondary antibodies are listed in the Key Resource Table, Antibodies. After library preparation, samples with unique dual-end indexes were pooled in equimolar concentrations and were sequenced by GTAC@MGI on the NovaSeq X Plus sequencer as 150 bp paired-end reads at a depth of ≥5 million (M) reads per sample. Raw reads from CUT&Tag samples were processed by AIAP (Liu et al., 2021) (v1.1) using human genome hg38 as a reference to perform read quality control, alignment, quantification, and peak calling. For downstream analysis of RFX3/4 binding, bound sites identified in respective HOM models were eliminated as likely non-specific antibody binding. Raw data is available from the gene expression omnibus under accession GSE316377.

Differential binding analysis was performed using DiffBind (Ross-Innes et al., 2012), using trimmed and aligned binary alignment and map files along with peaks identified by MACS2 within the AIAP (v1.1) workflow. Blacklisted sequences were removed, and a minimum peak overlap of 3 samples was used to generate the consensus peak set for each biological condition. Read depth data was normalized using the Reads in Peaks library normalization method and significantly differentially bound peaks were identified using DESeq2, with a Benjamini and Hochberg FDR of <0.05. Complete result tables for all differential binding analysis are presented in supplementary data. Peak sets, including differentially bound peaks, were annotated using ChIPseeker and curated peak sets were visualized using deepTools (Ramirez et al., 2016).

Footprinting analysis was conducted using TOBIAS (Bentsen et al., 2020), merging all RFX4 genome-wide binding data prior to ATAC correction and using peaks identified in all 4 replicates of control D-NPCs but not HOM D-NPCs as the peak set. Deep learning modeling was performed using ChromBPNet (Pampari et al., 2025) using all RFX4 genome-wide binding data to generate both the bias and bias-factorized models. Multiple bias-factorized models were trained using separate sets of chromosomes for training and validation with the results averaged after contribution scores were calculated. Denovo motif discovery was performed using TF-MoDISco (Pampari et al., 2025), utilizing all seqlets identified across bias-factorized models and results were compared to the JASPAR2022 database of vertebrate TFBS motifs. Finally, instances of motifs identified by motif discovery which contribute to the predictions of the bias-factorized models were called using the GPU-accelerated hit caller finemo_gpu (Pampari et al., 2025). All antibodies used in this study are detailed in the Key Resource Table, Antibodies.

### Luciferase Assays

Oligonucleotides used to amplify RFX4 bound sites for subcloning into a luciferase vector were based on RFX4 binding sites and centered on an RFX TFBS, extending sequences to 119 base pairs total (Key Resource Table, Oligonucleotides). Each amplicon subcloned into the pTK-Gaussia Luc Vector (Thermo Scientific™) using XhoI and HindIII restriction enzyme sites, with successful cloning confirmed by colony PCR. Bacterial transformation was used to introduce each plasmid into NEB® Stable Competent E. coli and plasmid stocks were isolated using the PureLink™ HiPure Plasmid Midiprep Kit (Invitrogen™). Each pTK-Gaussia plasmid was cotransfected at equimolar concentrations at a 1:1 ratio with the pTK-Cypridina Luc Vector into D-NPCs using Lipofectamine™ 3000 Transfection Reagent (Invitrogen™). Supernatant was collected 72-hours post transfection and luciferase signal from Cypridina luciferase was measured using the Pierce™ Cypridina Luciferase Flash Assay Kit (Thermo Scientific™), while Gaussia luciferase was measured using Pierce™ Gaussia Luciferase Flash Assay Kit (Thermo Scientific™) on a Synergy2 instrument. All signals from experimental reporter constructs were normalized to signal from the empty pTK-Gaussia Luc plasmid, with the Cypridina luciferase signal used as the loading control. Results are presented as fold difference in HOM D-NPCs relative to the WT signal. Oligonucleotides are detailed below in the Key Resource Table under Oligonucleotides.

### Quantification and Statistical Analysis

Where appropriate, statistical analysis was carried out using a combination of tools available in GraphPad Prism version 9 (GraphPad Software; La Jolla, CA, USA, available from www.graphpad.com) or RStudio version 3.5.1 (RStudio: Integrated development environment for R; Boston, MA, USA. Available from www.rstudio.org). All technical replicates were averaged before statistical analysis and statistical tests used for analysis of each data set are detailed in the figure legends or in methods section for specific analysis paradigms, including differential gene expression and differential binding analysis. A minimum of 3 independent differentiations (3 biological replicates) were used for each time point and biological condition, with the number of differentiations used for each sample listed in the figure legends as “n”. The results in figures are presented as group mean +/-standard error (SE) indicating each biological replicate used for the analysis, unless otherwise specified in figure legends. Statistical significance is indicated with asterisks as follows: ns, not significant; ∗, p<0.05; ∗∗, p<0.01; ∗∗∗, p<0.001; ∗∗∗∗, p<0.0001 unless otherwise specified in figure legends.

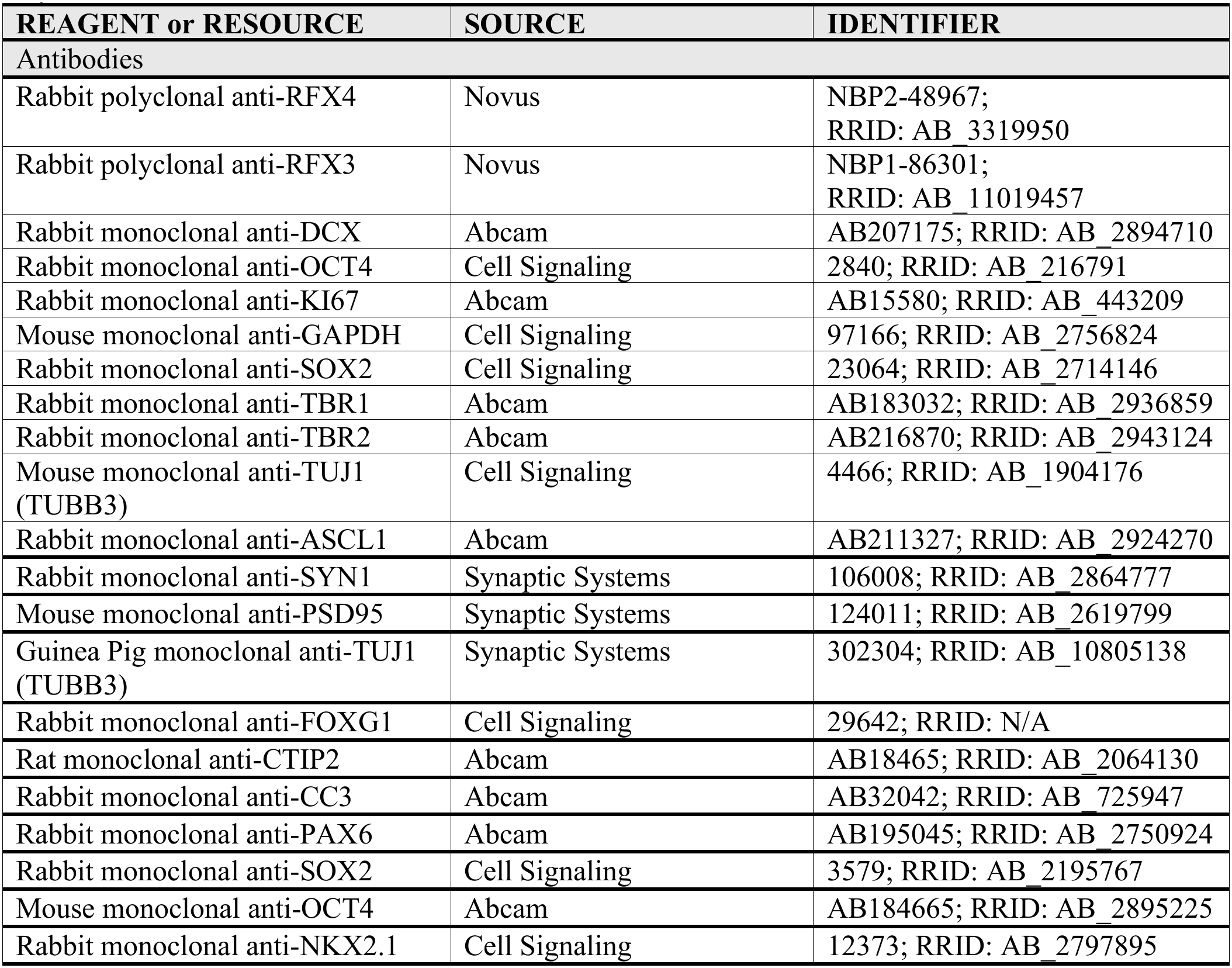
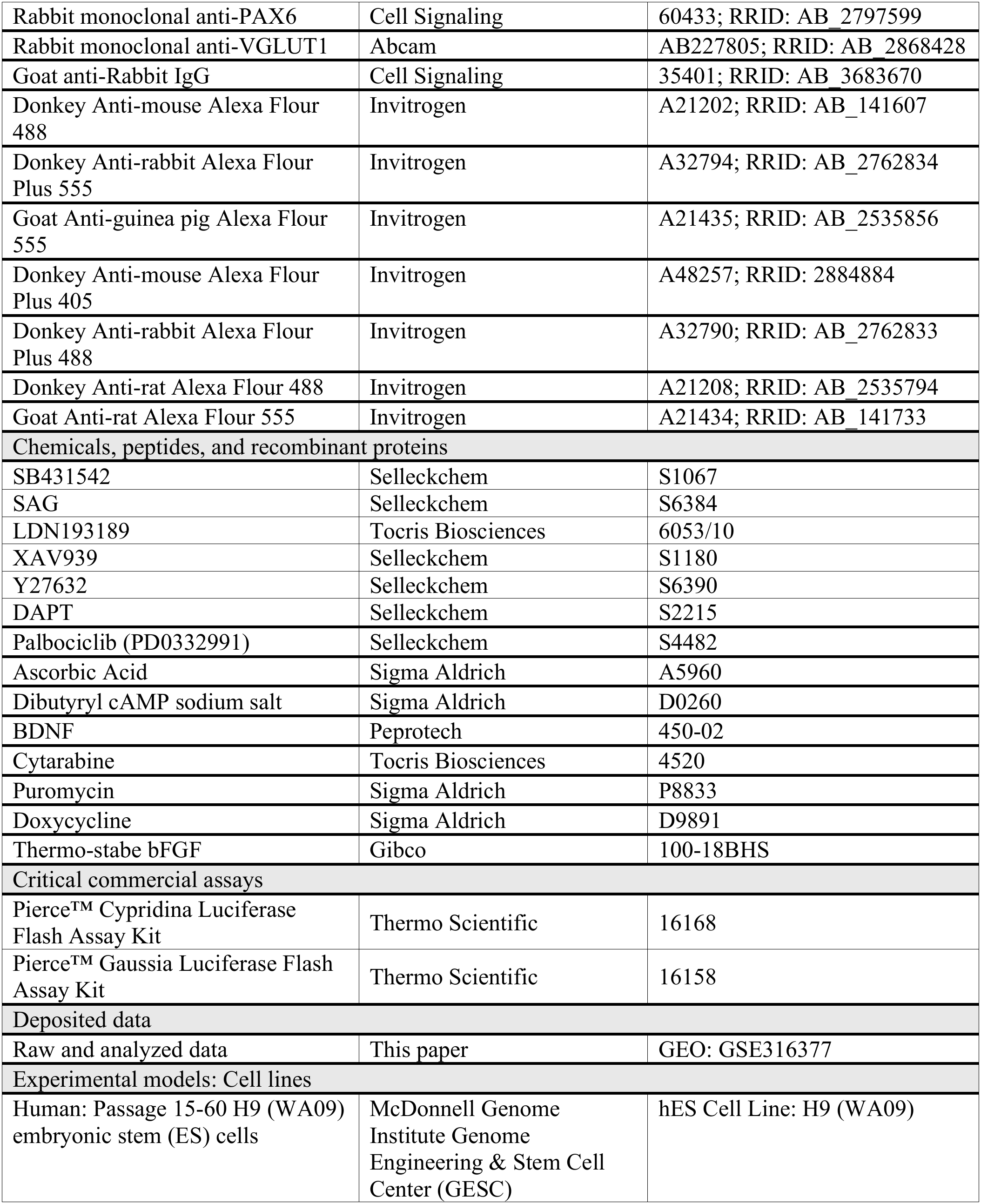
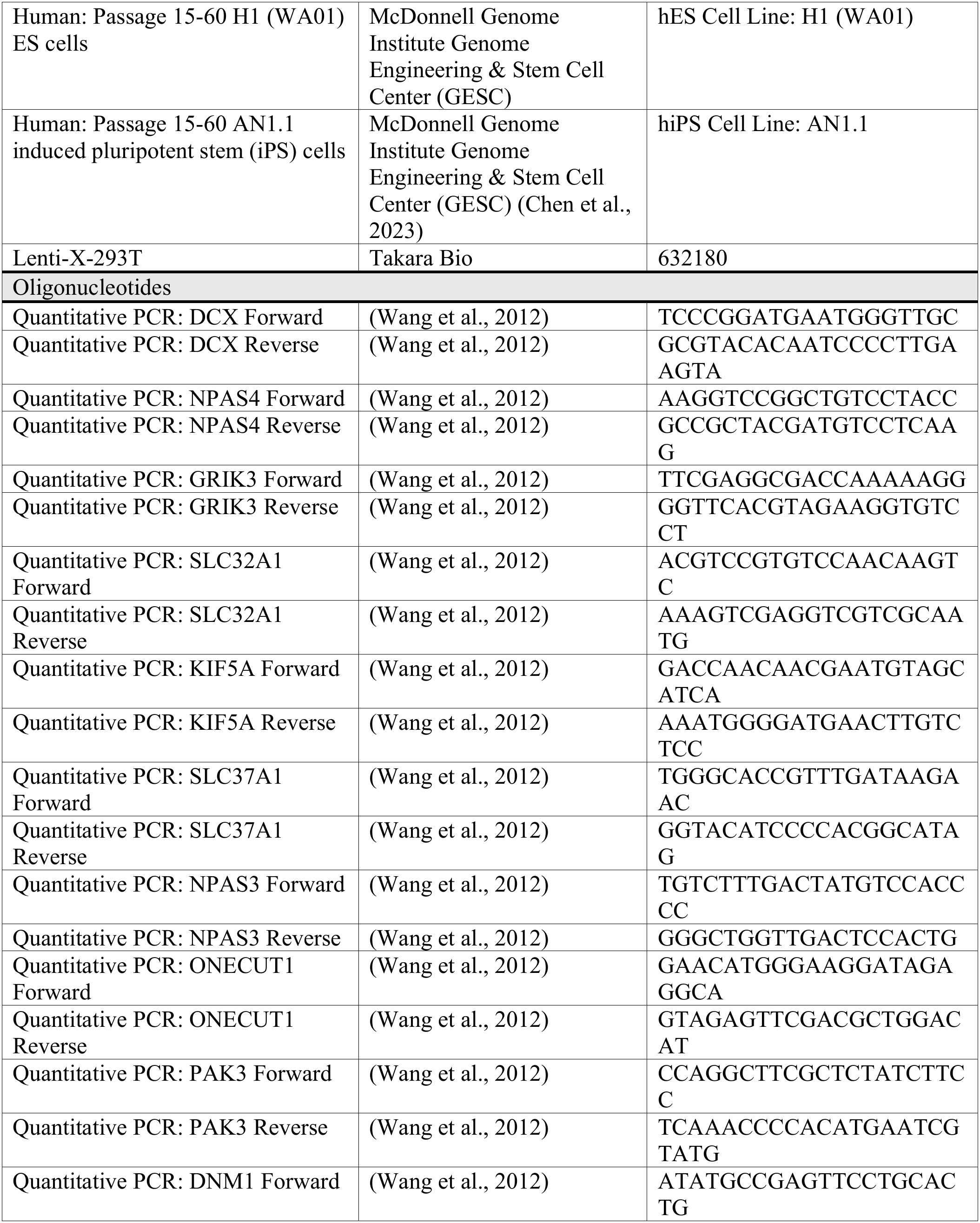
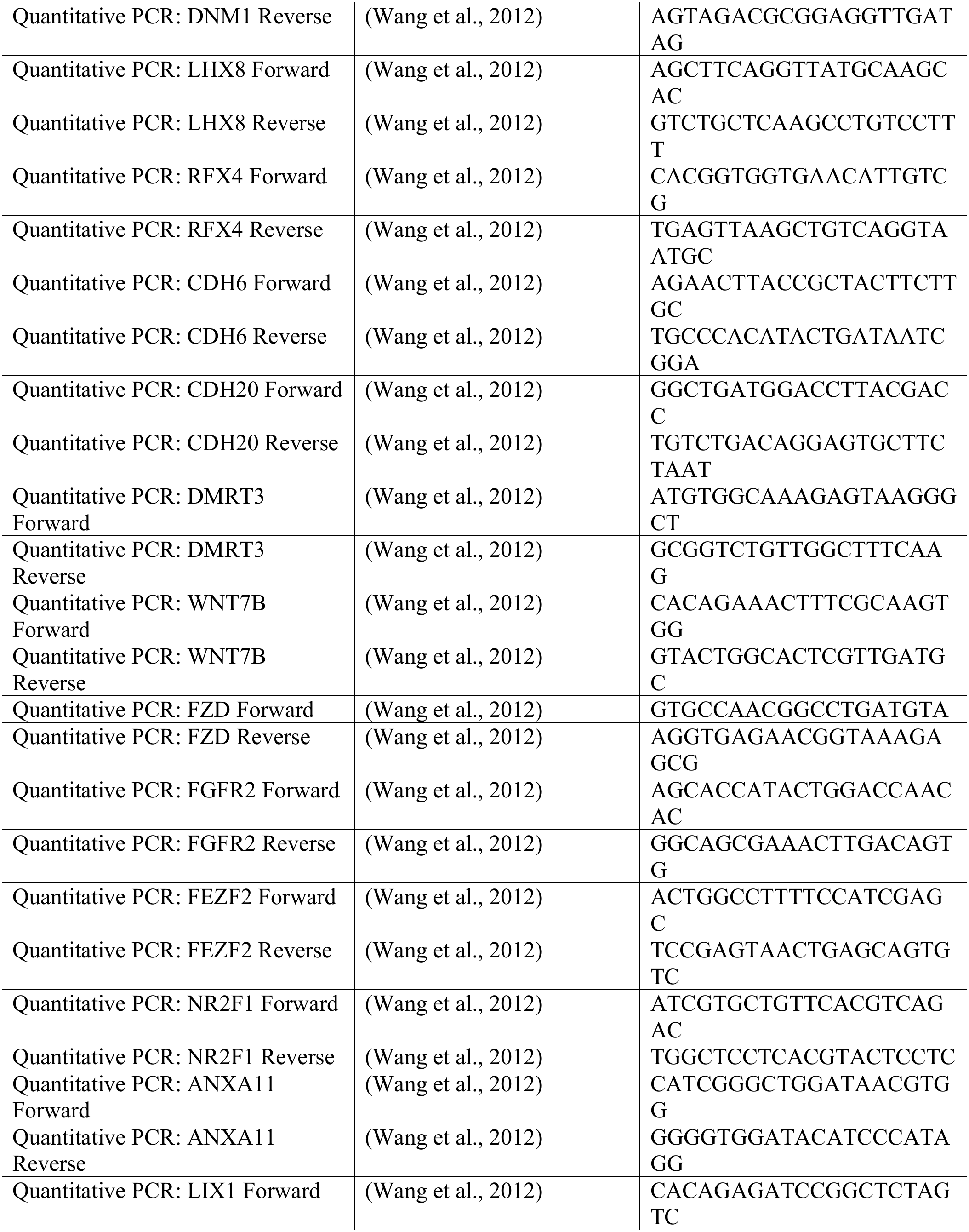
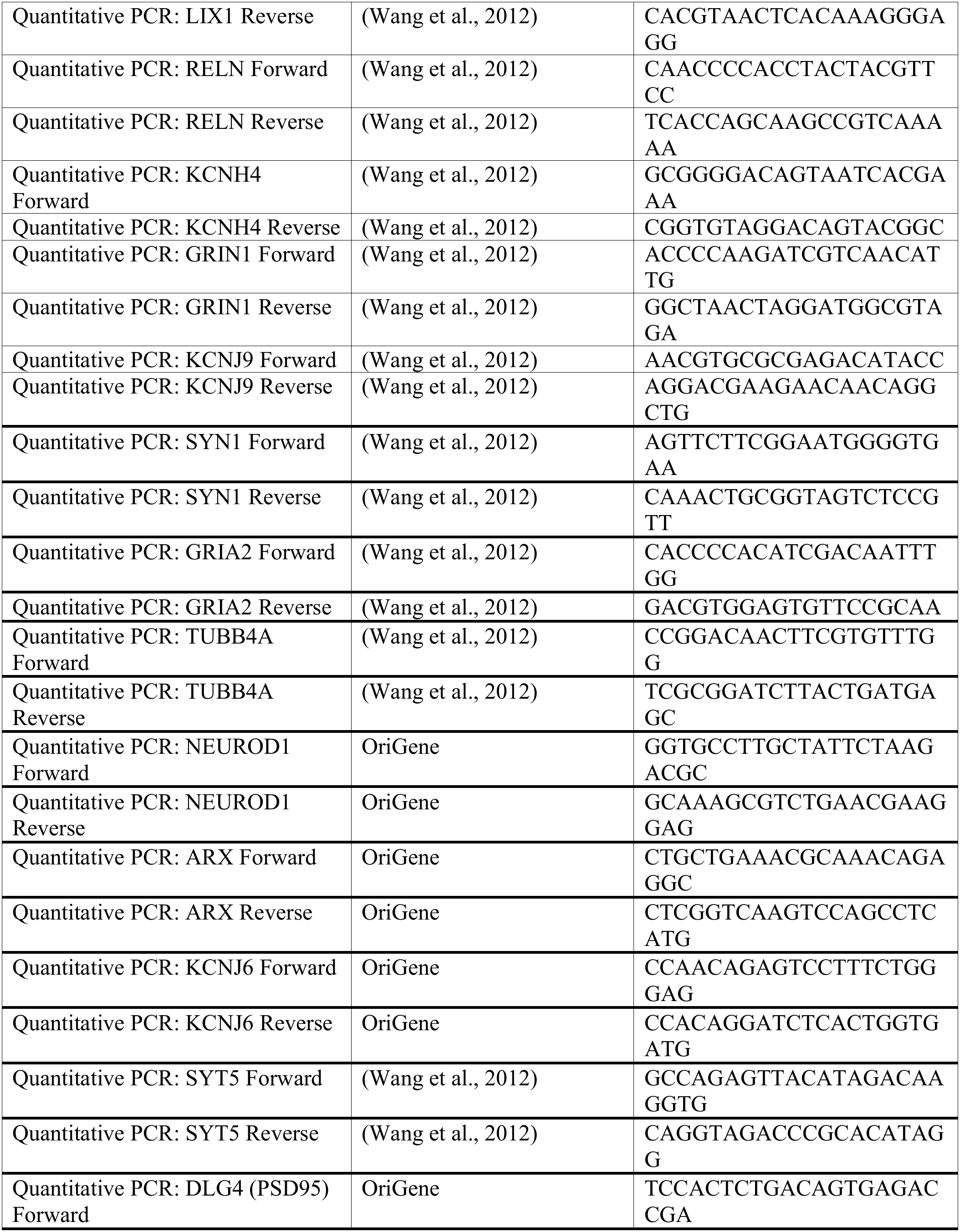
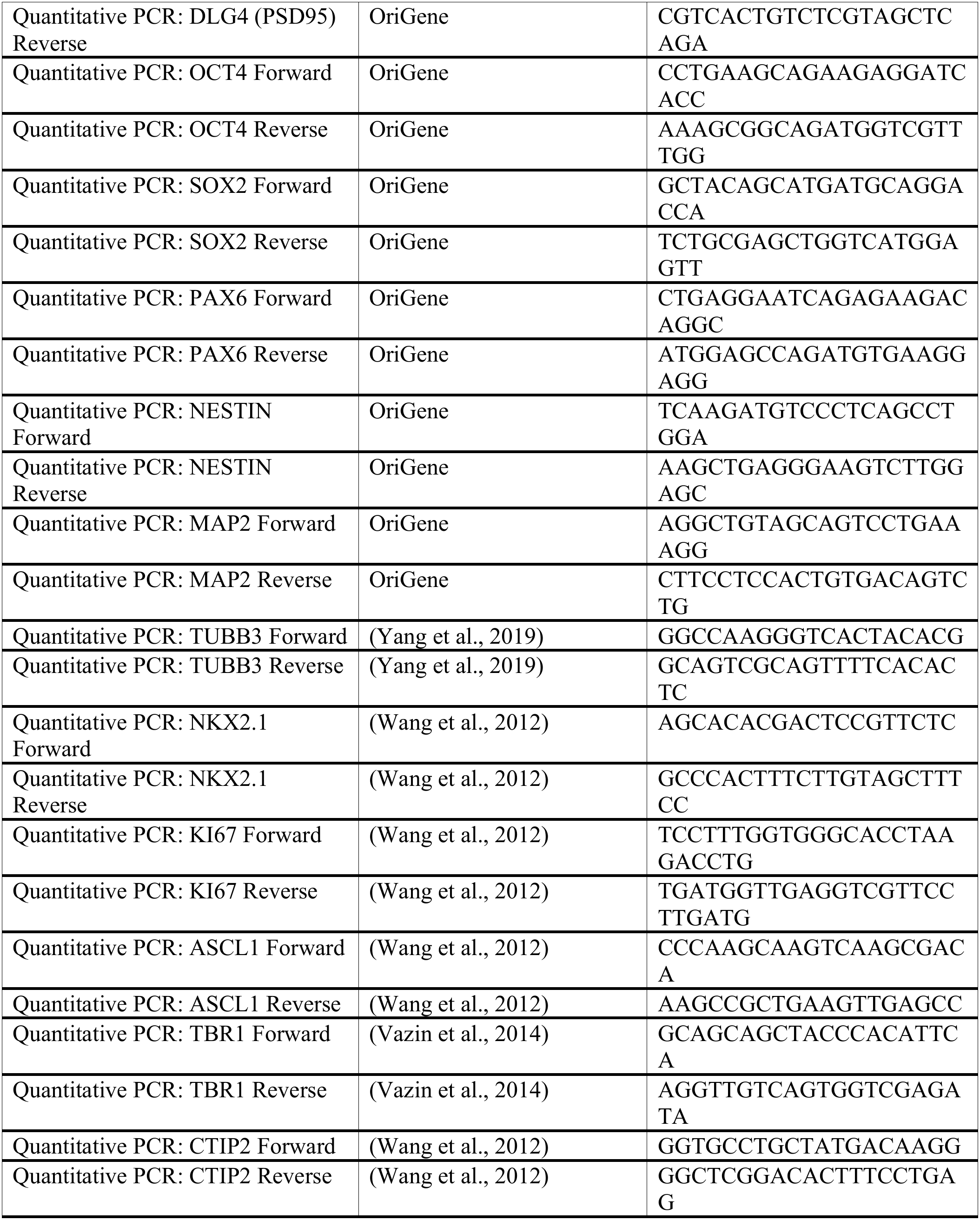
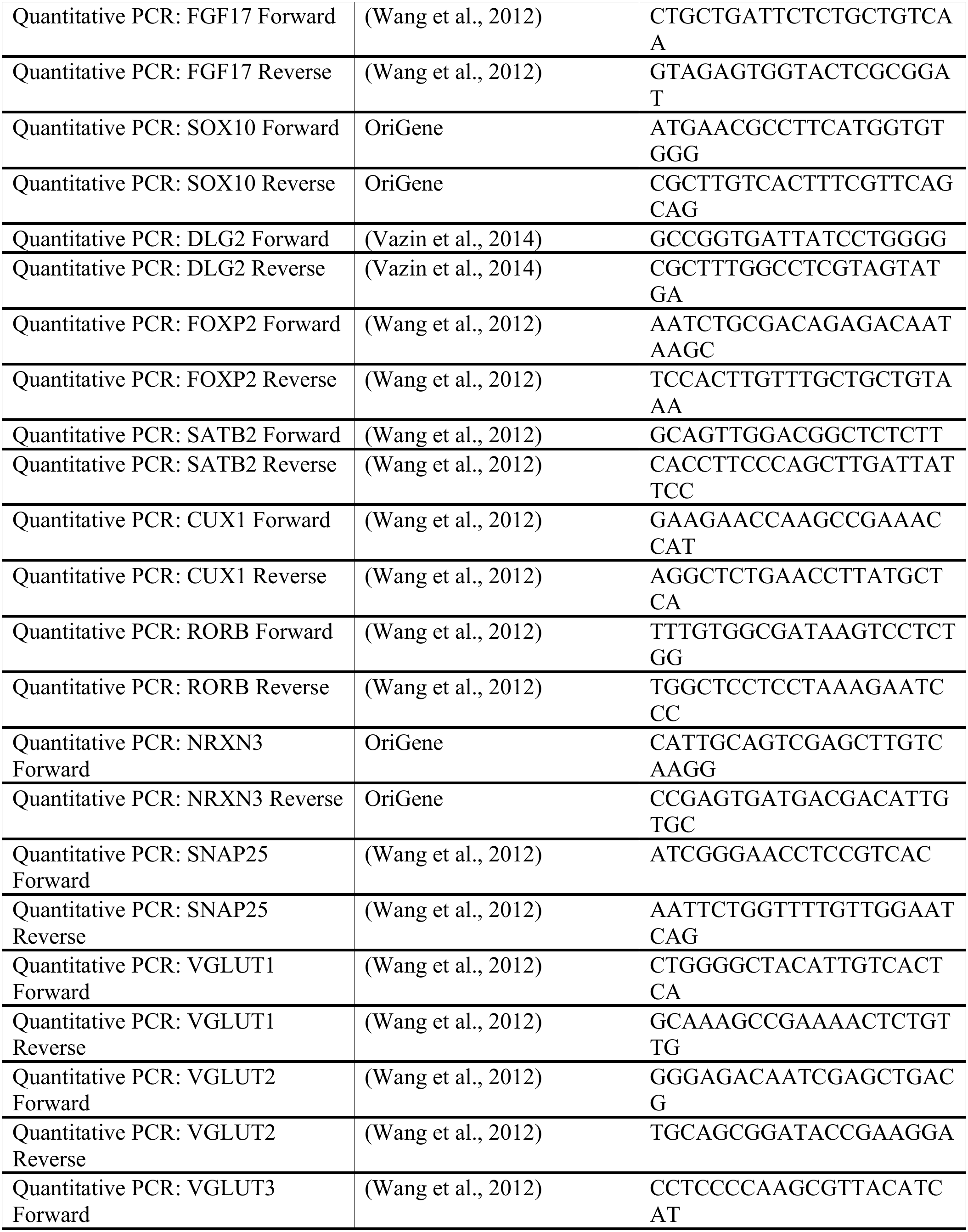
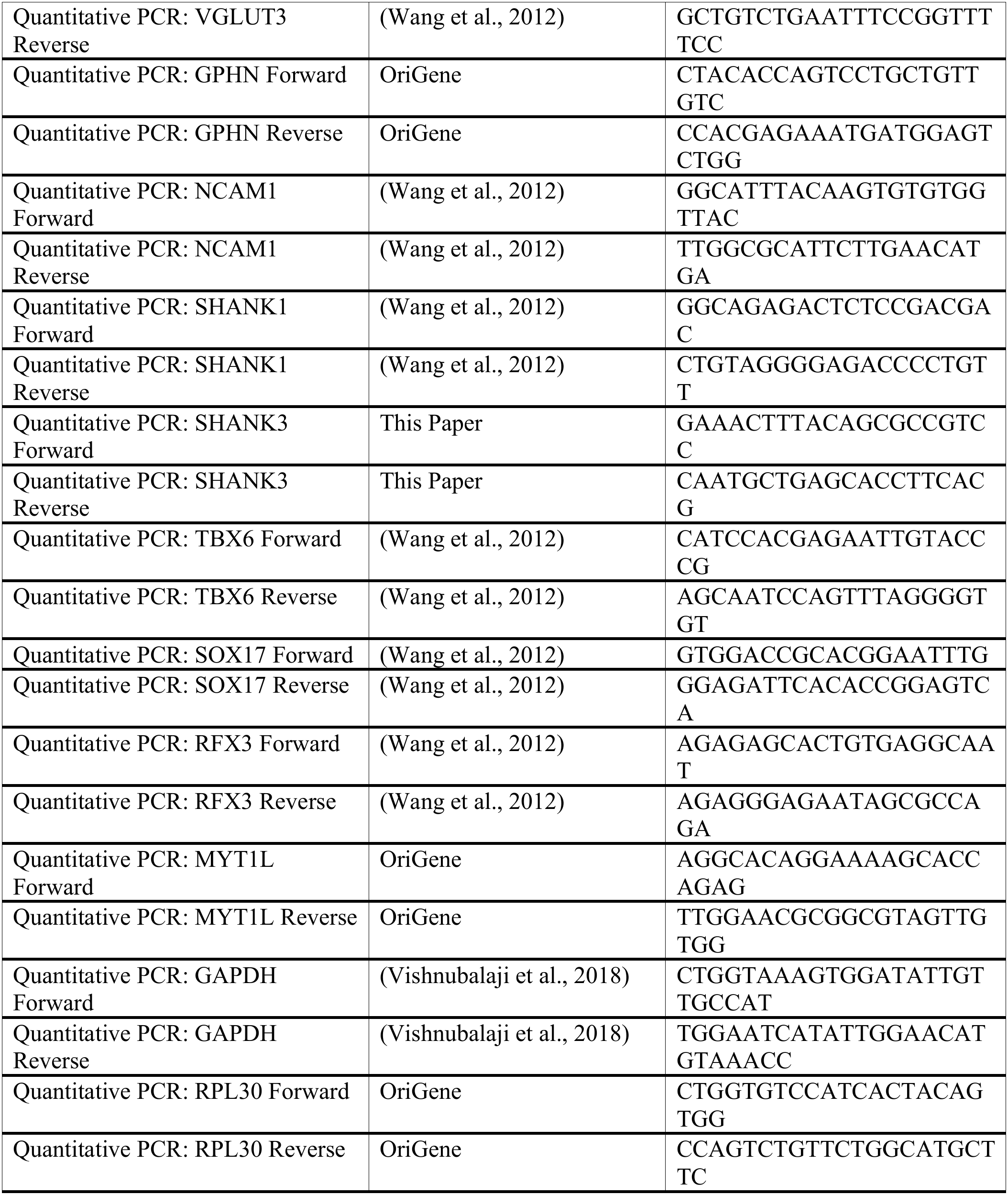
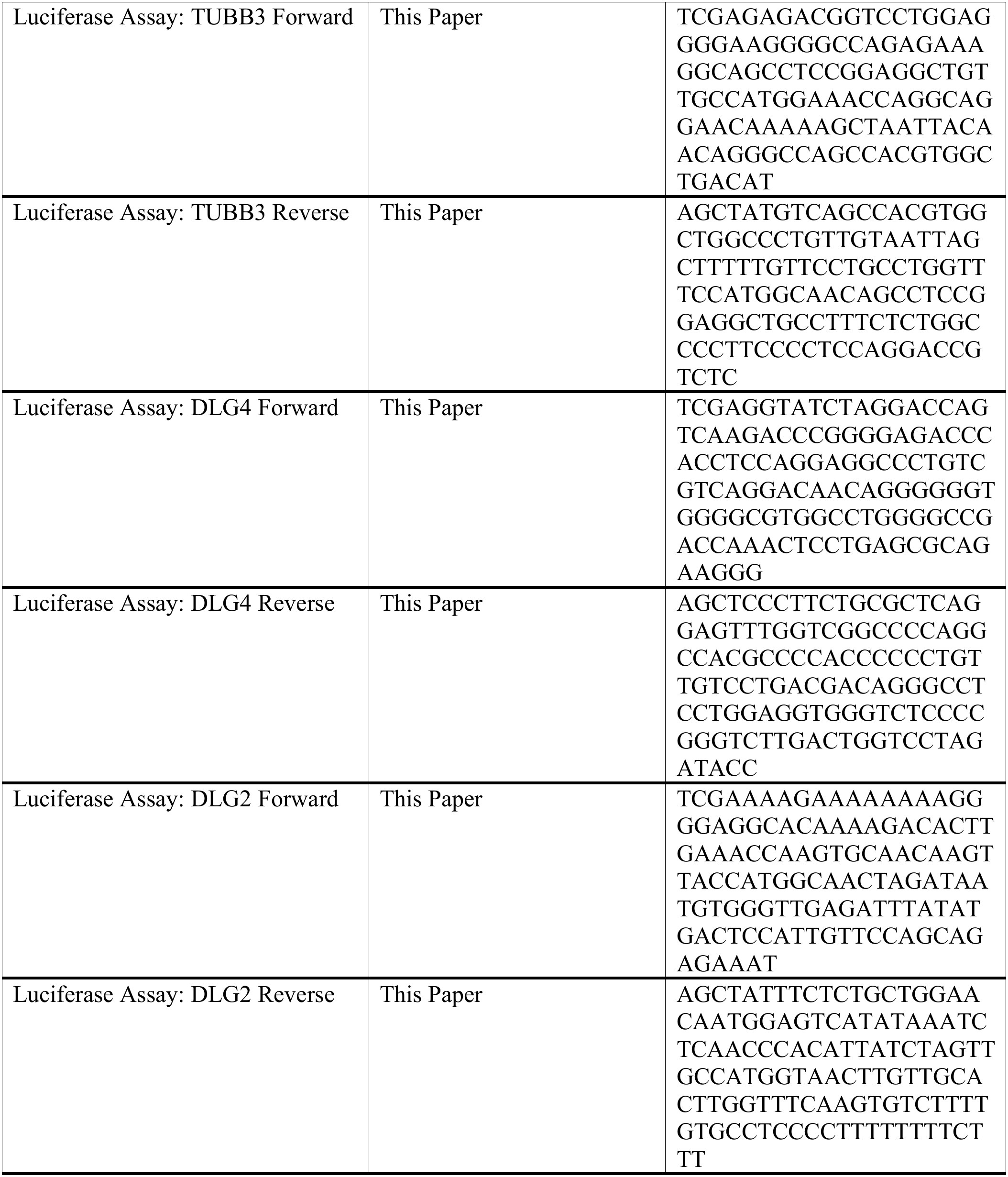
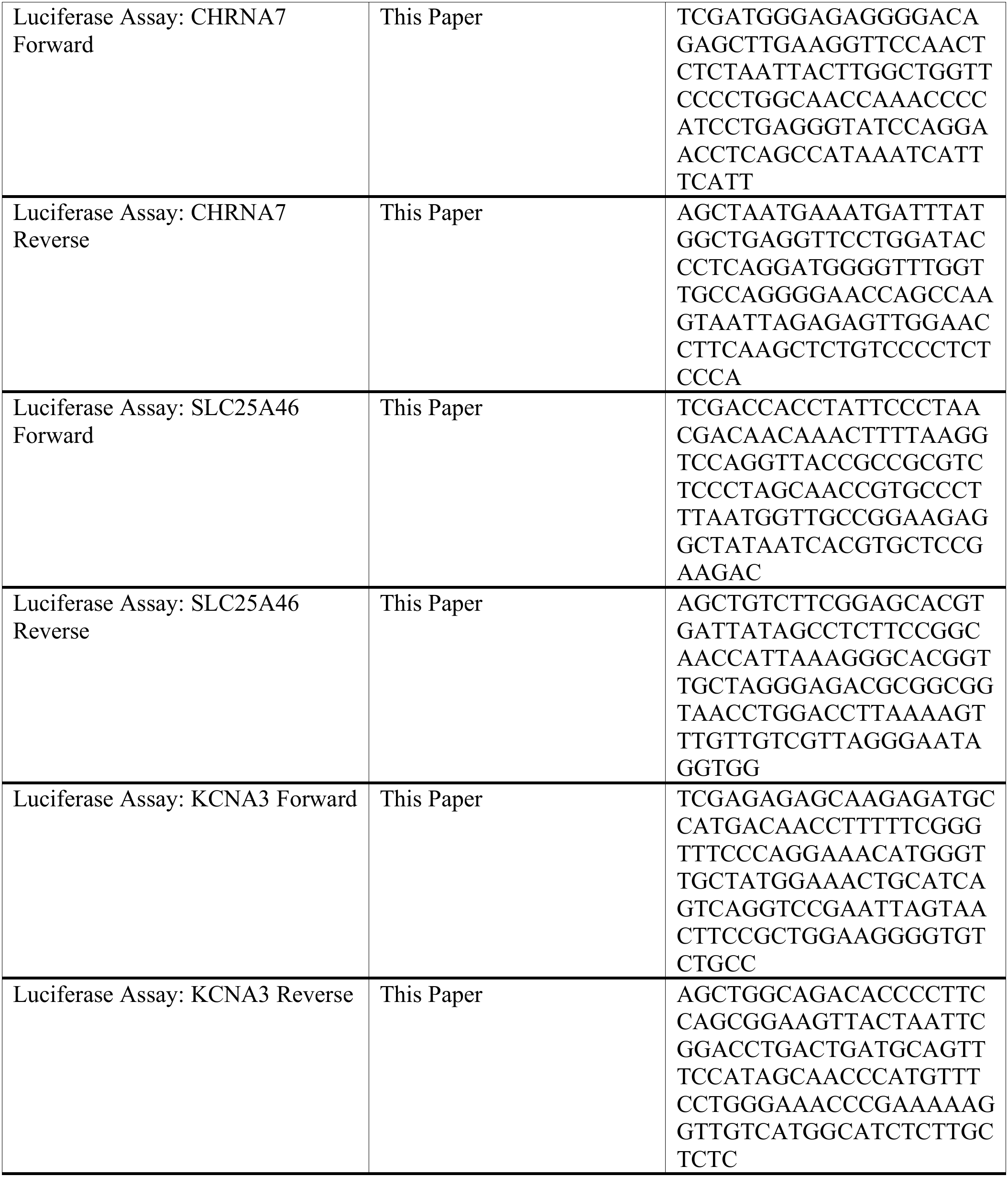
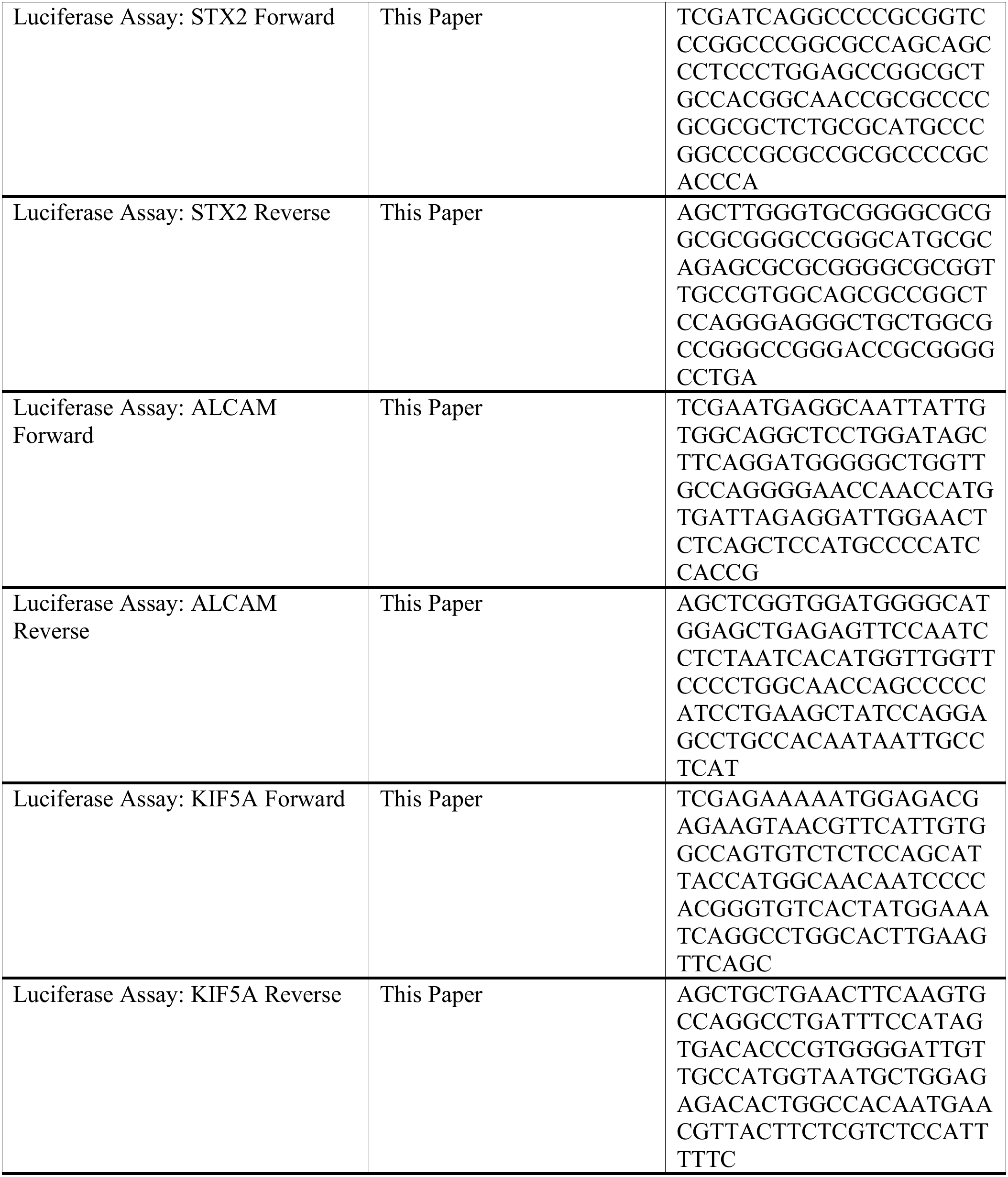
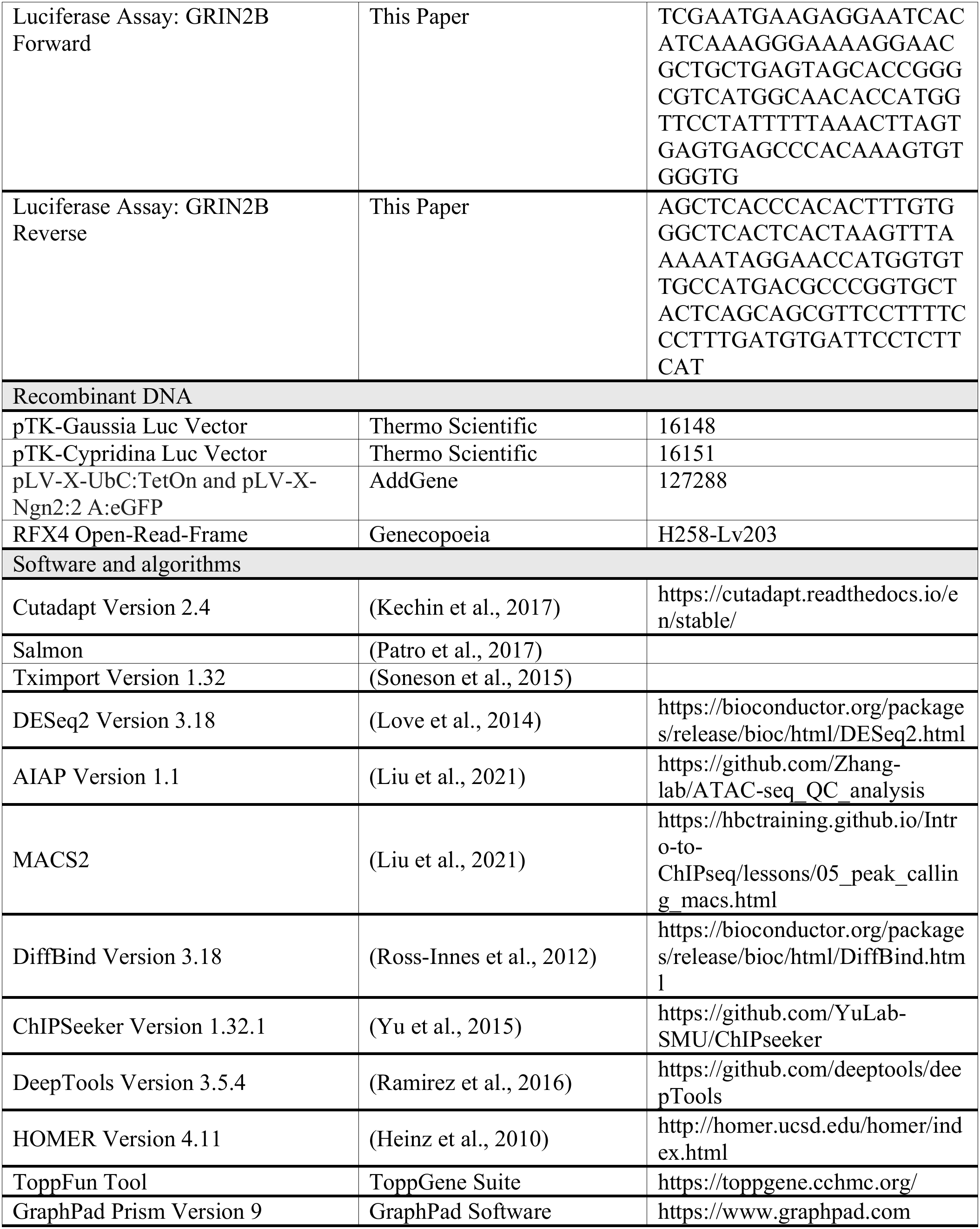
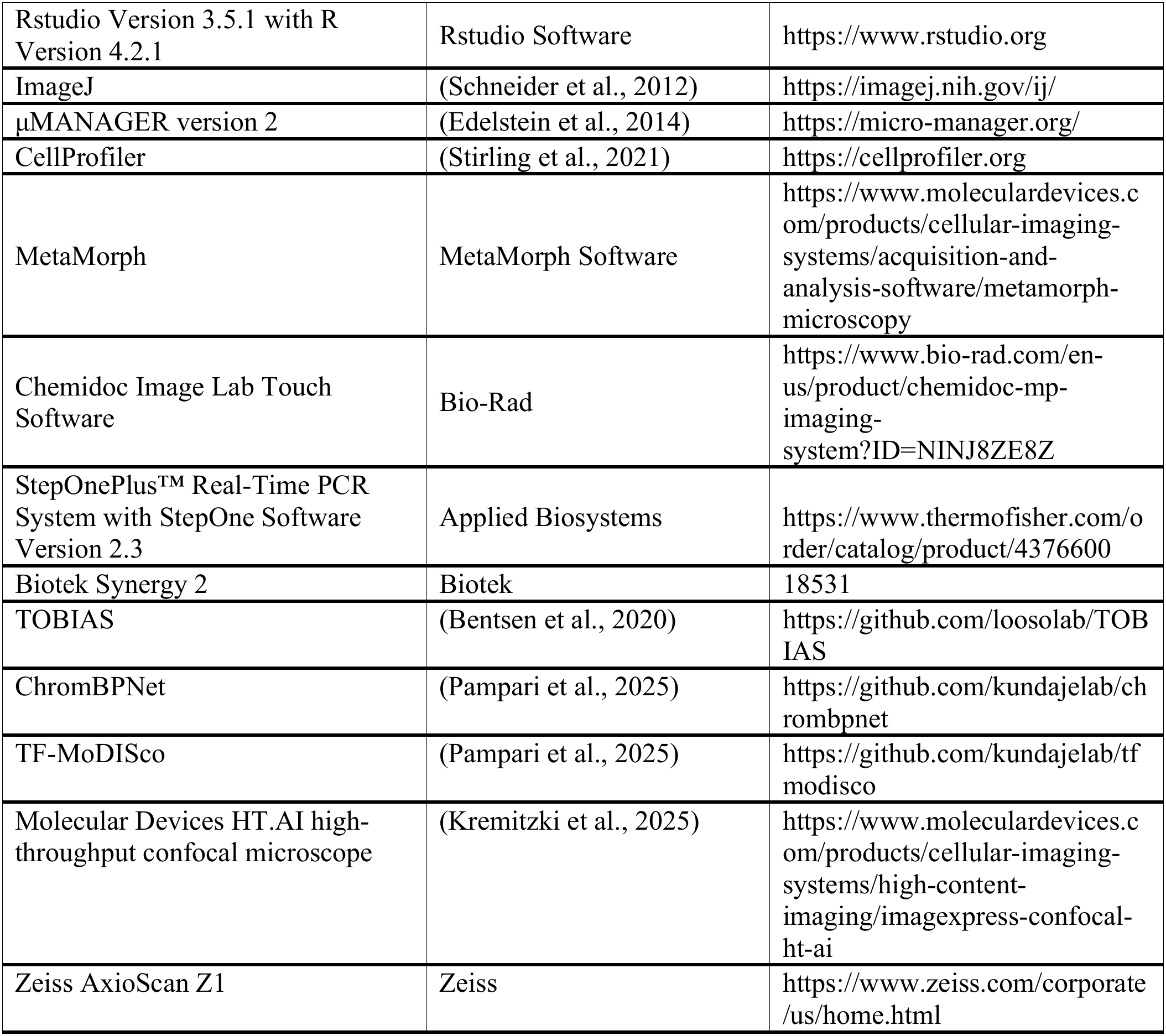
Key Resource Table.

## References

1. Adam, M. A., & Harwell, C. C. (2020). Epigenetic regulation of cortical neurogenesis; orchestrating fate switches at the right time and place. Curr Opin Neurobiol, 63, 146–153. 10.1016/j.conb.2020.03.012

2. Ajami, N., Kerachian, M. A., Toosi, M. B., Ashrafzadeh, F., Hosseini, S., Robinson, P. N., & Abbaszadegan, M. R. (2023). Inherited deletion of 9p22.3-p24.3 and duplication of 18p11.31-p11.32 associated with neurodevelopmental delay: Phenotypic matching of involved genes. J Cell Mol Med, 27(4), 496–505. 10.1111/jcmm.17662

3. Albert, M., Kalebic, N., Florio, M., Lakshmanaperumal, N., Haffner, C., Brandl, H., Henry, I., & Huttner, W. B. (2017). Epigenome profiling and editing of neocortical progenitor cells during development. EMBO J, 36(17), 2642–2658. 10.15252/embj.201796764

4. Ashique, A. M., Choe, Y., Karlen, M., May, S. R., Phamluong, K., Solloway, M. J., Ericson, J., & Peterson, A. S. (2009). The Rfx4 transcription factor modulates Shh signaling by regional control of ciliogenesis. Sci Signal, 2(95), ra70. 10.1126/scisignal.2000602

5. Banerjee, I., Senniappan, S., Laver, T. W., Caswell, R., Zenker, M., Mohnike, K., Cheetham, T., Wakeling, M. N., Ismail, D., Lennerz, B., Splitt, M., Berberoglu, M., Empting, S., Wabitsch, M., Potzsch, S., Shah, P., Siklar, Z., Verge, C. F., Weedon, M. N., … Flanagan, S. E. (2019). Refinement of the critical genomic region for congenital hyperinsulinism in the Chromosome 9p deletion syndrome. Wellcome Open Res, 4, 149. 10.12688/wellcomeopenres.15465.2

6. Bentsen, M., Goymann, P., Schultheis, H., Klee, K., Petrova, A., Wiegandt, R., Fust, A., Preussner, J., Kuenne, C., Braun, T., Kim, J., & Looso, M. (2020). ATAC-seq footprinting unravels kinetics of transcription factor binding during zygotic genome activation. Nat Commun, 11(1), 4267. 10.1038/s41467-020-18035-1

7. Blackshear, P. J., Graves, J. P., Stumpo, D. J., Cobos, I., Rubenstein, J. L., & Zeldin, D. C. (2003). Graded phenotypic response to partial and complete deficiency of a brain-specific transcript variant of the winged helix transcription factor RFX4. Development, 130(19), 4539–4552. 10.1242/dev.00661

8. Cadwell, C. R., Bhaduri, A., Mostajo-Radji, M. A., Keefe, M. G., & Nowakowski, T. J. (2019). Development and Arealization of the Cerebral Cortex. Neuron, 103(6), 980–1004. 10.1016/j.neuron.2019.07.009

9. Chapman, G., Alsaqati, M., Lunn, S., Singh, T., Linden, S. C., Linden, D. E. J., van den Bree, M. B. M., Ziller, M., Owen, M. J., Hall, J., Harwood, A. J., & Syed, Y. A. (2022). Using induced pluripotent stem cells to investigate human neuronal phenotypes in 1q21.1 deletion and duplication syndrome. Mol Psychiatry, 27(2), 819–830. 10.1038/s41380-021-01182-2

10. Chapman, G., Determan, J., Edwards, J. R., Huettner, J. E., Crump, S., Jetter, H., Gabel, H. W., & Kroll, K. L. (2025). TBRS-associated DNMT3A mutations disrupt cortical interneuron differentiation and neuronal networks. bioRxiv. 10.1101/2025.06.24.661324

11. Chapman, G., Determan, J., Jetter, H., Kaushik, K., Prakasam, R., & Kroll, K. L. (2024). Defining cis-regulatory elements and transcription factors that control human cortical interneuron development. iScience, 27(6), 109967. 10.1016/j.isci.2024.109967

12. Chen, Y. H., Connelly, J. P., Florian, C., Cui, X., & Pruett-Miller, S. M. (2023). Short tandem repeat profiling via next-generation sequencing for cell line authentication. Dis Model Mech, 16(10). 10.1242/dmm.050150

13. Choi, W., Choe, M. S., Kim, S. M., Kim, S. J., Lee, J., Lee, Y., Lee, S. M., Dho, S. H., Lee, M. Y., & Kim, L. K. (2024). RFX4 is an intrinsic factor for neuronal differentiation through induction of proneural genes POU3F2 and NEUROD1. Cell Mol Life Sci, 81(1), 99. 10.1007/s00018-024-05129-y

14. Cholfin, J. A., & Rubenstein, J. L. (2007). Genetic regulation of prefrontal cortex development and function. Novartis Found Symp, 288, 165–173; discussion 173-167, 276-181. https://www.ncbi.nlm.nih.gov/pubmed/18494258

15. Ciceri, G., Baggiolini, A., Cho, H. S., Kshirsagar, M., Benito-Kwiecinski, S., Walsh, R. M., Aromolaran, K. A., Gonzalez-Hernandez, A. J., Munguba, H., Koo, S. Y., Xu, N., Sevilla, K. J., Goldstein, P. A., Levitz, J., Leslie, C. S., Koche, R. P., & Studer, L. (2024). An epigenetic barrier sets the timing of human neuronal maturation. Nature, 626(8000), 881–890. 10.1038/s41586-023-06984-8

16. Edelstein, A. D., Tsuchida, M. A., Amodaj, N., Pinkard, H., Vale, R. D., & Stuurman, N. (2014). Advanced methods of microscope control using muManager software. J Biol Methods, 1(2). 10.14440/jbm.2014.36

17. Fantuzzo, J. A., Mirabella, V. R., Hamod, A. H., Hart, R. P., Zahn, J. D., & Pang, Z. P. (2017). Intellicount: High-Throughput Quantification of Fluorescent Synaptic Protein Puncta by Machine Learning. eNeuro, 4(6). 10.1523/ENEURO.0219-17.2017

18. Gajiwala, K. S., Chen, H., Cornille, F., Roques, B. P., Reith, W., Mach, B., & Burley, S. K. (2000). Structure of the winged-helix protein hRFX1 reveals a new mode of DNA binding. Nature, 403(6772), 916–921. 10.1038/35002634

19. Gracia-Diaz, C., Zhou, Y., Yang, Q., Maroofian, R., Espana-Bonilla, P., Lee, C. H., Zhang, S., Padilla, N., Fueyo, R., Waxman, E. A., Lei, S., Otrimski, G., Li, D., Sheppard, S. E., Mark, P., Harr, M. H., Hakonarson, H., Rodan, L., Jackson, A., … Akizu, N. (2023). Gain and loss of function variants in EZH1 disrupt neurogenesis and cause dominant and recessive neurodevelopmental disorders. Nat Commun, 14(1), 4109. 10.1038/s41467-023-39645-5

20. Harris, H. K., Nakayama, T., Lai, J., Zhao, B., Argyrou, N., Gubbels, C. S., Soucy, A., Genetti, C. A., Suslovitch, V., Rodan, L. H., Tiller, G. E., Lesca, G., Gripp, K. W., Asadollahi, R., Hamosh, A., Applegate, C. D., Turnpenny, P. D., Simon, M. E. H., Volker-Touw, C. M. L., … Yu, T. W. (2021). Disruption of RFX family transcription factors causes autism, attention-deficit/hyperactivity disorder, intellectual disability, and dysregulated behavior. Genet Med, 23(6), 1028–1040. 10.1038/s41436-021-01114-z

21. Heinz, S., Benner, C., Spann, N., Bertolino, E., Lin, Y. C., Laslo, P., Cheng, J. X., Murre, C., Singh, H., & Glass, C. K. (2010). Simple combinations of lineage-determining transcription factors prime cis-regulatory elements required for macrophage and B cell identities. Mol Cell, 38(4), 576–589. 10.1016/j.molcel.2010.05.004

22. Hollestein, V., Poelmans, G., Forde, N. J., Beckmann, C. F., Ecker, C., Mann, C., Schafer, T., Moessnang, C., Baumeister, S., Banaschewski, T., Bourgeron, T., Loth, E., Dell’Acqua, F., Murphy, D. G. M., Puts, N. A., Tillmann, J., Charman, T., Jones, E. J. H., Mason, L., … Naaijen, J. (2023). Excitatory/inhibitory imbalance in autism: the role of glutamate and GABA gene-sets in symptoms and cortical brain structure. Transl Psychiatry, 13(1), 18. 10.1038/s41398-023-02317-5

23. Joung, J., Ma, S., Tay, T., Geiger-Schuller, K. R., Kirchgatterer, P. C., Verdine, V. K., Guo, B., Arias-Garcia, M. A., Allen, W. E., Singh, A., Kuksenko, O., Abudayyeh, O. O., Gootenberg, J. S., Fu, Z., Macrae, R. K., Buenrostro, J. D., Regev, A., & Zhang, F. (2023). A transcription factor atlas of directed differentiation. Cell, 186(1), 209–229 e226. 10.1016/j.cell.2022.11.026

24. Kaushik, G., & Zarbalis, K. S. (2016). Prenatal Neurogenesis in Autism Spectrum Disorders. Front Chem, 4, 12. 10.3389/fchem.2016.00012

25. Kaushik, K., Chapman, G., Prakasam, R., Batool, F., Saleh, M., Determan, J., Huettner, J. E., & Kroll, K. L. (2025). Requirements for the neurodevelopmental disorder-associated gene ZNF292 in human cortical interneuron development and function. Cell Rep, 44(5), 115597. 10.1016/j.celrep.2025.115597

26. Kechin, A., Boyarskikh, U., Kel, A., & Filipenko, M. (2017). cutPrimers: A New Tool for Accurate Cutting of Primers from Reads of Targeted Next Generation Sequencing. J Comput Biol, 24(11), 1138–1143. 10.1089/cmb.2017.0096

27. Kremitzki, C., Waligorski, J., Bachman, G., Ali, L. M., Bramley, J., Vakaki, M., Chandrasekaran, V., Patel, P., Mathur, D., Hime, P., Mitra, R., Milbrandt, J., & Buchser, W. (2025). Pathogenic morphological signatures of perturbations in mitochondrial-related genes revealed by pooled imaging assay. Npj Imaging, 3(1), 35. 10.1038/s44303-025-00097-9

28. Lai, J., Demirbas, D., Phillips, K., Zhao, B., Wallace, H., Seferian, M., Nakayama, T., Harris, H., Chatzipli, A., Lee, E. A., & Yu, T. W. (2025). Multi-omic analysis of the ciliogenic transcription factor RFX3 reveals a role in promoting activity-dependent responses via enhancing CREB binding in human neurons. bioRxiv. 10.1101/2025.02.27.640588

29. Liu, S., Li, D., Lyu, C., Gontarz, P. M., Miao, B., Madden, P. A. F., Wang, T., & Zhang, B. (2021). AIAP: A Quality Control and Integrative Analysis Package to Improve ATAC-seq Data Analysis. Genomics Proteomics Bioinformatics, 19(4), 641–651. 10.1016/j.gpb.2020.06.025

30. Love, M. I., Huber, W., & Anders, S. (2014). Moderated estimation of fold change and dispersion for RNA-seq data with DESeq2. Genome Biol, 15(12), 550. 10.1186/s13059-014-0550-8

31. Lui, J. H., Hansen, D. V., & Kriegstein, A. R. (2011). Development and evolution of the human neocortex. Cell, 146(1), 18–36. 10.1016/j.cell.2011.06.030

32. Lunn, S. (2022). Development and application of human cortical organoids: A 1q21.1 study Cardiff University].

33. Mariani, J., Coppola, G., Zhang, P., Abyzov, A., Provini, L., Tomasini, L., Amenduni, M., Szekely, A., Palejev, D., Wilson, M., Gerstein, M., Grigorenko, E. L., Chawarska, K., Pelphrey, K. A., Howe, J. R., & Vaccarino, F. M. (2015). FOXG1-Dependent Dysregulation of GABA/Glutamate Neuron Differentiation in Autism Spectrum Disorders. Cell, 162(2), 375–390. 10.1016/j.cell.2015.06.034

34. Meganathan, K., Lewis, E. M. A., Gontarz, P., Liu, S., Stanley, E. G., Elefanty, A. G., Huettner, J. E., Zhang, B., & Kroll, K. L. (2017). Regulatory networks specifying cortical interneurons from human embryonic stem cells reveal roles for CHD2 in interneuron development. Proc Natl Acad Sci U S A, 114(52), E11180–E11189. 10.1073/pnas.1712365115

35. Morotomi-Yano, K., Yano, K., Saito, H., Sun, Z., Iwama, A., & Miki, Y. (2002). Human regulatory factor X 4 (RFX4) is a testis-specific dimeric DNA-binding protein that cooperates with other human RFX members. J Biol Chem, 277(1), 836–842. 10.1074/jbc.M108638200

36. Neumann, S., Achiro, J. M., Watanabe, M., Bonnano, S. L., Deng, W., Wohlschlegel, J. A., & Martin, K. C. (2025). Cytoplasmic and nuclear protein interaction networks of the synapto-nuclear messenger CRTC1 in neurons reveal cooperative chromatin binding between CREB1 and CRTC1, MEF2C and RFX3. bioRxiv. 10.1101/2025.07.02.662820

37. Niu, W., Yu, S., Li, X., Wang, Z., Chen, R., Michalski, C., Jahangiri, A., Zohdy, Y., Chern, J. J., Whitworth, T. J., Wang, J., Xu, J., Zhou, Y., Qin, Z., Li, B., Gambello, M. J., Peng, J., & Wen, Z. (2024). Longitudinal multi-omics reveals pathogenic TSC2 variants disrupt developmental trajectories of human cortical organoids derived from Tuberous Sclerosis Complex. bioRxiv. 10.1101/2024.10.07.617121

38. Pampari, A., Shcherbina, A., Kvon, E. Z., Kosicki, M., Nair, S., Kundu, S., Kathiria, A. S., Risca, V. I., Kuningas, K., Alasoo, K., Greenleaf, W. J., Pennacchio, L. A., & Kundaje, A. (2025). ChromBPNet: bias factorized, base-resolution deep learning models of chromatin accessibility reveal cis-regulatory sequence syntax, transcription factor footprints and regulatory variants. bioRxiv. 10.1101/2024.12.25.630221

39. Patro, R., Duggal, G., Love, M. I., Irizarry, R. A., & Kingsford, C. (2017). Salmon provides fast and bias-aware quantification of transcript expression. Nat Methods, 14(4), 417–419. 10.1038/nmeth.4197

40. Pereira, J. D., Sansom, S. N., Smith, J., Dobenecker, M. W., Tarakhovsky, A., & Livesey, F. J. (2010). Ezh2, the histone methyltransferase of PRC2, regulates the balance between self-renewal and differentiation in the cerebral cortex. Proc Natl Acad Sci U S A, 107(36), 15957–15962. 10.1073/pnas.1002530107

41. Prakasam, R., Determan, J., Chapman, G., Narasimhan, M., Shen, R., Saleh, M., Kaushik, K., Gontarz, P., Meganathan, K., Hakim, B., Zhang, B., Huettner, J. E., & Kroll, K. L. (2025). Autism-and intellectual disability-associated MYT1L mutation alters human cortical interneuron differentiation, maturation, and physiology. Stem Cell Reports, 20(3), 102421. 10.1016/j.stemcr.2025.102421

42. Ramirez, F., Ryan, D. P., Gruning, B., Bhardwaj, V., Kilpert, F., Richter, A. S., Heyne, S., Dundar, F., & Manke, T. (2016). deepTools2: a next generation web server for deep-sequencing data analysis. Nucleic Acids Res, 44(W1), W160–165. 10.1093/nar/gkw257

43. Ross-Innes, C. S., Stark, R., Teschendorff, A. E., Holmes, K. A., Ali, H. R., Dunning, M. J., Brown, G. D., Gojis, O., Ellis, I. O., Green, A. R., Ali, S., Chin, S. F., Palmieri, C., Caldas, C., & Carroll, J. S. (2012). Differential oestrogen receptor binding is associated with clinical outcome in breast cancer. Nature, 481(7381), 389–393. 10.1038/nature10730

44. Schafer, S. T., Paquola, A. C. M., Stern, S., Gosselin, D., Ku, M., Pena, M., Kuret, T. J. M., Liyanage, M., Mansour, A. A., Jaeger, B. N., Marchetto, M. C., Glass, C. K., Mertens, J., & Gage, F. H. (2019). Pathological priming causes developmental gene network heterochronicity in autistic subject-derived neurons. Nat Neurosci, 22(2), 243–255. 10.1038/s41593-018-0295-x

45. Schneider, C. A., Rasband, W. S., & Eliceiri, K. W. (2012). NIH Image to ImageJ: 25 years of image analysis. Nat Methods, 9(7), 671–675. 10.1038/nmeth.2089

46. Sedykh, I., Keller, A. N., Yoon, B., Roberson, L., Moskvin, O. V., & Grinblat, Y. (2018). Zebrafish Rfx4 controls dorsal and ventral midline formation in the neural tube. Dev Dyn, 247(4), 650–659. 10.1002/dvdy.24613

47. Sentmanat, M. F., Peters, S. T., Florian, C. P., Connelly, J. P., & Pruett-Miller, S. M. (2018). A Survey of Validation Strategies for CRISPR-Cas9 Editing. Sci Rep, 8(1), 888. 10.1038/s41598-018-19441-8

48. Siegel-Ramsay, J. E., Romaniuk, L., Whalley, H. C., Roberts, N., Branigan, H., Stanfield, A. C., Lawrie, S. M., & Dauvermann, M. R. (2021). Glutamate and functional connectivity –support for the excitatory-inhibitory imbalance hypothesis in autism spectrum disorders. Psychiatry Res Neuroimaging, 313, 111302. 10.1016/j.pscychresns.2021.111302

49. Soneson, C., Love, M. I., & Robinson, M. D. (2015). Differential analyses for RNA-seq: transcript-level estimates improve gene-level inferences. F1000Res, 4, 1521. 10.12688/f1000research.7563.2

50. Stirling, D. R., Swain-Bowden, M. J., Lucas, A. M., Carpenter, A. E., Cimini, B. A., & Goodman, A. (2021). CellProfiler 4: improvements in speed, utility and usability. BMC Bioinformatics, 22(1), 433. 10.1186/s12859-021-04344-9

51. Vasan, L., Moffat, A., Mattar, P., & Schuurmans, C. (2025). Neocortical neurogenesis: a proneural gene perspective. FEBS J, 292(21), 5580–5610. 10.1111/febs.70158

52. Vazin, T., Ball, K. A., Lu, H., Park, H., Ataeijannati, Y., Head-Gordon, T., Poo, M. M., & Schaffer, D. V. (2014). Efficient derivation of cortical glutamatergic neurons from human pluripotent stem cells: a model system to study neurotoxicity in Alzheimer’s disease. Neurobiol Dis, 62, 62–72. 10.1016/j.nbd.2013.09.005

53. Villalba, A., Gotz, M., & Borrell, V. (2021). The regulation of cortical neurogenesis. Curr Top Dev Biol, 142, 1–66. 10.1016/bs.ctdb.2020.10.003

54. Vishnubalaji, R., Manikandan, M., Aldahmash, A., AlJarbou, A., Habous, M., Alhajeri, D., Almannie, R., Alfayez, M., Alajez, N. M., & Binsaleh, S. (2018). Whole genome mRNA expression profiling revealed multiple deregulated pathways in stromal vascular fraction from erectile dysfunction patients. Biosci Rep, 38(6). 10.1042/BSR20181015

55. Wang, L., Wang, C., Moriano, J. A., Chen, S., Zuo, G., Cebrian-Silla, A., Zhang, S., Mukhtar, T., Wang, S., Song, M., de Oliveira, L. G., Bi, Q., Augustin, J. J., Ge, X., Paredes, M. F., Huang, E. J., Alvarez-Buylla, A., Duan, X., Li, J., & Kriegstein, A. R. (2025). Molecular and cellular dynamics of the developing human neocortex. Nature. 10.1038/s41586-024-08351-7

56. Wang, X., Spandidos, A., Wang, H., & Seed, B. (2012). PrimerBank: a PCR primer database for quantitative gene expression analysis, 2012 update. Nucleic Acids Res, 40(Database issue), D1144–1149. 10.1093/nar/gkr1013

57. Xu, P., Morrison, J. P., Foley, J. F., Stumpo, D. J., Ward, T., Zeldin, D. C., & Blackshear, P. J. (2018). Conditional ablation of the RFX4 isoform 1 transcription factor: Allele dosage effects on brain phenotype. PLoS One, 13(1), e0190561. 10.1371/journal.pone.0190561

58. Yang, H., Hao, D., Liu, C., Huang, D., Chen, B., Fan, H., Liu, C., Zhang, L., Zhang, Q., An, J., & Zhao, J. (2019). Generation of functional dopaminergic neurons from human spermatogonial stem cells to rescue parkinsonian phenotypes. Stem Cell Res Ther, 10(1), 195. 10.1186/s13287-019-1294-x

59. Yu, G., Wang, L. G., & He, Q. Y. (2015). ChIPseeker: an R/Bioconductor package for ChIP peak annotation, comparison and visualization. Bioinformatics, 31(14), 2382–2383. 10.1093/bioinformatics/btv145

60. Zhang, D., Stumpo, D. J., Graves, J. P., DeGraff, L. M., Grissom, S. F., Collins, J. B., Li, L., Zeldin, D. C., & Blackshear, P. J. (2006). Identification of potential target genes for RFX4_v3, a transcription factor critical for brain development. J Neurochem, 98(3), 860–875. 10.1111/j.1471-4159.2006.03930.x

61. Zhang, D., Zeldin, D. C., & Blackshear, P. J. (2007). Regulatory factor X4 variant 3: a transcription factor involved in brain development and disease. J Neurosci Res, 85(16), 3515–3522. 10.1002/jnr.21356

