## Supplemental Data Table Titles and Supplemental Figures for "Intellectual disability risk gene RFX4 regulates cortical neurogenesis by restraining neuronal differentiation"

**Supplemental Data Legends**

**Data S1.** Germ layer and lineage-specific marker CT values and relative abundance as measured by RT-qPCR at day 12 of V- and D-NPC specification

**Data S2.** Differential gene expression (DEG) analysis in RFX4 HOM versus isogenic control NPCs

**Data S3.** Gene ontology enrichment analysis results for NPC DEGs

**Data S4.** Gene ontology enrichment analysis for endogenous RFX4 binding sites associated with the nearest transcription start site or RFX4 DEGs in D-NPCs and V-NPCs

**Data S5.** Differential DNA binding of RFX4 in D-NPCs and Finemohits summary of enriched sequences, as generated by ChromBPNet for D-NPCs

**Data S6.** Transcription factor binding site enrichment under endogenous RFX4 binding sites associated with D-NPC RFX4 HOM DEGs

**Data S7.** Differentially bound RFX3 binding sites identified in RFX4 HOM versus control or in RFX3 HOM versus isogenic control D-NPC comparisons

**Data S8.** Gene ontology enrichment analysis results for genes associated with endogenous RFX3 binding sites (based on the nearest transcription start site), RFX3 binding sites associated with RFX3 HOM DEGs or with shared RFX3 and RFX4 HOM DEGs

**Data S9.** Transcription factor binding site enrichment under RFX3 and RFX4 co-bound sites associated with RFX4 HOM D-NPC DEGs

**Data S10.** DEG analysis in RFX3 HOM versus isogenic control D-NPCs

**Data S11.** Differential gene expression analysis of RFX4 HOM versus isogenic control cEXs and D-NPCs

**Data S12.** Gene ontology enrichment analysis results of RFX4 HOM cEX and shared cEX and D-NPC DEGs

**Data S13.** Gene ontology enrichment analysis for genes (based upon the nearest transcription start site) associated with RFX3-bound sites in cEXs

**Data S14.** Differential DNA binding of RFX3 in RFX3-HOM and RFX4-HOM versus isogenic control cEXs

**Data S15.** Transcription factor binding site enrichment RFX3 binding sites that were significantly lost in RFX4-HOM cEXs

**Data S16.** Differential RFX4 binding in R79C-HOM and R79C-HET versus isogenic control D-NPCs and differential RFX3 binding in R79C-HOM D-NPCs

**Data S17.** Gene ontology enrichment analysis results for R79C-HOM and/or RFX4 HOM DEGs and for genes also associated with differential RFX4 binding in R79C D-NPCs

**Data S18.** Differential gene expression analysis of R79C-HOM versus isogenic control D-NPCs

**Data S19.** Detailed information on genetic backgrounds of hPSC models used in this study

**Data S20.** Individual data points and calculations for neurosphere outgrowth and n values for organoid immunocytochemistry quantifications

**Data S21.** Raw and unmodified western blot images

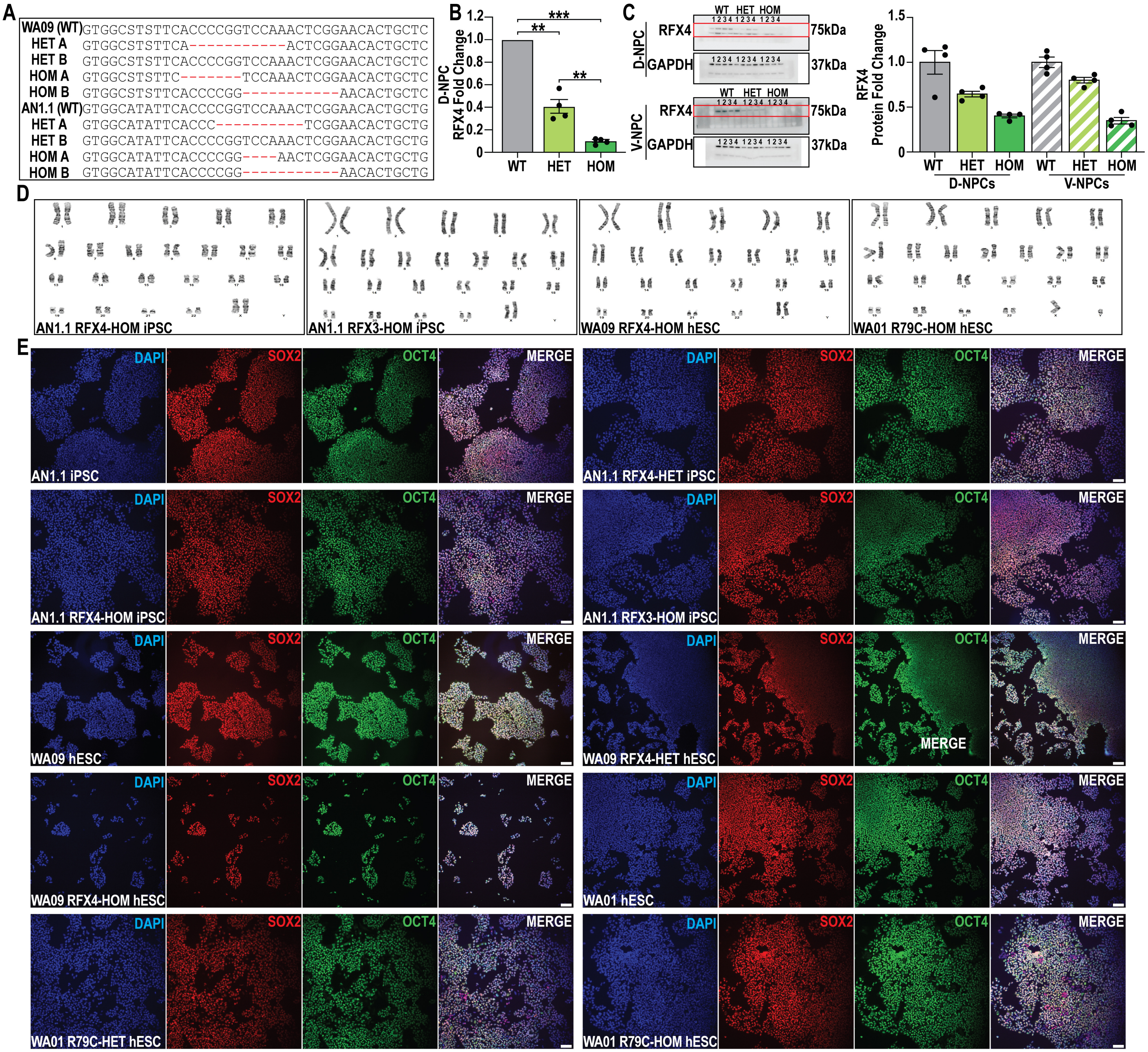


**Figure S1: Human pluripotent stem cell (hPSC) and RFX4 loss of function model validation.**

**(A)** Schematic representations of mutations introduced to generate heterozygous (HET) and homozygous (HOM) RFX4 LOF models. **(B)** RT-qPCR assessing the RFX4 mRNA expression levels in wildtype (WT), HET, and HOM D-NPCs. **(C)** Western blot image and quantification of RFX4 protein levels in WT, HET and HOM V- and D-NPCs relative to the GAPDH loading control. **(D)** Karyotyping results for cell lines indicated following CRISPR-Cas9 gene editing of induced pluripotent stem cell (iPSC) and human embryonic stem cell (hESC) lines. **(E)** Representative images of SOX2+ (red) and OCT4+ (green) iPSC and hESC models, validating that hPSC lines used in this study express pluripotency markers. **(B)** Data was analyzed by one-way ANOVA (n=4). Scale bar = 100 µm.

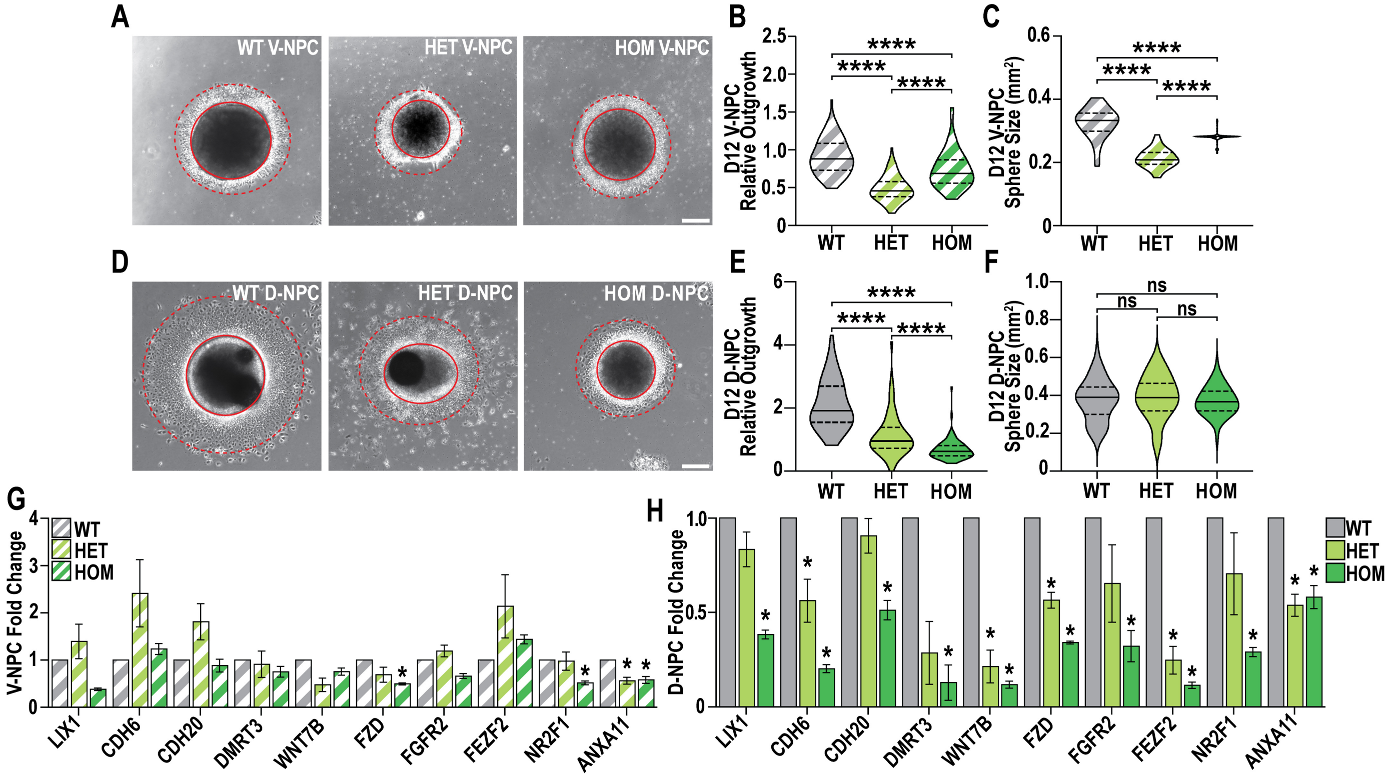


**Figure S2: RFX4 LOF NPC spheres exhibit decreased cell outgrowth and early diminished developmental gene expression.** **(A-F)** Quantification of **(A-C)** V-NPC and **(D-F)** D-NPC neurosphere outgrowth measuring **(B,E)** neurosphere body outgowth (marked by dotted red circle) and **(C,F)** neurosphere size (marked by solid red circle) after 12 days (D12) of differentiation. **(G-H)** RT-qPCR for DEGs downregulated in RFX4 HOM **(G)** V-NPCs or **(H)** D-NPCs across the WT, HET and HOM models. Data are represented as **(B,C,E,F)** distributions with quartiles (dashed lines) and means (solid line) or **(G-H)** mean fold change +/- SEM relative to GAPDH endogenous control and were analyzed by **(B,C,E,F)** one-way ANOVA (n=4) or **(G-H)** Student's t-test (n=4); ns = not significant, *p<0.05 and ****p< 0.0001. **(A,D)** Scale bar = 100 µm.

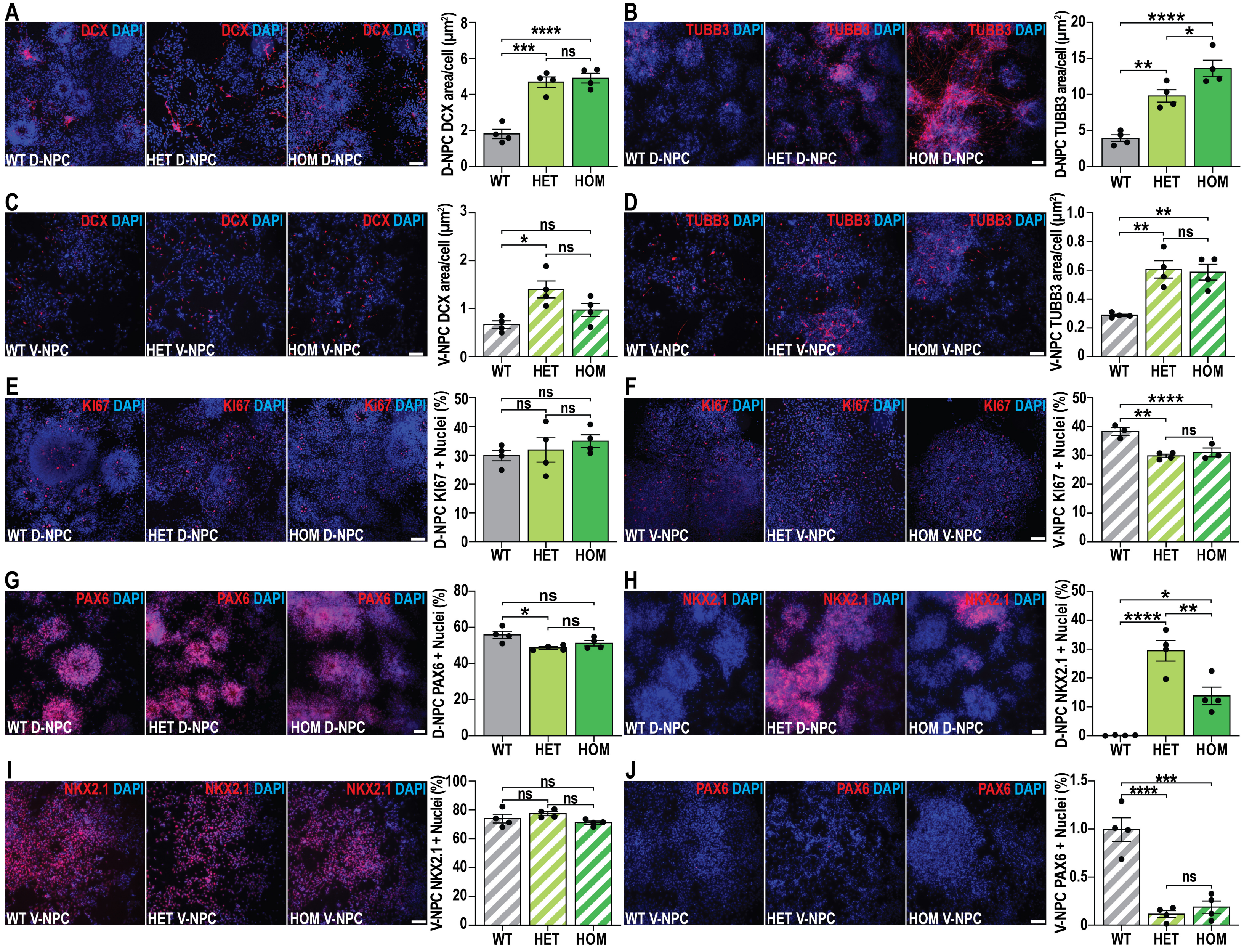


**Figure S3: RFX4 LOF alters regional neural specification and causes lineage-specific effects on proliferation.**
**(A-D)** Representative images and quantification of **(A,C)** DCX+ or **(B,D)** TUBB3+ area relative to total DAPI in WT, HET and HOM **(A-B)** D-NPCs and **(C-D)** V-NPCs. **(E-F)** Representative images and the quantification of KI67+ relative to total DAPI immunopositive nuclei in **(E)** D-NPCs or **(F)** V-NPCs. **(G-J)** Representation images and quantification of **(G,J)** PAX6+ and **(H-I)** NKX2.1+ relative to total DAPI in **(G-H)** D-NPCs and **(I-J)** V-NPCs. Data are represented as **(A-J)** mean +/- SEM and was analyzed by **(A-J)** one-way ANOVA (n=3-4); ns = not significant, *p<0.05, **p<0.01, ***p<0.001, and ****p< 0.0001, scale bars = 50 µm.

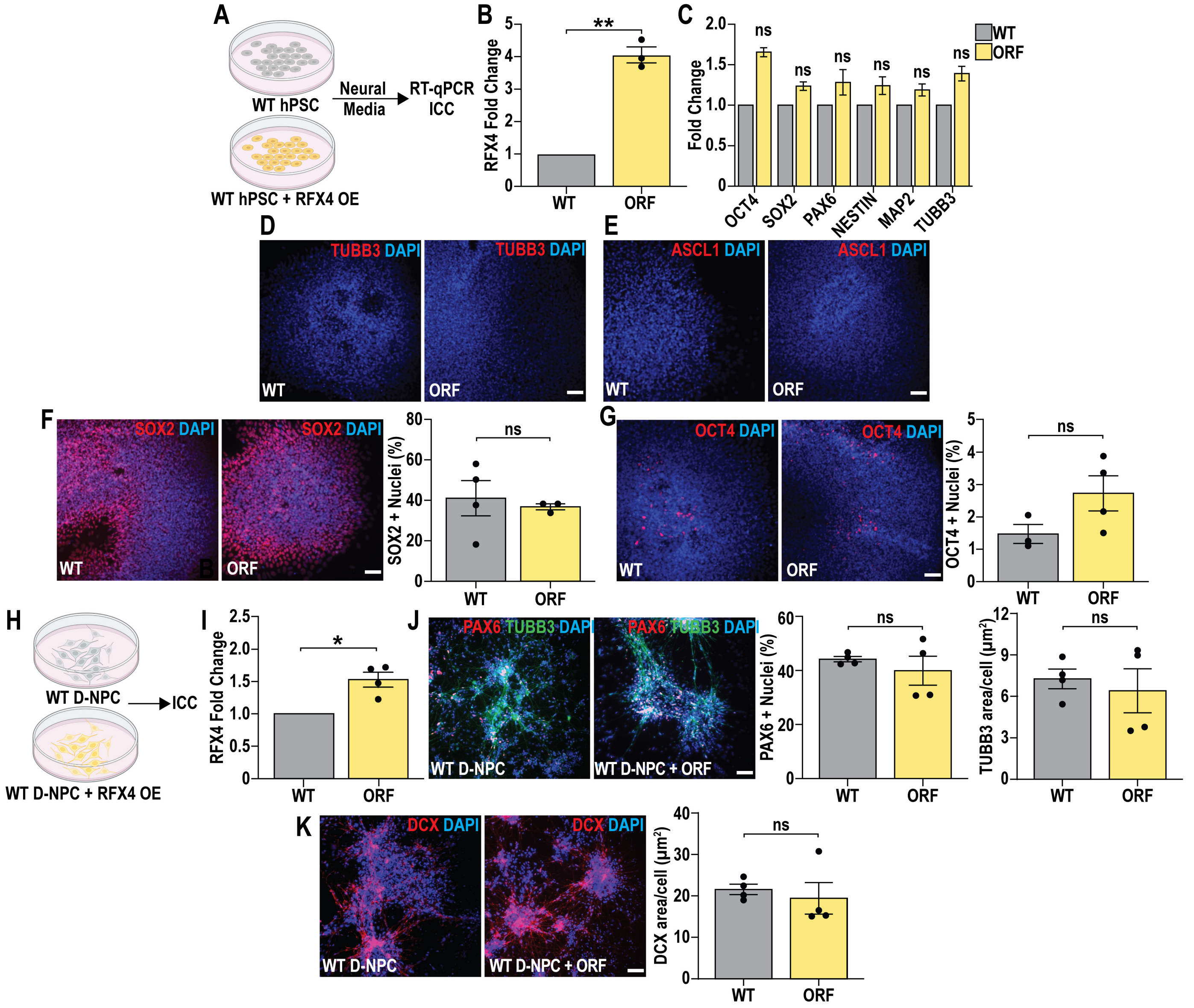


**Figure S4: RFX4 overexpression is insufficient to induce neuronal differentiation.**

**(A)** Schematic representation of RFX4 overexpression (OE) using an RFX4 open-reading frame (ORF) containing expression construct with downstream experimental analysis by RT-qPCR (mRNA) and immunocytochemistry (ICC). **(B-C)** RT-qPCR for the expression of **(B)** RFX4, **(C)** pluripotency and neuronal markers in hPSCs with or without RFX4 ORF OE. **(D-G)** Representative images and quantification of **(D)** TUBB3+, **(E)** ASCL1+, **(F)** SOX2+, and **(G)** OCT4+ cells relative to total DAPI in hPSCs with or without RFX4 ORF OE. **(H)** Schematic representation of RFX4 OE in D-NPCs and **(I)** confirmation of RFX4 OE by RT-qPCR. **(J)** Representative images and quantification of PAX6+ and TUBB3+ area relative to toal DAPI in WT and RFX4 OE D-NPCs. **(K)** Representative images and quantification of the DCX+ area in WT and RFX4 OE D-NPCs. Data are represented as mean +/- SEM, with fold change calculated relative to **(B-C)** GAPDH **(I)** and RPL30 loading controls, with **(F,G,J,K)** showing mean +/- SEM. All data was analyzed by Student's t-test (n≥3); ns = not significant and **p<0.01, **(D-G,J,K)** scale bars = 50 µm.

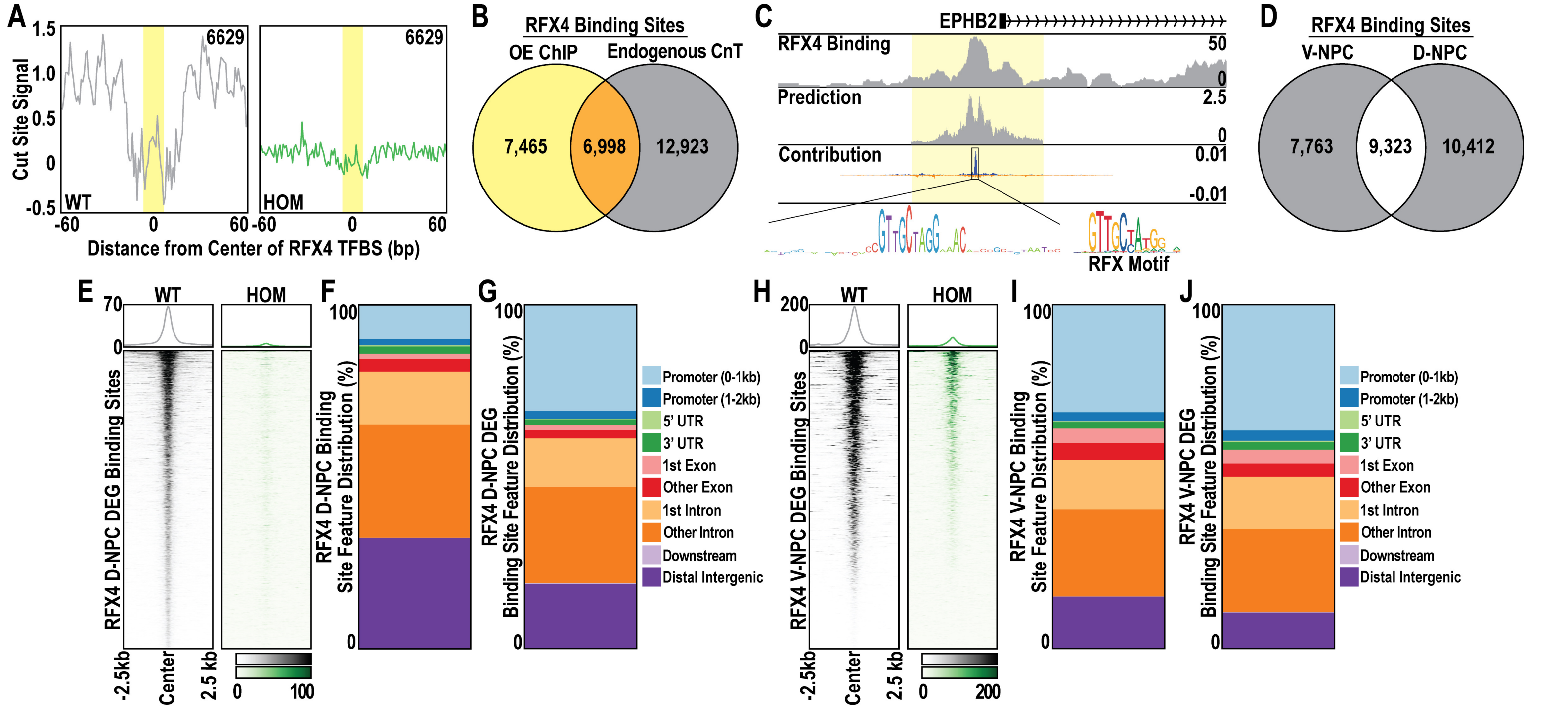


**Figure S5: Characterizing endogenous RFX4 genome-wide occupancy.**
**(A)** Cut site enrichment was defined by CUT&Tag analysis of RFX4 genome-wide binding, by comparing RFX4 bound sites containing an RFX4 TFBS in WT versus RFX4 HOM D-NPCs; these data highlighted reduction of cutting activity associated with sites of RFX occupancy and an RFX TFBS specifically in the WT D-NPC control; yellow highlight. These were lost in the HOM model. **(B)** Venn diagram showing overlap between previously identified RFX4 binding sites generated using ChIP for an overexpressed, epitope tagged form of RFX4 (Joung et al., 2023), with comparison with our endogenous RFX4 D-NPC genome-wide occupancy data. **(C)** Browser track example from BPNet highlights the RFX motif present at an RFX4 bound site in D-NPCs. **(D)** Venn diagram showing overlap between endogenous RFX4 binding sites, defined by CUT&Tag analysis in V- and D-NPCs. **(E)** Heatmap of RFX4 bound sites associated with RFX4 DEGs in D-NPCs. **(F-G)** Feature distribution of **(F)** all D-NPC RFX4 binding sites and **(G)** RFX4 D-NPC binding sites associated with DEGs. **(H)** Heatmap of RFX4 bound sites associated with RFX4 DEGs in V-NPCs. **(I-J)** Feature distribution of **(I)** all V-NPC RFX4 binding sites and **(J)** RFX4 V-NPC binding sites associated with DEGs.

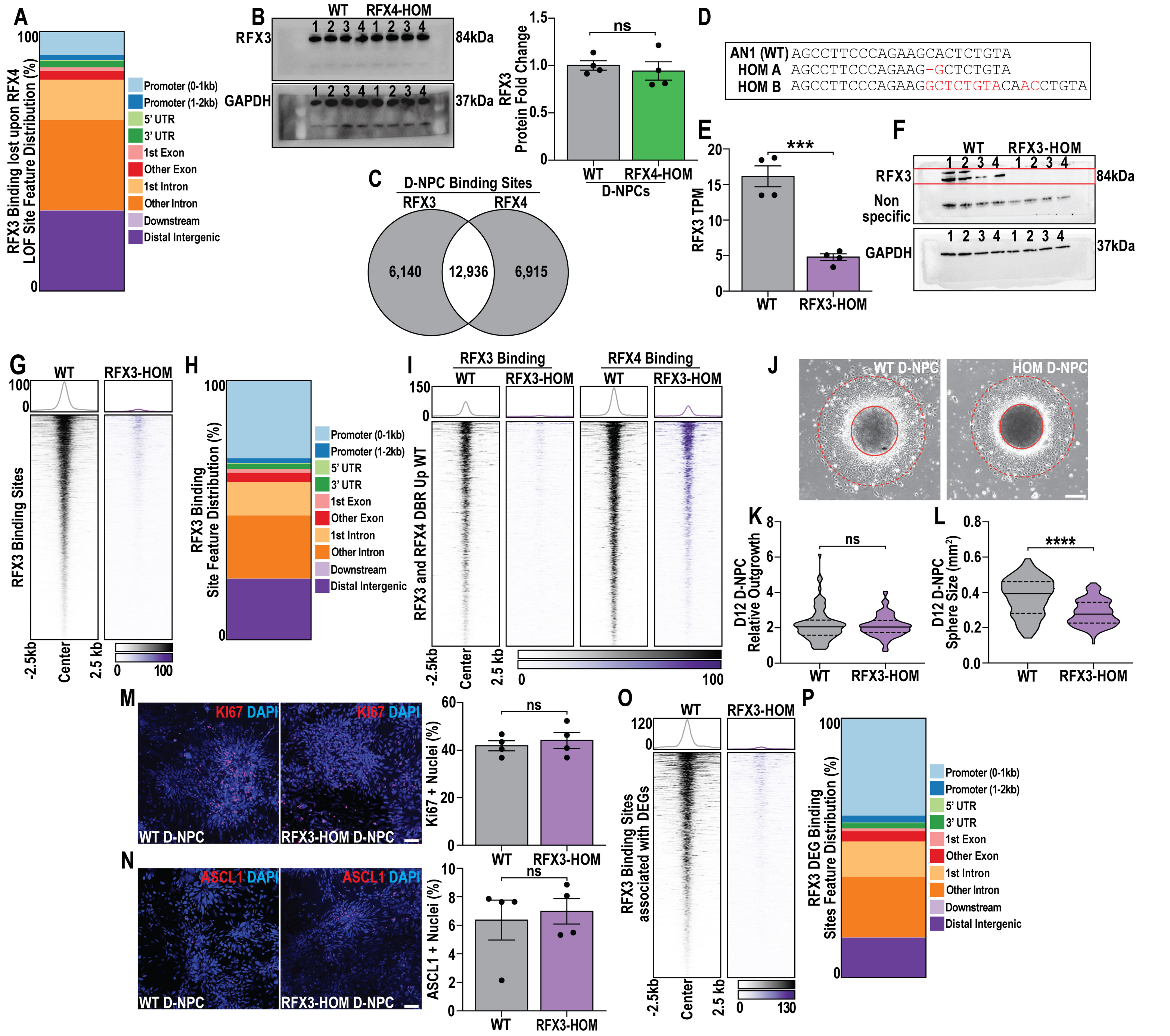


**Figure S6: RFX3 LOF produce phenotypes distinct from RFX4 LOF.**

**(A)** Feature distribution of RFX3 binding sites that were significantly lost upon RFX4 LOF. **(B)** Western blot image and quantification of RFX3 and GAPDH protein expression in WT and RFX4-HOM D-NPCs. **(C)** Venn diagram showing overlap between endogenous RFX3 and RFX4 binding sites, defined by CUT&Tag analysis in D-NPCs **(D)** Schematic representations of mutations introduced to generate homozygous (HOM) RFX3 LOF models. **(E)** Quantification of RFX3 mRNA expression in WT and RFX3-HOM D-NPCs **(F)** Western blot image of RFX3 and GAPDH protein expression in WT and RFX3-HOM D-NPCs. **(G)** Heatmap of RFX3 binding sites lost in RFX3-HOM D-NPCs. **(H)** Feature distribution of RFX3 binding sites lost upon RFX3 LOF. **(I)** Heatmap of RFX3 and RFX4 binding at co-bound sites which lose RFX3 and RFX4 binding in RFX3-HOM D-NPCs. **(J-L)** Quantification of **(K)** D-NPC neurosphere outgrowth (marked by dotted red circle) and **(L)** neurosphere size (marked by solid red circle) after 12 days of differentiation. **(M-N)** Representative images and quantification of the **(M)** KI67+ and **(N)** ASCL1+ cells relative to total DAPI in WT and RFX3-HOM D-NPCs. **(O)** Heatmap of RFX3 binding sites associated with RFX3 HOM D-NPC DEGs. **(P)** Feature distribution of RFX3 binding sites associated with RFX3 DEGs. Data are represented as **(B)** Fold change relative to GAPDH, **(E,M,N)** mean +/- SEM or as **(K-L)** data distributions with quartiles (dashed lines) and mean (solid line) marked and data was analyzed by **(E)** differential gene expression analysis (n=4) or **(B,K-N)** Student's t-test (n = 4); ns = not significant, ***p<0.001, and ****p<0.0001, scale bars = **(J)** 100 µm and **(M,N)** 50 µm.

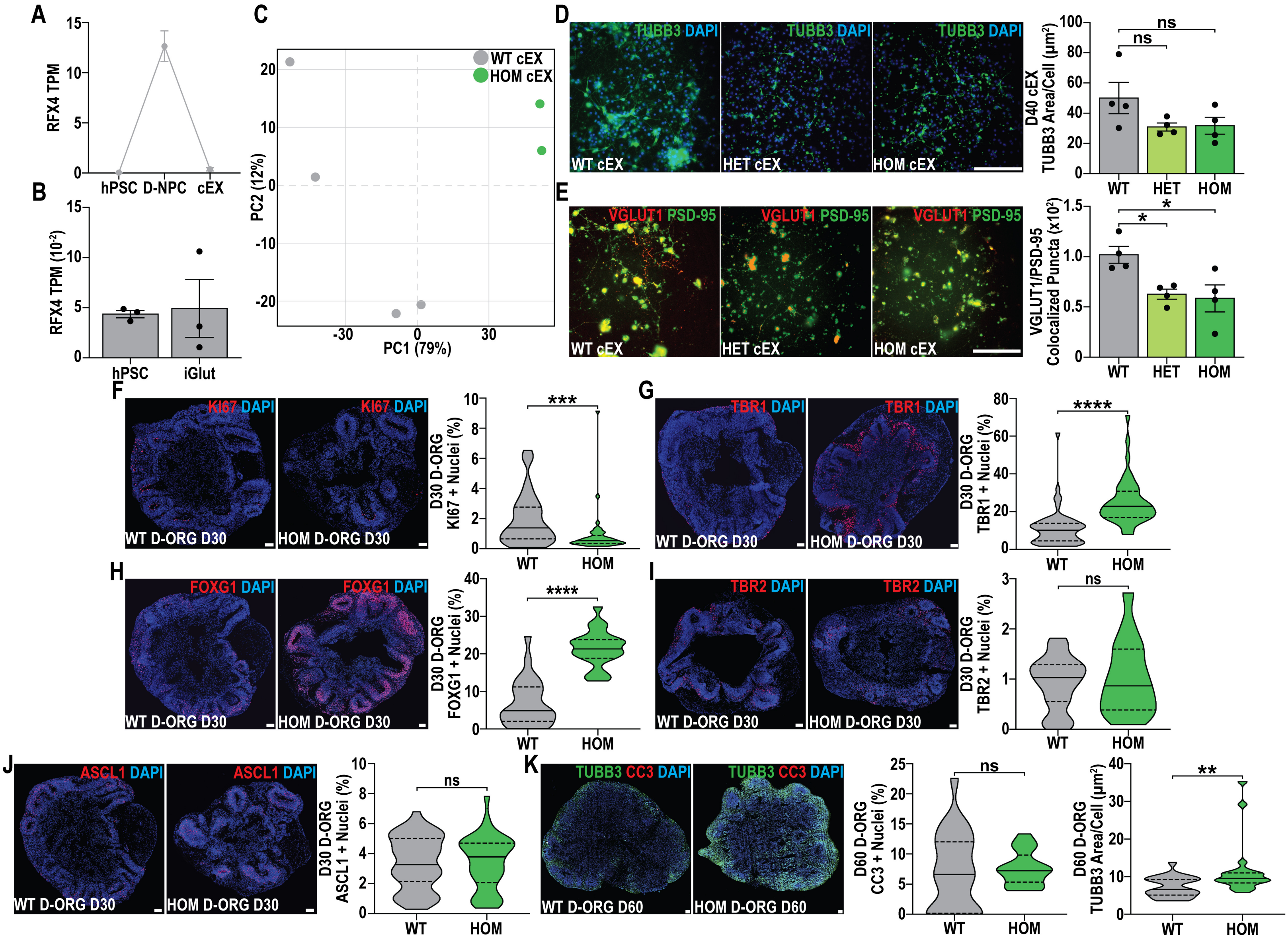


**Figure S7: RFX4 LOF also causes precocious neurogenesis and altered regional specification in D-ORGs.**

**(A-B)** Changes in the expression of RFX4 during **(A)** cEX or **(B)** iGlut differentiation. **(C)** PCA plot highlighting transcriptomic variation between WT (grey circles, n=4) and HOM (green circles, n=2) cEXs. **(D)** Representative images and quantification of TUBB3+ area relative to total DAPI in WT, HET, and HOM day 40 cEXs. **(E)** Representative images and quantification of VGLUT+ and PSD-95+ colocalized puncta in WT, HET, and HOM day 50 cEXs. **(F-J)** Representative images and quantification of the **(F)** KI67+, **(G)** TBR1+, **(H)** FOXG1+, **(I)** TBR2+, and **(J)** ASCL1+ cells relative to total DAPI in WT and HOM day 30 (D30) D-ORGs. **(K)** Representative images and quantification of the CC3+ and TUBB3+ area relative to total DAPI in WT and HOM day 60 (D60) D-ORGs. Data are represented as **(A,B,D,E)** mean +/- SEM and **(F-K)** data distributions with quartiles (dashed lines) and median (solid line) marked. Data was analyzed by **(A,B)** differential gene expression analysis (n≥3), **(D,E)** one-way ANOVA (n=4), or **(F-K)** Mann-Whitney test (for n values, see Data S20); ns = not significant, *p<0.05, **p<0.01, ***p<0.001, and ****p<0.0001, scale bars = 100 µm.

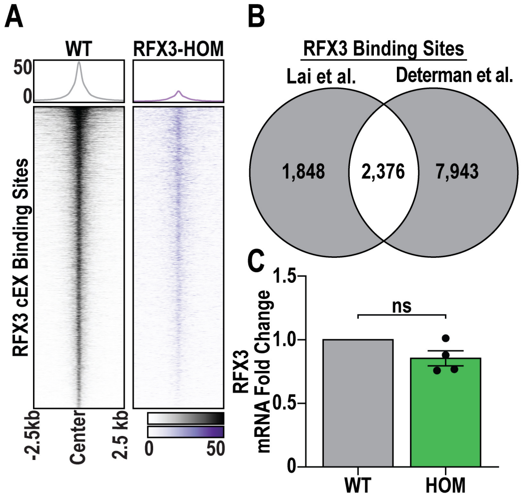


**Figure S8: Characterization of endogenous RFX3 genome-wide binding in cEXs.**

**(A)** Heatmap of RFX3 genome-wide binding in WT and RFX3-HOM cEXs, highlighting an almost complete loss of RFX3 binding upon RFX3 LOF. **(B)** Venn diagram shows overlap between previously identified RFX3 binding sites generated by using CUT&RUN in iGluts (Lai et al., 2025) and by CUT&Tag in cEXs (our work here), highlighting a set of shared sites. **(C)** RT-qPCR of RFX3 mRNA levels in WT and RFX4-HOM cEXs. Data are represented as mean +/- SEM with fold change calculated relative to RPL30. ns = not significant.

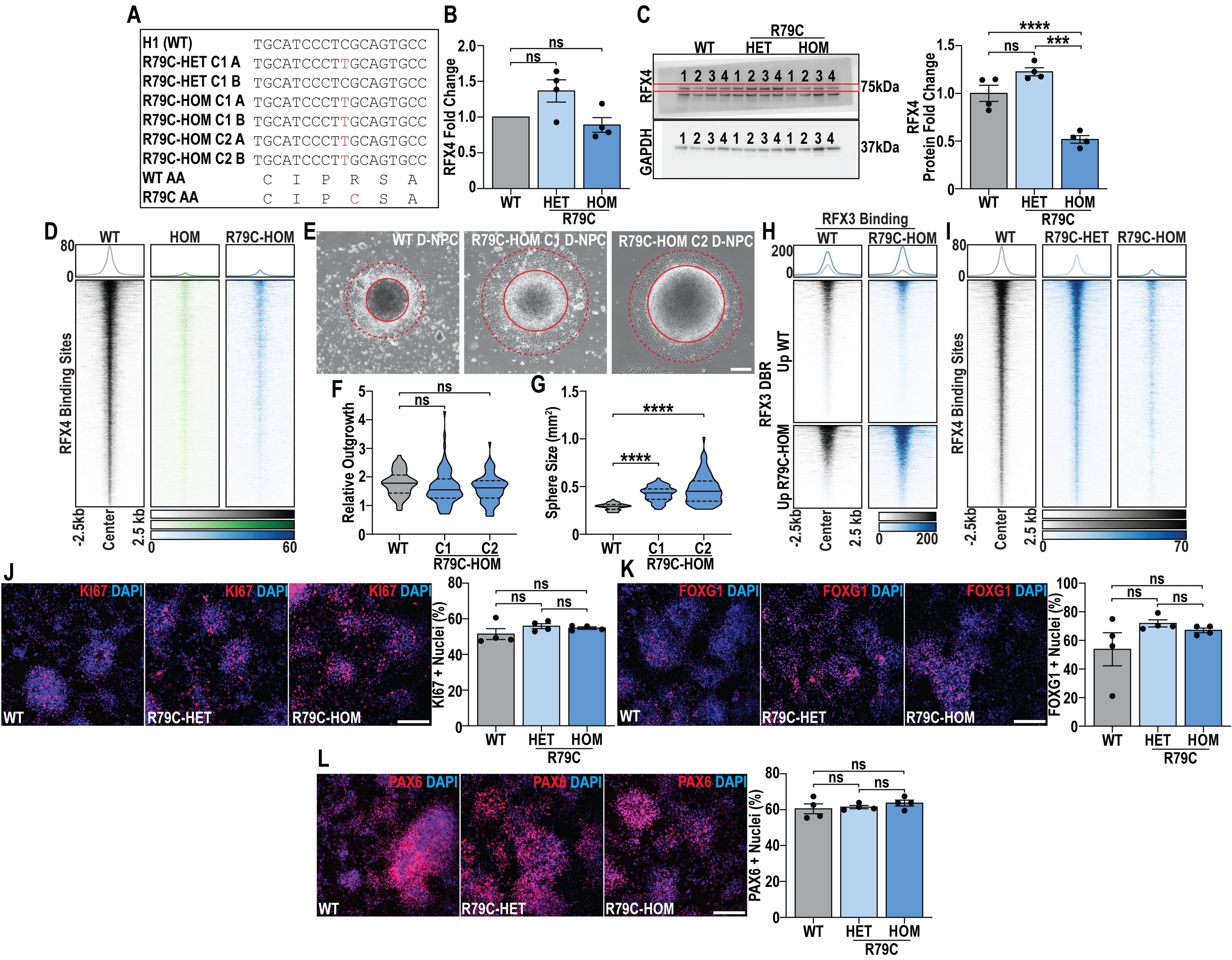

**Figure S9: p.R79C pathogenic mutation causes unique neurodevelopmental phenotypes.**
**(A)** Schematic representations of mutations introduced to generate heterozygous (R79C-HET) and homozygous (R79C-HOM) RFX4 pathogenic mutant models. **(B)** RT-qPCR for RFX4 mRNA expression in WT, R79C-HET, and R79C-HOM D-NPCs. **(C)** Western blot image and quantification of RFX4 protein levels relative to GAPDH in WT, R79C-HET, and R79C-HOM D-NPCs. **(D)** Heatmap of RFX4 bound peaks across WT, RFX4 HOM and RFX4 R79C-HOM D-NPCs. **(E-G)** Quantification of **(E)** D-NPC neurosphere outgrowth, measuring **(F)** neurosphere outgrowth (marked by dotted red circle) and **(G)** neurosphere size (marked by solid red circle) after 12 days of differentiation. **(H)** Heatmap of RFX3 differentially bound regions (DBRs) in WT and R79C-HOM D-NPCs. **(I)** Heatmap of RFX4 binding sites in WT, R79C-HET, and R79C-HOM D-NPCs. **(J-L)** Quantification and representative images of the **(J)** KI67+, **(K)** FOXG1+, and **(L)** PAX6+ cells relative to toal DAPI in WT, R79C-HET, and R79C-HOM D-NPCs. Data are represented as **(B,C,J,K,L)** mean +/- SEM, with fold changes calculated relative to the **(B)** RPL30 or **(C)** GAPDH endogenous control, or as **(F,G)** data distributions with quartiles (dashed lines) and mean (solid line) marked. Data was analyzed by **(B,C,F,G,J-L)** one-way ANOVA (n=4); ns = not significant, ***p<0.001, and ****p<0.0001, scale bar = **(E)** 100 µm or **(J-L)** 50 µm.
