## Supplementary figures and images for "Intellectual disability risk gene RFX4 regulates cortical neurogenesis by restraining neuronal differentiation"

### Figure S1C_D-NPCs_RFX4_GAPDH_WT_HET_HOM.tif

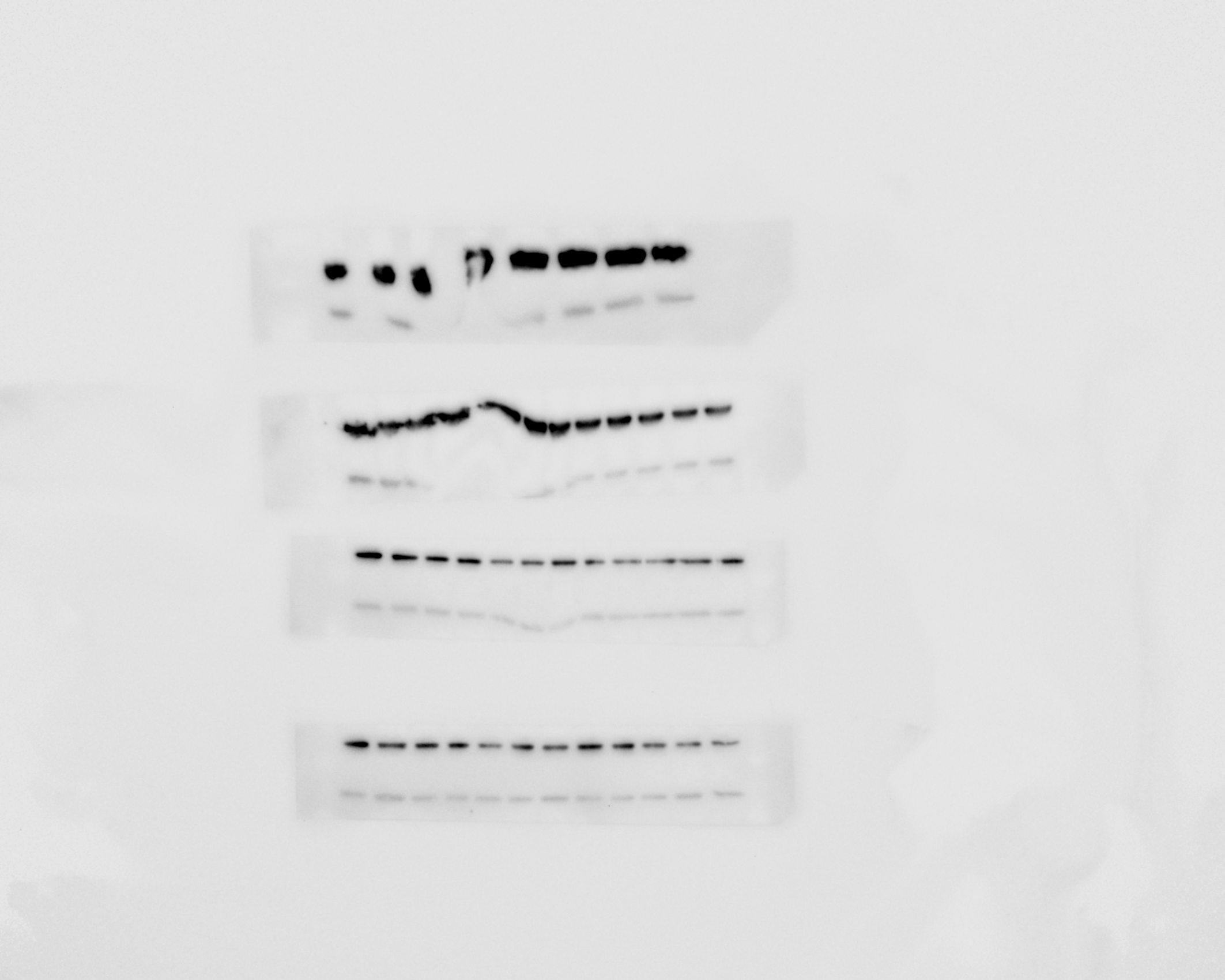

### Figure S1C_D-NPCs_RFX4_WT_HET_HOM.tif

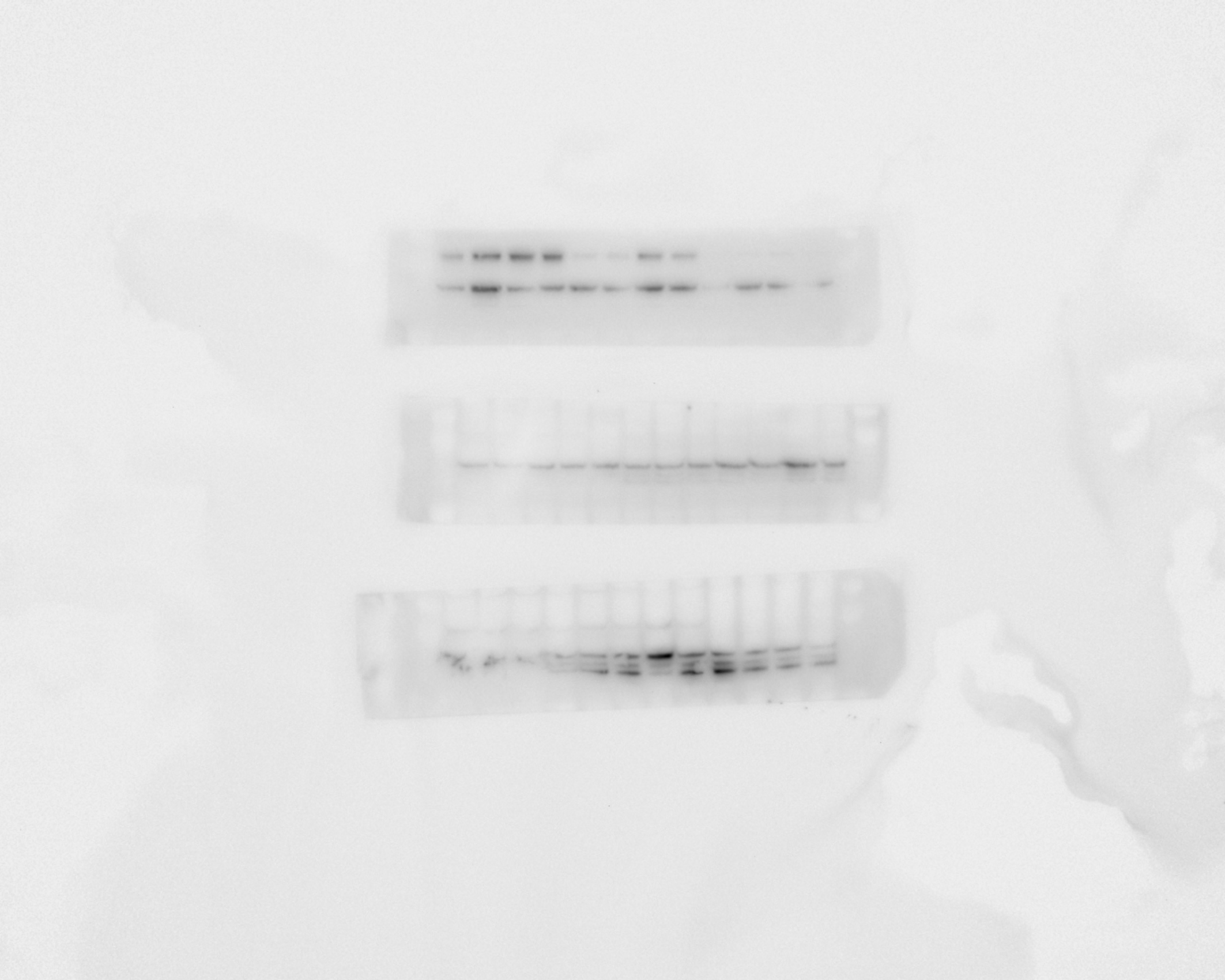

### Figure S1C_V-NPCs_RFX4_WT_HET_HOM.tif

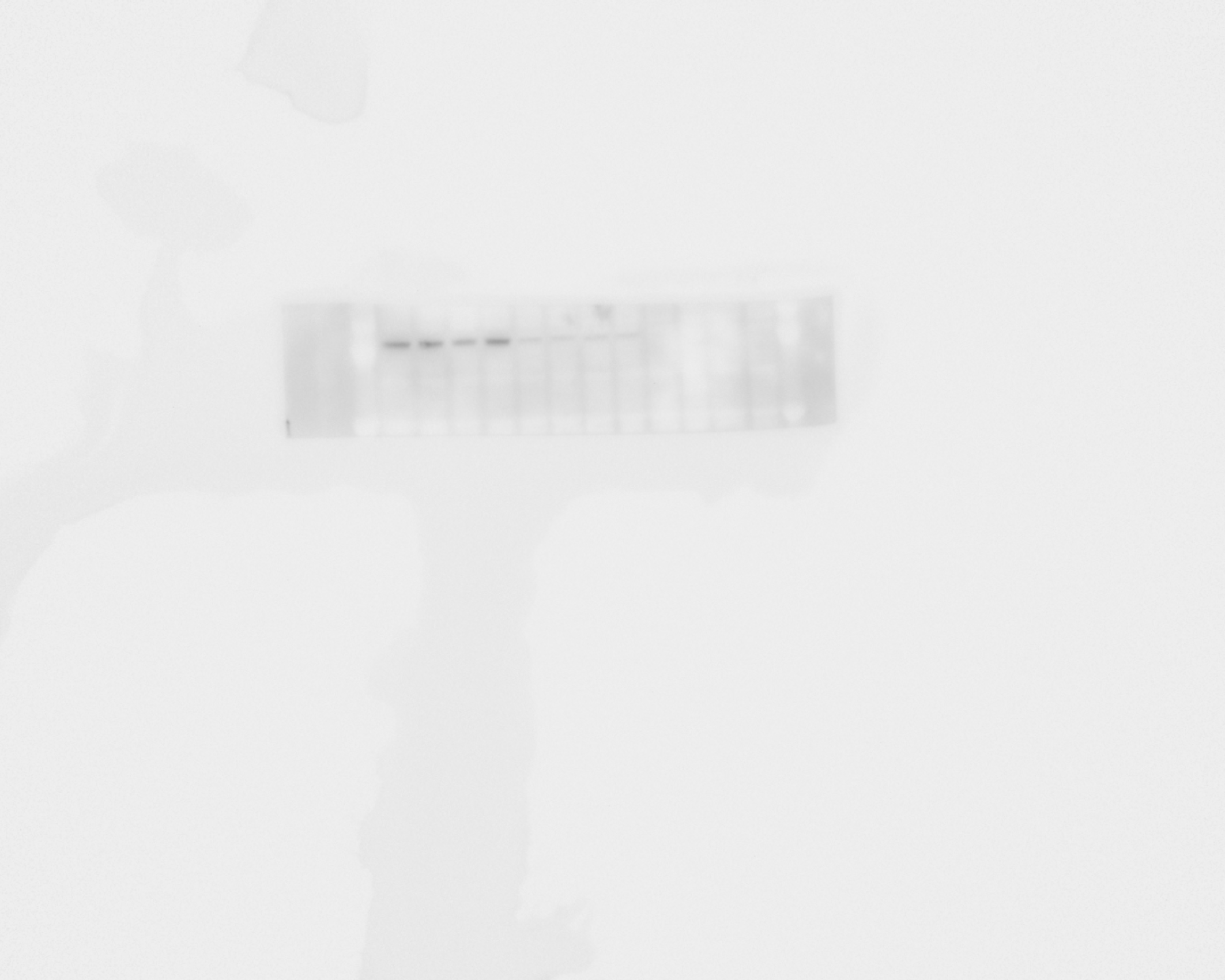

### Figure S6B_D-NPCs_RFX3_GAPDH_WT_RFX4-HOM.tif

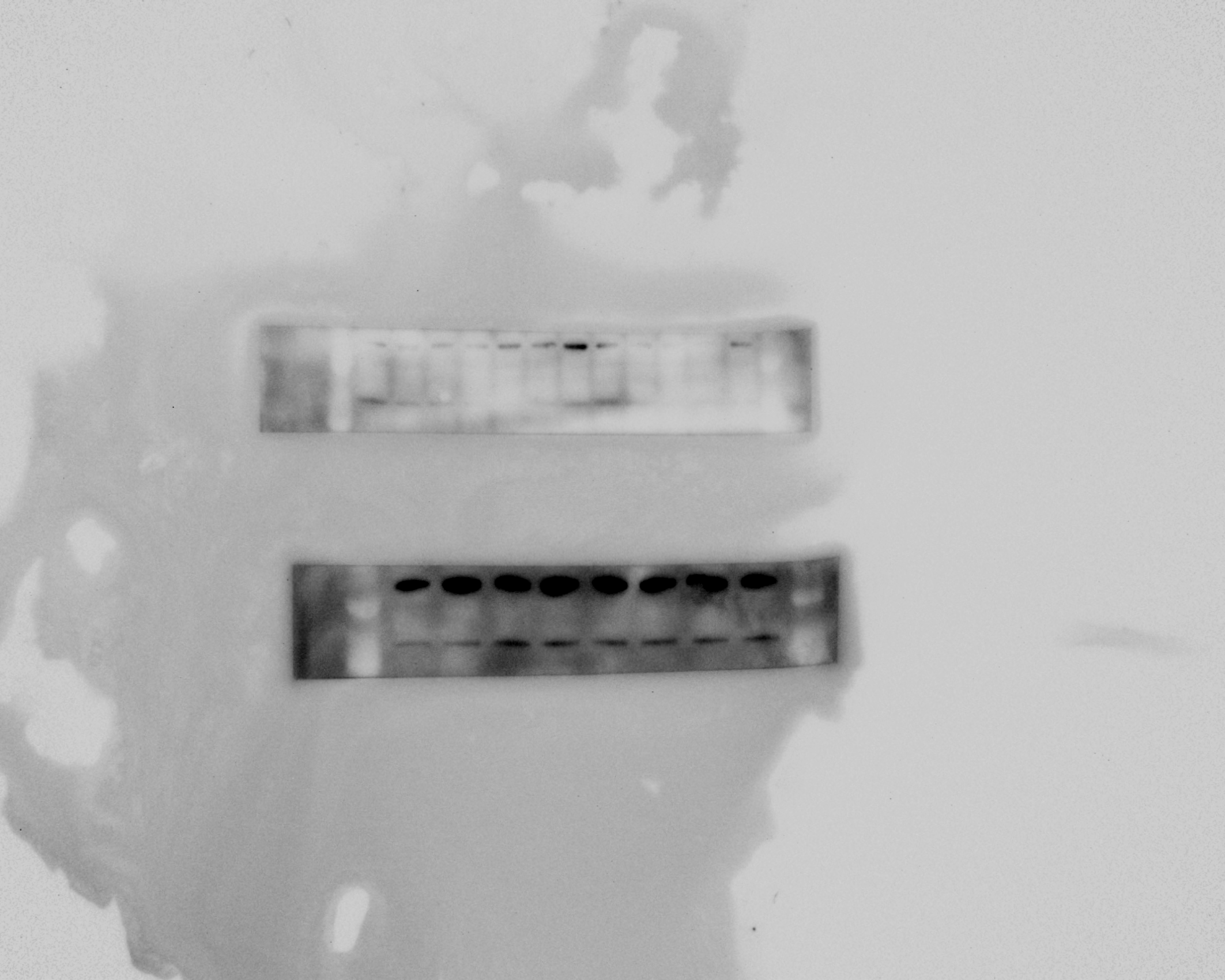

### Figure S6B_D-NPCs_RFX3_WT_RFX4-HOM.tif

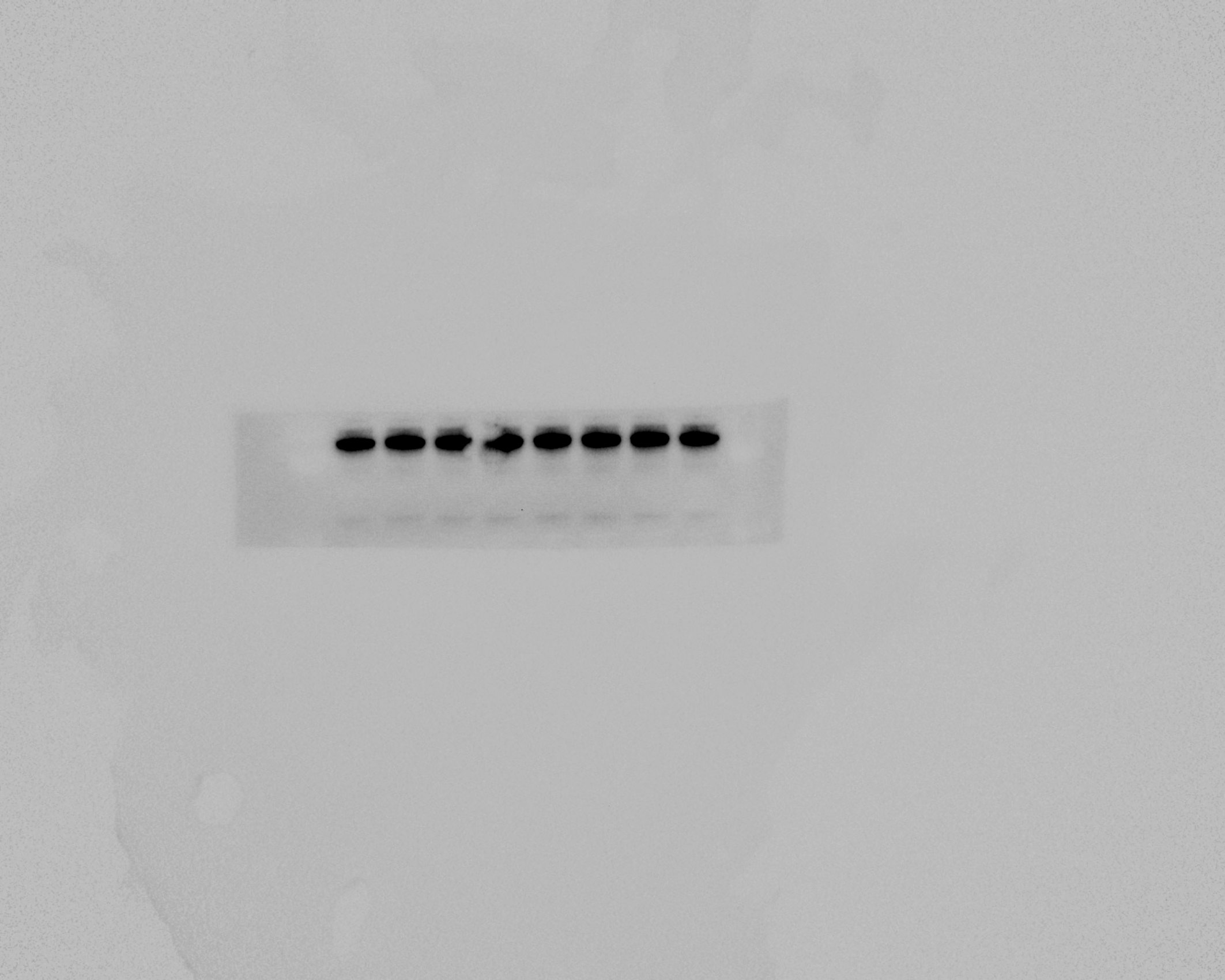

### Figure S6F_D-NPCs_RFX3_GAPDH_WT_RFX3-HOM.tif

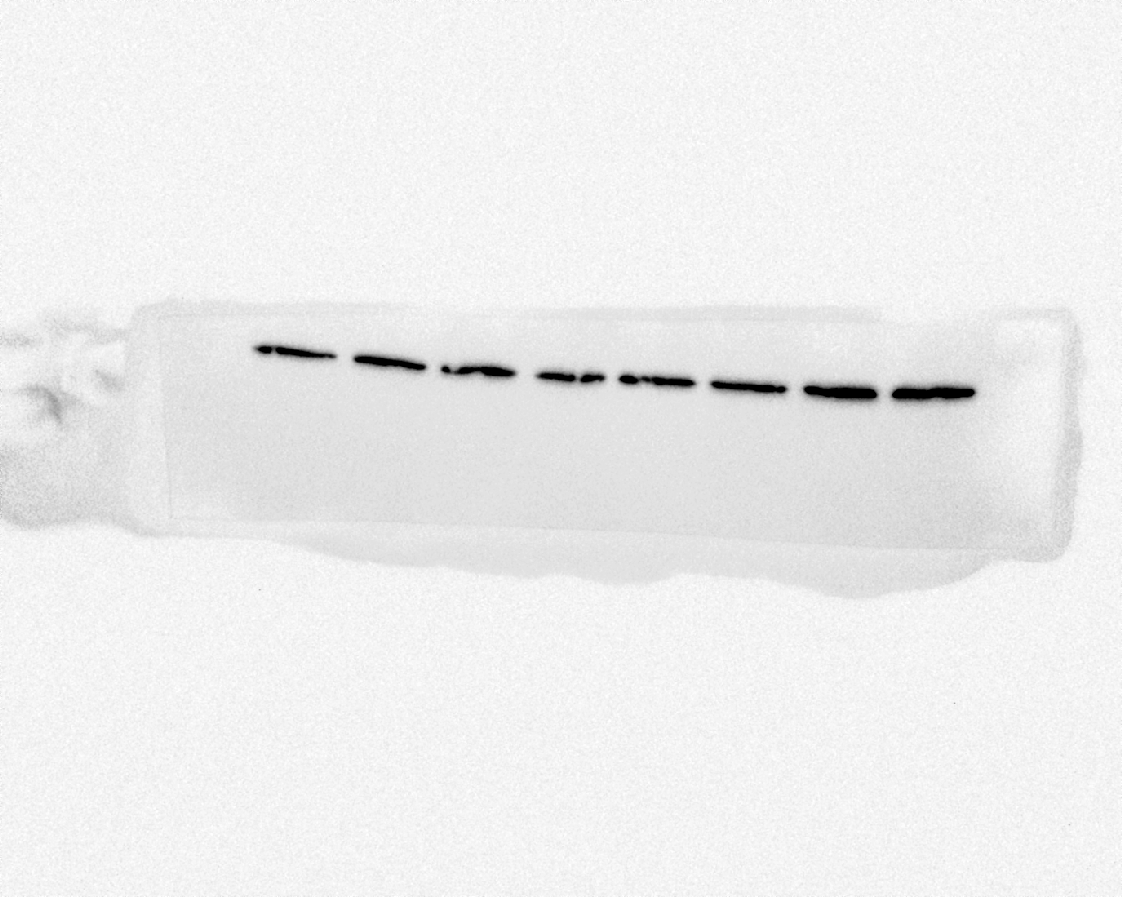

### Figure S6F_D-NPCs_RFX3_WT_RFX3-HOM.tif

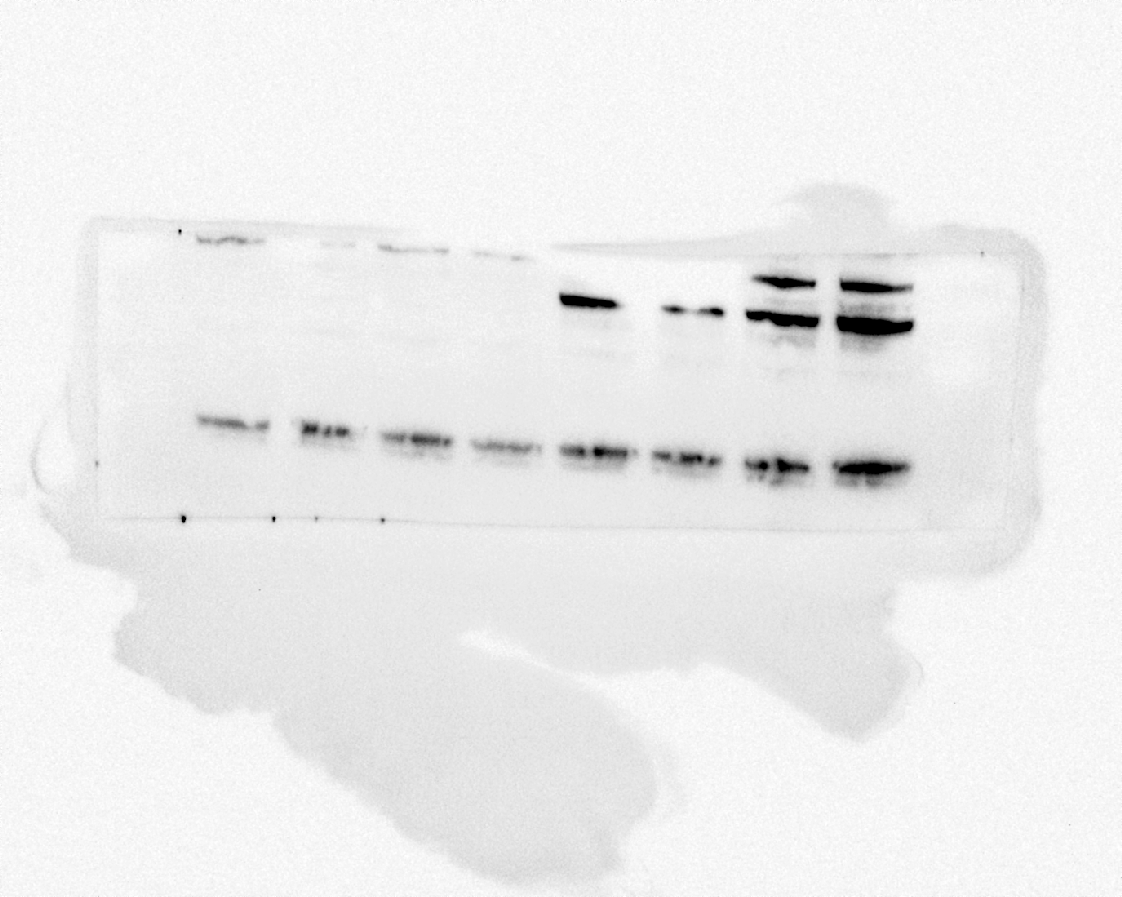

### Figure S9C_D-NPCs_RFX4_GAPDH_WT_R79C-HET_R79C-HOM.jpg

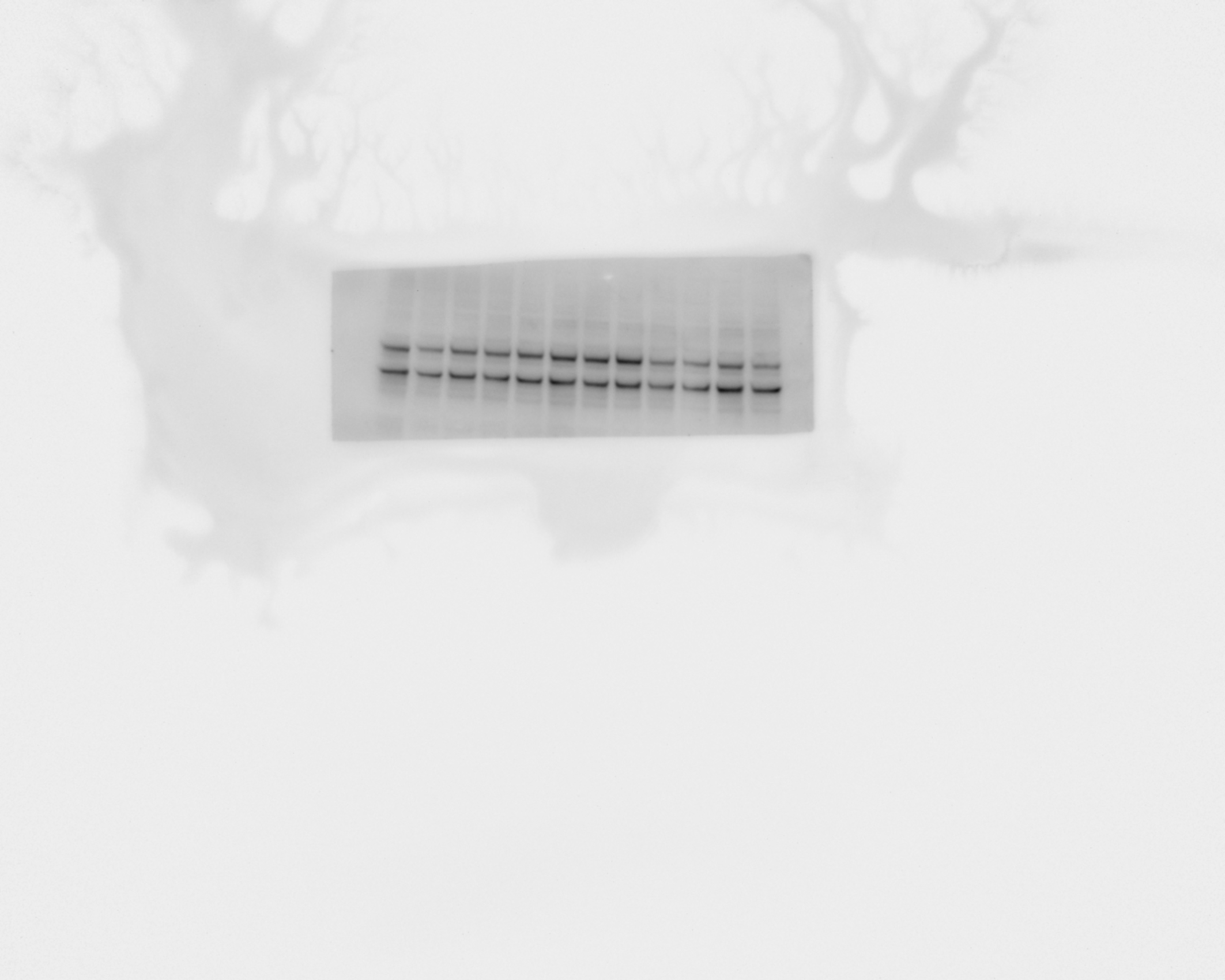

### Figure S9C_D-NPCs_RFX4_WT_R79C-HET_R79C-HOM.jpg

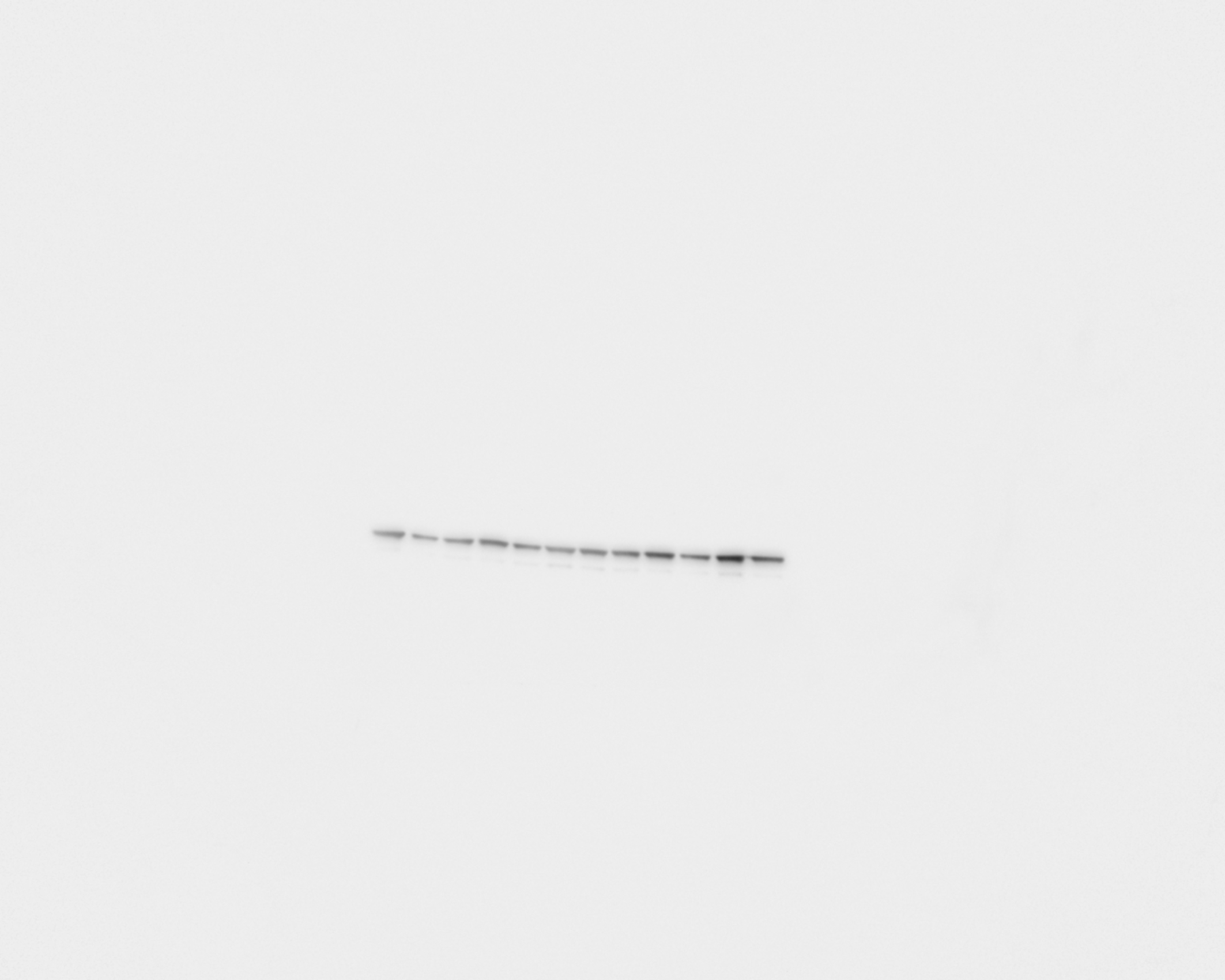
